# Generating antimicrobial peptides via genomic transfer learning

**DOI:** 10.64898/2026.06.16.732639

**Authors:** Lucrezia Polloni, Igor Gonteri, Kamil D. Bieniasz, Jarvist Moore Frost

**Affiliations:** Department of Chemistry, Imperial College London, Exhibition Road, London SW7 2AZ, UK; Institut des sciences et ingénierie chimiques (ISIC), Ecole polytechnique fédérale de Lausanne (EPFL), Station Z, CH-1015 Lausanne; Department of Physics, Imperial College London, Exhibition Road, London SW7 2AZ, UK

## Abstract

We present a generative machine learning pipeline for the design of linear antimicrobial peptides (AMPs). To extend diversity beyond synthetically validated peptide datasets (∼7,000 entries), we apply transfer learning by training a Generative Pre-trained Transformer (GPT) on the genomically derived AMPSphere dataset (∼863,000 entries), before fine-tuning on the Database of Antimicrobial Activity and Structure of Peptides (DBAASP). We assess the filtered sequences with a committee of Minimum Inhibitory Concentration (MIC) predictive models built with a Bi-LSTM architecture, and ESM-2 and QSAR feature vectors.

The fine-tuned GPT model produced a 28% reduction in test loss compared to training on DBAASP alone, and generates peptides that are simultaneously more novel and more physicochemically plausible. Our top-ranked candidates are predicted to possess antimicrobial activity comparable to polymyxin B.

We anticipate this transfer-learning approach is broadly applicable for leveraging massive, unlabelled genomic datasets to enrich targeted peptide discovery. Our identified sequences have been submitted to the 2027 AMP Challenge^1^ (team name VINCI) for experimental validation, and the complete codebase and workflow are open source^2^.

## I. INTRODUCTION

Antimicrobial resistance is projected to cause ten million deaths annually by 2050, making it a defining health threat of the 21st century^3^. The World Health Organization (WHO) Bacterial Priority Pathogen List (BPPL) identifies Gram-negative (GN) bacteria as among the greatest threats, owing to their intrinsic resistance to last-resort antibiotics and rapid acquisition of novel resistance mechanisms^4^. Yet the conventional commercial drug-development pipeline has stalled^5,6^. Overcoming this stalled pipeline requires innovative, open-source approaches to drug design^7^.

Antimicrobial peptides (AMPs) are one promising drug class. These short, cationic sequences of typically 8–50 residues form a core component of innate immunity and exploit a physical asymmetry between cell envelopes^8^: cationic residues (Arg, Lys) bind the anionic bacterial surface, while bulky hydrophobic residues (Trp, Phe, Ile, Leu) drive insertion into the acyl-chain region, permeabilising the membrane via carpet-like detergent disruption or, for longer sequences, toroidal- or barrel-stave-type pore assembly^8^. Because they target a conserved physicochemical feature rather than a single protein, AMPs tend to be broad spectrum and slow to provoke resistance^9^. Despite these advantages, few have reached the clinic (colistin^10^, polymyxin B^11^, and daptomycin^12^ being notable examples, all originating in natural products). Rational drug design is limited by a lack of a target, and the combinatorial sequence space is too vast to screen computationally, let alone experimentally.

Considerable recent efforts have built open databases of antimicrobial peptides and experimentally validated activities^13,14^. Deep generative models trained on these synthetic databases have accelerated computational design of peptides (see Section S1 of the Supplementary Information for further details), but these models are constrained by data sparsity. Our own experience with building a baseline LSTM based model (see Section S2 of the Supplementary Information) demonstrated that the high scorers from these models interpolate within the established data, providing limited chemical novelty. Conversely, foundational protein models trained on generic sequence data introduce a severe domain mismatch: the sequence grammar of large structural proteins is not matched to the short, and biologically unusual, AMP sequences required for membrane disruption.

We hypothesised that pre-training on these larger unlabelled (or semi-labelled) genomic based data, and then fine-tuning on the much sparser experimental datasets would bridge this gap. By constructing a suitably expressive and fast-training model (a generative pre-trained transformer), we are able to pre-train on the large AMP-Sphere dataset^15^ (863, 498 non-redundant prokaryotic peptides identified as being AMP-like), and then fine-tune on the Database of Antimicrobial Activity and Structure of Peptides (DBAASP)^13^, producing a model that extrapolates beyond the smaller database into a physicochemically plausible chemical space with strong predicted activity.

## II. METHODS

A visual summary of our workflow is shown in Figure 1.

**FIG. 1.**
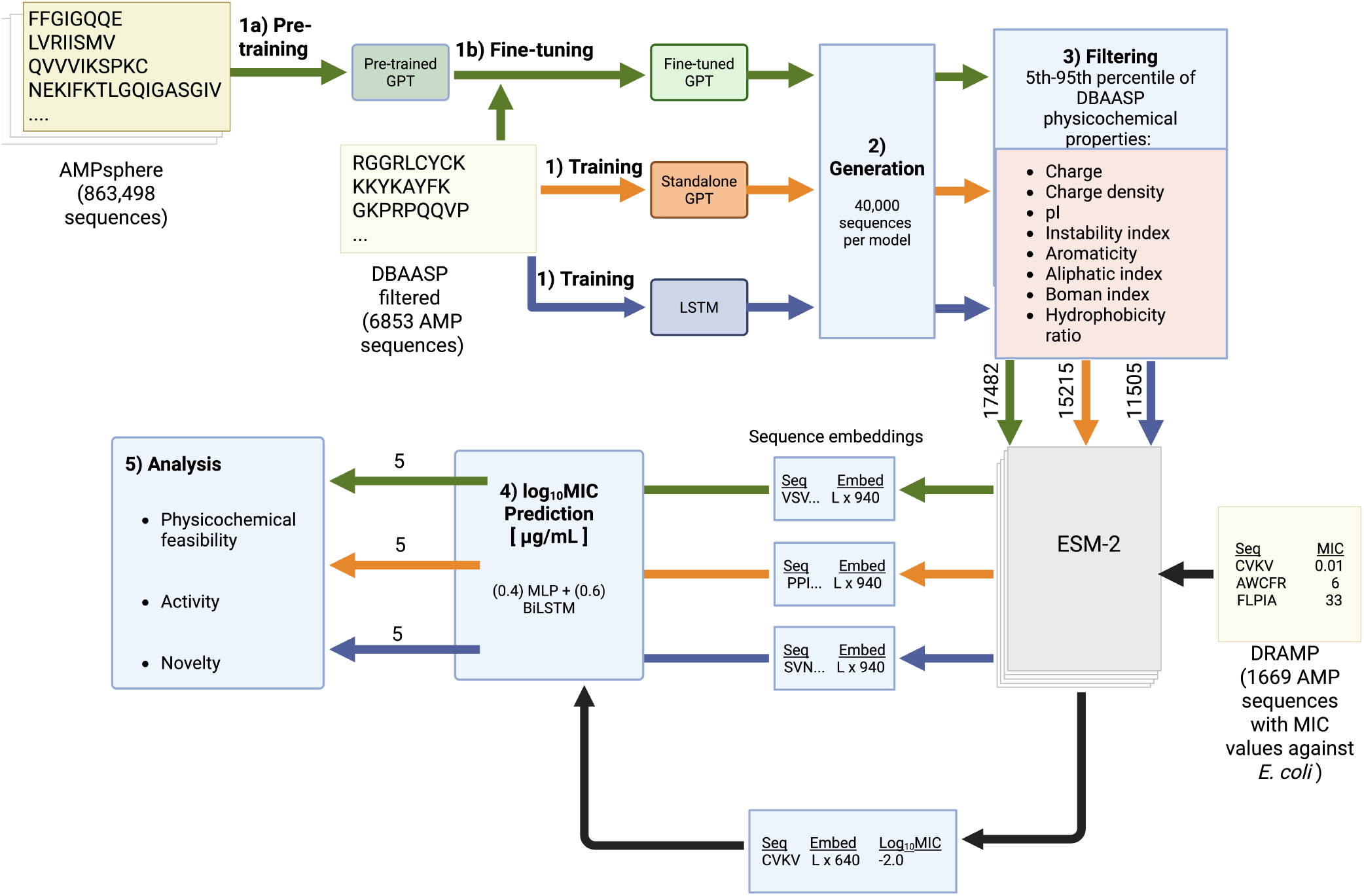
Overall workflow of the study comprising five key parts: (1) Generative model training; (2) Generation of 40,000 peptides by each trained model; (3) Physicochemical filtering of the generated peptides using 5th-95th percentile of DBAASP as boundaries for each property; (4) Prediction of log_10_(MIC) [µg mL^−1^] using the ensemble predictor; (5) UCB-based ranking of peptides and analysis of the top five candidates per model

### A. MIC Predictor

#### 1. Datasets

Training MIC data was sourced from DRAMP, which provides curated activity-labelled subsets with explicit organism-level annotations^14^. All values were converted to µg mL^−1^ using the calculated peptide molecular weight. If multiple values existed for the same peptide, the geometric mean was calculated. All MIC values were log_10_-transformed prior to modelling to compress the MIC range and reduce the influence of extremes. The dataset was filtered to retain only peptides with activity against Gram-negative (GN) bacteria; prioritised as they constitute the majority of WHO critical priority pathogens and present a greater therapeutic challenge due to their outer membrane permeability barrier^3,4^. Subsequently, sequences containing lowercase letters, non-canonical amino acids, special characters and duplicates were removed, yielding 1,793 peptides. These were used for initial hyperparameter optimisation across all eight MIC prediction models.

To improve internal consistency and reduce biological and experimental heterogeneity, the GN dataset was filtered to retain only *E. coli* activity data. This resulted in a curated dataset of 1,669 peptides. Final model training and evaluation were conducted exclusively on this *E. coli*-specific dataset.

#### 2. Sequence Representation

Three featurisation strategies were tested, each capturing different aspects of sequence information. First, raw sequence tokenisation mapped each amino acid to an integer index representing the 20 canonical amino acids plus a padding token (`<PAD>`) and unknown residues (`<UNK>`), giving a vocabulary size of 22. This strategy captured only amino acid identity and positional information and was chosen as the performance baseline: if richer representations (ESM-2 embeddings, QSAR descriptors) did not outperform this, augmentation may not be justified.

To augment the data, Quantitative Structure-Activity Relationship (QSAR) descriptors were calculated using the PyPro library, yielding a 267-dimensional feature vector composed of three descriptor families: the amino acid composition (AAC), Composition, Transition and Distribution (CTD) and Quasi-sequence order (QSO)^16^. Critically, for peptides composed of canonical amino acids, QSAR descriptors are grounded in experimentally established physicochemical properties that govern AMP-membrane interactions, making them interpretable and domain-specific. The final approach involved leveraging the Evolutionary Scale Modeling 2 (ESM-2) model. ESM-2 is a transformer-based encoder trained on hundreds of millions of protein sequences^17^. The variant used here (150M parameters) produces a 640-dimensional embedding for each residue, implicitly encoding complex, non-linear relationships related to protein structure and function, that are not captured by either tokenisation or QSAR descriptors alone.

#### 3. Model Architectures

Eight candidate architectures were evaluated across two model families (Table S2), differing in whether residues were modelled sequentially or independently. All models were implemented in PyTorch^18^ and produced a single scalar log_10_(MIC) [µg mL^−1^] output.

##### Multilayer Perceptron (MLP)

MLP variants operated on fixed-length inputs and served as a performance baseline for global-context-only prediction: if richer sequential representations did not outperform a pooled summary, the added complexity would not be justified. MLP (ESM+QSAR) concatenated mean-pooled ESM-2 embeddings with QSAR descriptors; MLP (ESM-only) used embeddings alone. Each input vector was passed through fully connected layers with GELU activations^19^ and dropout, followed by a linear output layer. Full architectural details are provided in Table S11.

##### Bidirectional Long Short-Term Memory (BiLSTM) Model

BiLSTM variants processed variable-length inputs sequentially, testing whether retaining residue-level order and local contextual structure improves MIC prediction beyond what is available from a pooled global summary. In a BiLSTM, sequences are simultaneously processed both forwards and backwards, improving global predictors. Forward and backward hidden states were concatenated at each position and passed through an MLP head to produce the log_10_(MIC) [µg mL^−1^] prediction^20^. Full architectural details and the LSTM equations are provided in Table S12 and Section S3 A.

#### 4. Training

The dataset was encoded and partitioned into training, validation, and test sets (70/15/15) using a fixed random seed. Before model training, continuous input features were normalised to ensure comparable feature scales. ESM-2 embeddings and QSAR descriptor vectors were standardised using scalers fitted only on the training split. The fitted scalers were then applied to the full dataset, including validation and test samples, ensuring that no information from held-out data influenced the normalisation parameters. Training was performed using mini-batch gradient descent for faster and more efficient training^21^. All models were trained to minimise the mean squared error (MSE) between predicted and observed target variable (log_10_(MIC) [µg mL^−1^]) over the training set, via backpropagation and the AdamW optimiser^22^.

The small dataset size necessitated the use of several regularisation and optimisation techniques to improve generalisation: a plateau learning rate scheduler and dropout were applied throughout^23^. Early stopping (patience = 5, minimum improvement threshold = 0.001) was monitored on validation loss; training was extended to three times the early-stopping epoch to ensure stable convergence and avoid premature termination due to validation noise. Sensitivity analyses confirmed that Huber loss (*δ* = 0.5, 1.0) did not consistently improve rank-based performance over MSE (Table S14). Hence, MSE was retained given its stronger penalisation of large deviations, which is directly relevant to identifying highly active peptides at the extremes of the MIC distribution. Weight decay of 1 × 10^−4^ was selected via systematic sweeps (Table S15), and controlled extension of training beyond early stopping yielded modest but consistent performance improvements (Figure S12, Table S16). Hyperparameter grid searches were conducted on the GN dataset; the top-five configurations per architecture were retrained on the *E. coli* subset (Figures S4–S11; Tables S3–S10).

#### 5. Evaluation

To account for the complexities and inherent noise of biological data, model performance was evaluated using a combination of standard regression metrics—including coefficient of determination (*R*^2^), root mean squared error (RMSE), mean absolute error (MAE), Spearman’s rank correlation coefficient (*ρ*)—alongside drug-discovery-oriented metrics—including Boltzmann-Enhanced Discrimination of Receiver Operating Characteristic (BEDROC Eq. (S6), *α* = 20) and Enrichment Factor at 10% (*EF*@10% Eq. (**??**))^24^. For the latter two metrics, “potent” sequences were defined as those within the lowest 10% of MIC values in the DRAMP training set. Model selection was guided by a combined score, EF@10% + 2*ρ*, where *ρ* is the Spearman rank correlation. The score was intended as a practical screening heuristic. *EF*@10% was incorporated to reflect the central objective of virtual screening: enrichment of active peptides among the top-ranked candidates. However, *EF*@10% can be sensitive to fluctuations within a small top-ranked subset which justified the additional inclusion of Spearman *ρ*; the larger weighting applied rewarded models that also maintained consistent global rank ordering across the test set. Therefore, the selected model balanced early enrichment with broader ranking robustness.

A configuration was accepted only if it achieved consistently high scores across both GN and *E. coli* test sets, further confirmed by manual inspection of training and validation loss curves. Particular attention was paid to stability and divergence between training and validation loss, which would indicate overfitting and reduced generalisation performance. The selected configuration was repeated with three different random seeds on the *E. coli* dataset to evaluate model stability.

#### 6. Ensemble Construction

Following individual model evaluation, three ensemble strategies were assessed: (i) equal-weight averaging of all model predictions; (ii) pairwise combinations with weights optimised by non-negative least squares (NNLS); and (iii) a full NNLS-weighted ensemble across all eight models. NNLS minimises 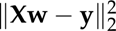 subject to **w** ≥ 2 0, preventing negative contributions from correlated predictors^25^. Weights were normalised to sum to unity. All ensemble variants were evaluated on the *E. coli* validation and held-out test sets using the same six metrics and combined score. The final MIC predictor was selected to balance predictive performance and reproducibility across seeds.

### B. Generative Model

#### 1. Datasets

DBAASP was filtered to peptides with unmodified termini, canonical amino acids, unique sequences, and length *<* 30 residues, yielding 6,853 sequences^13^. Length restriction and exclusion of non-canonical modifications ensured compatibility with standard solid-phase peptide synthesis (SPPS) workflows^8,26^. The AMPSphere corpus, comprising 863,498 distinct prokaryotic peptide candidates identified by machine learning-based genome mining of 63,410 metagenomes and 87,920 bacterial and archaeal genomes, was used for GPT pre-training^15^. subject mining of 63,410 metagenomes and 87,920 bacterial and archaeal genomes, was used for GPT pre-training^15^.

#### 2. Shared Training Implementation

All generative models were implemented in Julia using the Flux.jl deep learning framework and trained on GPU^27^. Peptide sequences were tokenised over the 20 canonical amino acids plus a boundary/padding token (’_’), mapped to integer indices. A single continuous corpus was constructed by concatenating all sequences separated by the ‘_’ token, providing an explicit boundary signal that allows the model to learn that adjacent peptides are independent. Targets were one-hot encoded for compatibility with the loss function (Eq. (**??**)). The corpus was segmented into fixed-length overlapping context windows of 32 tokens (standalone GPT and LSTM) or 128 tokens (AMPSphere-pretrained GPT), forming (*X*, *Y*) next-token prediction pairs. Blocks were split sequentially: 90% training, 10% test. All models were trained using the Adam optimiser to minimise the cross-entropy loss (Eq. (**??**)) over next-token predictions^28^, and also calculated the per-character perplexity (Perplexity = exp(*J*)). Lower perplexity indicates greater model confidence in the correct next token; a perplexity equal to the vocabulary size, here 21, corresponds to random uniform prediction^20^.

#### 3. Architecture Comparison

To debug and optimise performance of the models, we made extensive use of standard natural language dataset, the complete works of Shakespeare^29^. Transformed to uppercase and dropping all punctuation (therefore 26 characters), it was a similar size and structure, to the antimicrobial peptide datasets. The outputs from this corpus were deeply interpretable; numerically subtle differences in loss between different architectures and hyperparameters led to massive variations in the perceived quality of the nonsense words generated.

With the goal of controlled comparison between a recurrent baseline and a GPT-style autoregressive model, rather than independent performance maximisation, all hyperparameters were matched across standalone model architectures, following established comparative practice in sequence modelling^30–32^—with one intentional exception. A layer sweep was carried out on the LSTM as recurrent depth is architecture-specific and has no direct analogue in transformer architectures (Figure S24). The 2-layer LSTM achieved near-identical test loss to the 3-layer variant with a more favourable train–test gap, and was therefore selected to minimise overfitting risk without sacrificing representational capacity^33,34^. All remaining hyperparameters were kept aligned, ensuring that observed performance differences reflect architectural bias rather than unequal training conditions.

#### 4. Standalone Generative Pre-trained Transformer (GPT) Model

We wrote a decoder-only autoregressive transformer following the nanoGPT architecture^35^. Input tokens were represented by learned embedding of dimension *n*_embed_, summed with learned positional embeddings over a context window of 32 tokens. The combined representation was passed through *n*_layer_ transformer blocks, each comprising a pre-layer-normalised multi-head self-attention module with causal mask followed by a position-wise feedforward network. The feedforward hidden dimension *n*_hidden_ was set to 4× *n*_embed_, matching the standard transformer expansion ratio, and used a GELU activation function. The number of attention heads *n*_heads_ was constrained to divide *n*_embed_ evenly, resulting per-head projections of dimensions *n*_embed_ / *n*_hidden_. A final layer normalisation and linear projection mapped hidden states to vocabulary logits. Dropout was applied throughout to mitigate overfitting on the small DBAASP dataset (Figure S28). Hyperparameters were selected by manual search on the Shakespeare corpus (Figure S21) followed by a learning rate sweep on DBAASP (Figure S29).

#### 5. Long Short Term Memory (LSTM) Model

A stacked character-level LSTM was implemented with the same embedding dimension, hidden size, sequence length, batch size, and learning rate as the GPT. Token embeddings were passed sequentially through LSTM layers—selected by layer sweep over one to four layers (Figure S24)—followed by a linear output projection to the vocabulary. Hidden size was fixed at 4× the embedding dimension, consistent with the feedforward expansion ratio. Inter-layer dropout was applied.

#### 6. AMPSphere-pre-trained GPT

A two-stage approach was employed. In Stage 1, a scaled-up GPT (increased *n*_embed_, *n*_hidden_ and context window of 128) was pre-trained on the full AMPSphere corpus to learn the statistical grammar of antimicrobial-like sequences without activity supervision. Early stopping (patience = 15) was applied, with a higher patience than the standalone models to accommodate the larger model and more diverse dataset. In Stage 2, the pretrained weights were fine-tuned on DBAASP with all parameters trainable. Fine-tuning hyperparameters (learning rate, dropout, batch size) were selected by grid search (27 configurations) minimising held-out test loss. The lowest learning rate among the top-five configurations was selected to reduce the risk of catastrophic forgetting. Model checkpoints were saved at epochs 3, 5, 7, and 10; the optimal checkpoint was selected by retention rate after physicochemical filtering (Section II C) rather than held-out loss, which was comparable across checkpoints. This design choice prioritised retention of pre-trained structural diversity over DBAASP-specific adaptation.

### C. Sequence Generation and Physicochemical Filtering

Each trained model generated 40,000 peptide sequences via autoregressive sampling, initialised with the boundary token ‘_’. At each step, logits were temperature-scaled (T = 1) and converted to probabilities via softmax, from which the next token was sampled stochastically. Generation continued until the boundary token was predicted or sequence length reached 30 residues.

Generated sequences were filtered against DBAASP-derived physicochemical bounds: for conservative filtering the 5th–95th percentile range across ten properties (length, molecular weight, charge, charge density, isoelectric point, instability index, aromaticity, aliphatic index, Boman index, hydrophobic ratio) calculated using modlAMP^36^. Sequences failing any bound were discarded. Sequences present in the DBAASP training set were additionally removed. Retention rate was computed for each model as a proxy for physicochemical plausibility:

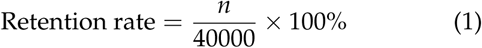

where n is the number of peptides remaining post-filtering.

Embedding-space coverage was assessed by UMAP applied to L2-normalised concatenated ESM-2 + QSAR feature vectors (907 dimensions), reduced to 50 principal components (retaining 96.2% of variance) prior to projection (*n*_neighbors_ = 15, *min*_dist_ = 0.15)^37^. Density structure in the projected space was characterised using HDBSCAN (*min*_cluster_size_= 200, *min*_n_samples_ = 10, Euclidean metric)^38^. Distribution distance from the reference peptide sets was quantified by calculating the Wasserstein distance^39^ between the candidate peptide and all peptides in the reference.

### D. Candidate Prioritisation

The selected ensemble MIC predictor was applied to all filtered sequences using three independent random seeds. For each peptide, the mean (*µ*) and standard deviation (*σ*) of predicted log_10_(MIC) [µg mL^−1^] across seeds were recorded, with *σ* treated as a proxy for prediction uncertainty. Candidates were ranked by their Upper Confidence Bound (UCB) UCB = *µ* + 0.5*σ*. Because a lower MIC denotes higher activity, adding the variance penalty yields an estimate of the peptide’s *worst-case* MIC. A relatively conservative weight of *w* = 0.5 was chosen so that inter-seed variability contributed to ranking without dominating the mean potency signal.

The top five candidates per generative model were evaluated for physicochemical plausibility, predicted antimicrobial potency, and sequence novelty.

Feasibility was assessed by calculating key physicochemical properties (Section II C) and comparing them with literature-derived design thresholds (Table S23) to determine whether the peptides were consistent with known AMP design principles. Potency was assessed using the predicted mean log_10_(MIC), with lower values indicating greater antimicrobial activity. Finally, novelty was quantified as the minimum weighted Levenshtein distance (Eq. (2); substitution cost = 1, insertion/deletion cost = 2) from each generated peptide to all sequences in DBAASP and AMPSphere separately^40^, serving as a direct sequence-level proxy for chemical space distance.

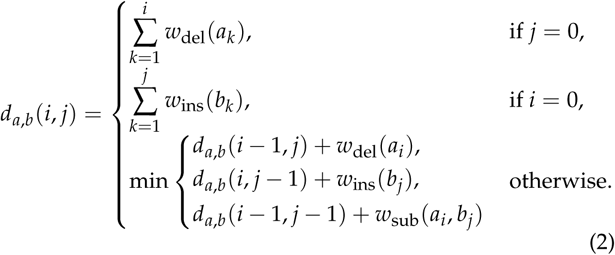

where:

- *w*_del_(*a_k_*) is the cost of deleting character *a_k_* at position *k* in string *a*;
- *w*_ins_(*b_k_*) is the cost of inserting character *b_k_* at position *k* in string *b*;
- *w*_sub_(*a_i_*, *b_j_*) is the cost of substituting *a_i_* with *b_j_*, which is 0 if *a_i_* = *b_j_*.

Setting higher insertion and deletion costs than substitution cost reflected the greater functional disruption caused by length-altering operations relative to single residue substitutions, which are more commonly tolerated in peptides. Greater distance to DBAASP indicates lower sequence-level similarity to known AMPs while proximity to AMPSphere reflects alignment with broader, natural AMP-like sequence space.

### E. Reproducibility and Synthesisability Filtering

Reproducibility was enforced throughout the computational pipeline. During ESM-2 embedding extraction, torch.backends.cudnn.deterministic = True and torch.backends.cudnn.benchmark = False were set to reduce nondeterministic CUDA kernel selection. Model training used fixed random seeds for PyTorch and NumPy, and dataset partitioning was performed with a seeded torch.Generator to ensure identical train/validation/test splits across runs^41^. To ensure consistent inference-time preprocessing, normalisation scaler parameters (feature means and scales) were serialised in the model checkpoint together with the full argument dictionary and reloaded during prediction. Reproducibility of the final candidate-selection outputs was verified using a dedicated utility, compare_top100.py, which checks array shape, column order, sequence-set identity, row order, and pairwise numerical agreement within an absolute tolerance of 1 × 10^−6^. Candidate peptides were filtered for synthetic accessibility. Each sequence was assigned a score across six physicochemical dimensions targeting a distinct, mechanistically grounded failure mode in SPPS or antimicrobial activity. Values ranged from 0 − 100 with higher values indicating greater synthetic tractability. The dimensions of the synthesisability score and their assigned weights were: instability index (17%), as defined by Guruprasad et al., penalising values above 40 indicative of in vivo instability^42^; sequence length (16%), favouring the 8-20 residue window established for solid-phase peptide synthesis and flagging sequences exceeding 30 residues for elevated truncation risk^43^; net charge at pH 7.4 (17%), requiring a minimum of +1 with the validated antimicrobial activity range of +2 to +9^8^; mean Kyte–Doolittle hydrophobicity (17%)^44^, penalising values above 0.5 associated with on-resin *β*-aggregation and reduced crude purity^45,46^; sequence motif penalties (17%), applying hard deductions for problematic features including diproline motifs, Asp-Pro junctions, multiple cysteine or methionine residues, N-terminal glutamine or glutamate, and hydrophobic runs of five or more residues^45^; and Eisenberg hydrophobic moment (16%)^47^, with the empirically-derived hydrophobic moment defined as 0.2–0.5^8,48^. Based on the number and severity of threshold breaches, each peptide was assigned one of three flags: PASS (no breaches), WARN (exactly one minor breach, comprising instability index exceedance, low charge, or insufficient length), or FAIL (any critical breach, including excessive hydrophobicity, length above 30 residues, or a flagged sequence motif, or any combination of two or more breaches of any kind). Peptides receiving a FAIL classification were excluded from further consideration. Surviving PASS and, optionally, WARN peptides were subsequently ranked by ascending ensemble-mean log_10_(MIC) [µg mL^−1^], and the top 100 most potent candidates were retained and written to both a CSV file and a FASTA file for downstream analysis.

## III. RESULTS AND DISCUSSION

### A. Minimum Inhibitory Concentration (MIC) Predictor: Benchmarking, Calibration, and Architecture Selection

The details of our full grid search and hyperparameter sweep are given in the Supplementary Information (section **??**).

Our final MIC predictor model was a committee of 60% a model trained with ESM and QSAR features only, and 40% the Bi-LSTM sequence level predictor.

### B. Generative Model Training: Architecture Comparison and Transfer Learning

The Shakespeare corpus was used extensively when developing and refining the generative models, after having noted that broken into words it was of a similar size and diversity as the synthetic peptide benchmarks, as we could intuitively and directly see improvements in the quality of the generated words. Full details of these early developments are in the Supplementary Information (section S4 A).

Having established our GPT as a superior model, we then applied it to transfer learning on AMPSphere. To avoid overfitting on the larger corpus (≈863,000 vs. ≈6,800 sequences in DBAASP), architectural scaling was necessary (Figure S33). Fine-tuning on DBAASP yielded a 28% reduction in best test loss relative to the standalone GPT trained on DBAASP alone (Figure S32). This aligns with transfer learning theory: AMPSphere pretraining provides a strong initialisation that captures general AMP-like sequence structure, enabling more efficient and better generalised fine-tuning on the smaller, experimentally validated dataset. After physicochemical filtering, the fine-tuned GPT achieved the highest retention rate, followed by the standalone GPT and LSTM (Table S22). The monotonically decreasing retention across fine-tuning epochs indicated that excessive fine-tuning erodes structural priors acquired during AMPSphere pretraining; the 3-epoch checkpoint was therefore used for all subsequent analyses. Using a lower learning rate or freezing parts of the network could be considered for future work. Overall, the GPT-architectures outperformed the LSTM baseline, implying that the ability of GPTs to model longer-range dependencies appears to produce more chemically coherent sequences even prior to any activity assessment.

### C. Characterisation of Generated Peptide Sets: Physicochemical Space, Activity, and Novelty

#### 1. Embedding-Space Coverage and Physicochemical Properties

UMAP on the full 907-dimensional embeddings established filtering as a technique to tighten the distribution, bringing models in closer alignment with the reference AMP feature space rather than forming obvious outliers (Figure 2). Post-filtering GPT-based models span a broader region of the DBAASP/DRAMP embedding space than the LSTM, indicating superior physicochemical coverage of validated AMP-like space. The fine-tuned GPT achieved this despite generating more novel sequences (Figure 3) — demonstrating that metagenomic pretraining expands the reachable AMP-like manifold rather than interpolating DBAASP.

**FIG. 2.**
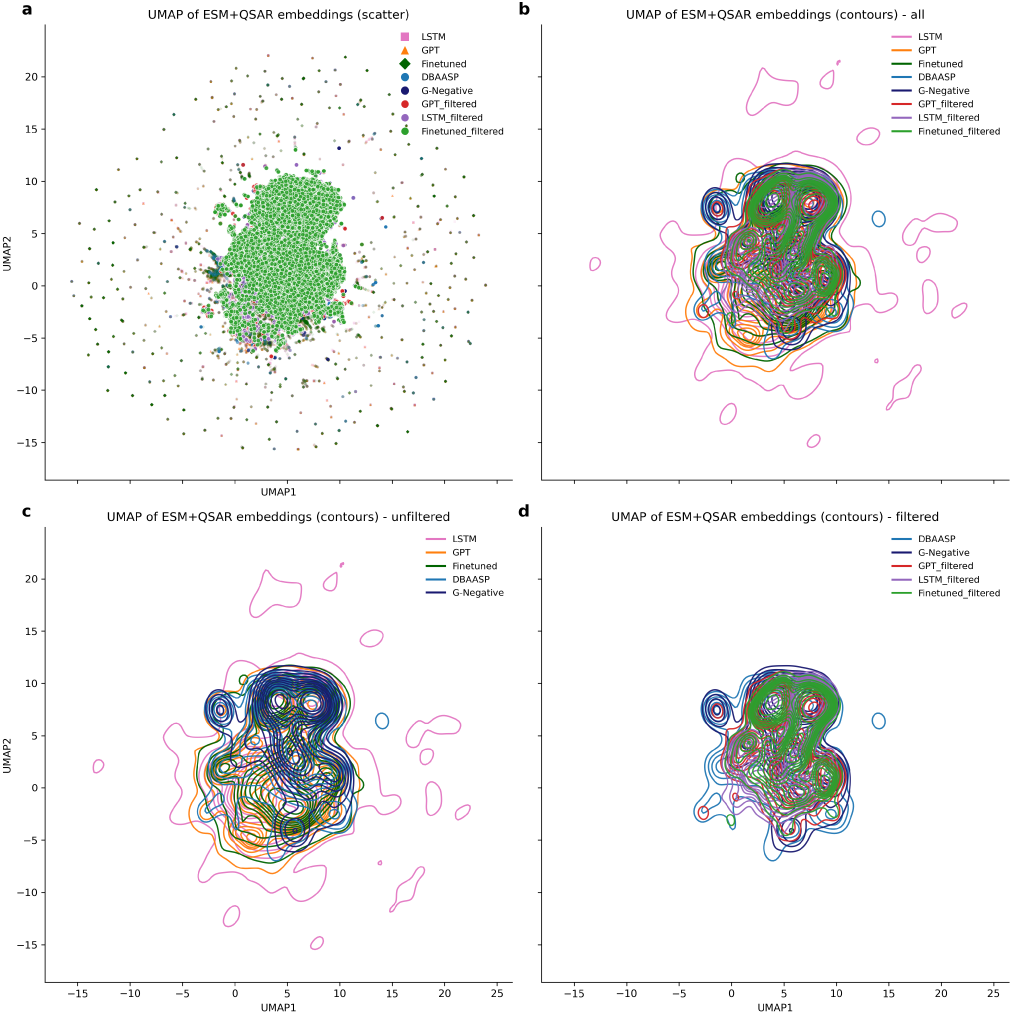
UMAP visualisation of ESM + QSAR embeddings across generated and reference antimicrobial peptide datasets.(a) Scatter plot of all groups, with raw generated datasets forming the background distribution and filtered subsets overlaid as larger markers. (b) Kernel density contours for all groups. (c) Contours for unfiltered generated datasets and reference peptide databases. (d) Contours for filtered generated datasets against reference peptide databases. Abbreviations: DBAASP = Database of Antimicrobial Activity and Structure of Peptides; G-Negative = experimentally validated Gram-negative-active peptides.

**FIG. 3.**
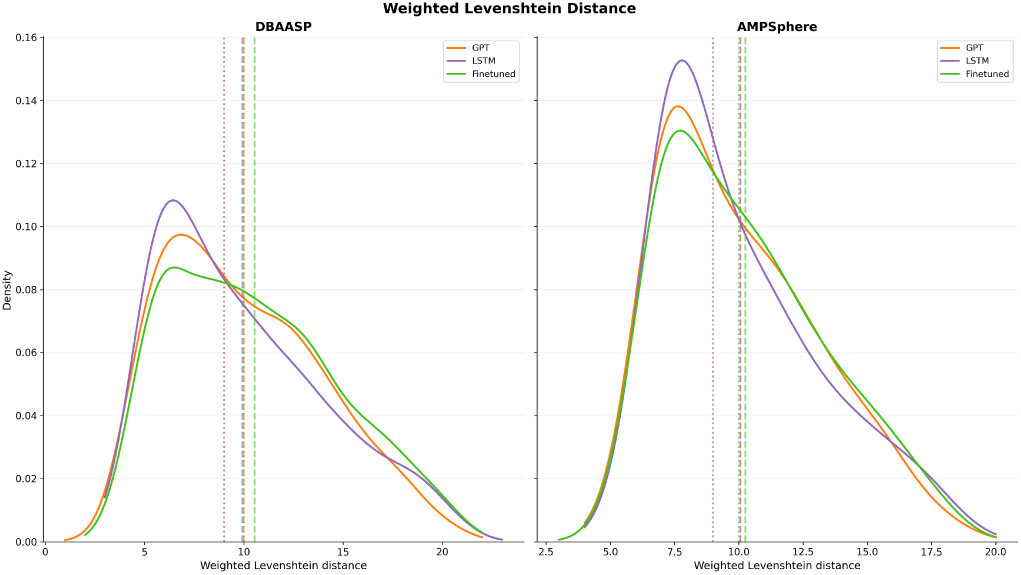
Distribution of weighted Levenshtein distances between filtered generated peptides and reference datasets (DBAASP, left; AMPSphere, right), with substitution cost = 1 and insertion/deletion cost = 2. Across models, mean distances (dotted lines) to DBAASP are 9.99 (GPT), 9.93 (LSTM), and 10.5 (fine-tuned), and to AMPSphere are 10.1 (GPT), 10.1 (LSTM), and 10.3 (fine-tuned). Median distances (vertical dashed lines) are consistent at 9.00 for GPT and LSTM and 10.00 for the fine-tuned model against both reference datasets. Lower distances indicate higher similarity to known sequences; higher values reflect increased novelty.

Given that UMAP provides only a global view of embedding-space similarity, individual physicochemical properties were examined next. Beginning with length (Figure S35), the DBAASP reference set exhibited a broad distribution with a higher central tendency compared to all generated datasets. The GPT-based models produced slightly shorter peptides, with the fine-tuned model more closely approximating the DBAASP distribution than the standalone GPT. In contrast, the LSTM generated the shortest sequences overall and showed a distribution shifted towards lower lengths. The limited ability of the LSTM to construct longer sequences was detected in the auxiliary analysis undertaken (Figure S25). Wasserstein distances from DBAASP were negligible for charge density, aromaticity, and hydrophobic ratio across all models, however, the largest deviations occurred in instability index and aliphatic index (Figure S36). Beyond the magnitude of divergence, the direction of the distributional shift provides insight into functional relevance (Figure S37). The GPT-based models shifted instability index towards lower values and aliphatic index towards higher values which are both associated with increased peptide stability. This effect was most pronounced in the fine-tuned model. Contrastingly, the LSTM shifted in the opposite direction, implying reduced stability. Clearly, the fine-tuned GPT independently learned to favour these stability-associated shifts without explicit supervision suggesting that AMPSphere pre-training encoded physically meaningful structural biases into the model’s sequence priors. This effect is reinforced in the top-candidate profiles examined in Section III D 1.

The distribution of these key physicochemical properties (Figure S37) belies the limitation of training solely on small synthetic databases, regardless of the power of the generative machine learning architecture. Both baseline LSTM and GPT models fail to capture the stability of the DBAASP reference database, when asked to generate novel sequences. The fine-tuned GPT spontaneously shifts the generated sequences towards higher stability and aliphatic indices, without directly conditioning on these requirements. The semi-supervised genomic pre-training has guided the generative model into a more plausible and biologically relevant part of chemical space.

### D. Top-Candidate Prioritisation

#### 1. Physicochemical Qualification and Predicted Potency

The top five candidates per model, selected by UCB ranking, were analysed in detail. As expected post-filtering, all fell within DBAASP 5th–95th percentile bounds across six AMP-relevant physicochemical descriptors (Figure 4). However, fine-tuned GPT candidates were more tightly distributed around literature defined AMP optima (Table S23). In particular, they displayed the lowest instability indices, highest aliphatic indices, and lowest Boman which is favourable for forming helical, amphipathic, membrane-disrupting peptides with relative protease resistance. Standalone GPT sequences showed greater structural diversity, with two peptides exceeding the Boman protein-binding threshold (>2.48 kcal/mol), suggesting possible non-membrane mechanisms. LSTM- and standalone GPT-generated peptides exhibited the weakest physicochemical profiles with respect to membrane-insertion and stability-related properties. Hydrophobicity is a critical determinant of antimicrobial activity, governing the ability of cationic AMPs to partition into the bacterial lipid bilayer and achieve the membrane association required for pore formation and disruption^8,46^. However, the majority of LSTM and GPT candidates presented low aliphatic indices (*<*100) and hydrophobic ratios (*<*0.4), indicating an insufficient hydrophobic character to drive stable membrane insertion. Furthermore, most LSTM peptides exceeded the instability index threshold of 40, above which sequences are predicted to have a shortened in vivo half-life owing to dipeptide-driven proteolytic susceptibility^42^. Collectively, these deficiencies suggest that both LSTM- and GPT-generated candidates are at elevated risk of reduced antimicrobial efficacy relative to fine-tuned model candidates, which presented more favourable profiles across these dimensions. That the fine-tuned model’s top-candidate profile emerged from UCB ranking without specific structural supervision suggests the model learned AMP structural grammar from sequence statistics alone.

**FIG. 4.**
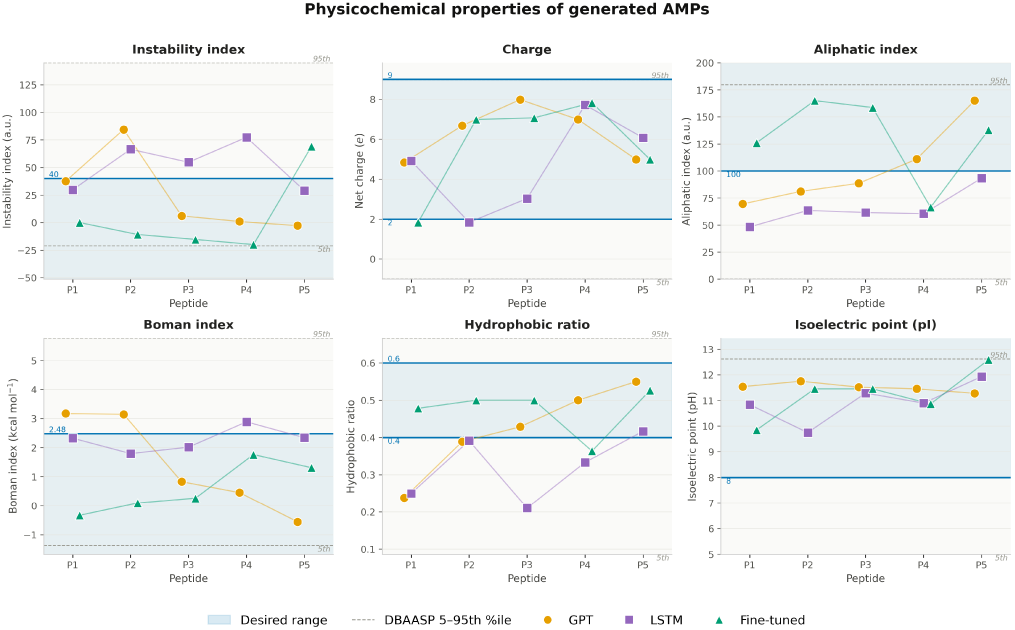
Top-candidate physicochemical properties. Six properties are shown: instability index, charge, aliphatic index, Boman index, hydrophobic ratio, and isoelectric point (pI). The top 5 candidates generated by the GPT- and LSTM- are shown in orange and purple, respectively; fine-tuned GPT peptides are shown in green. The peptide identifiers (P1–P5) correspond to the numbering provided in Figure S39, where the full sequences and descriptor values are reported. Blue shaded region marks established AMP design thresholds are summarised in Table S23. Dashed lines denote the DBAASP feasibility range (5th and 95th percentile, shaded).

The fine-tuned model produced the most potent top candidates, followed by the standalone GPT and the LSTM (Figure S42). The highest-priority candidate, VSVGVGVKVGVGVGVKVGVRESC, was predicted within the range of clinically significant activity against *E. coli*, comparable to that of Polymyxin B — a last-resort clinical agent^11^. Sequence distances to the nearest DBAASP neighbours were comparable and consistently ranked within the top 20% of predicted activity, confirming that UCB selection identifies sequences in high-activity regions of sequence space (Figure S42). Generated peptides, with one LSTM exception, were predicted to exhibit lower log_10_(MIC) than their closest database counterparts, indicating improved predicted potency while retaining sequence similarity to known AMPs. Discrepancies between predicted and experimental DBAASP MIC values are attributable to organism-specific variability, assay heterogeneity, or the lack of experimental testing on *E. coli* specifically. Hence, the DBAASP comparison should be interpreted as supportive evidence rather than definitive proof of efficacy.

#### 2. Sequence Novelty and Structural Origin

The novelty analysis revealed a clear dissimilarity between models. The standalone GPT and LSTM top candidates showed comparable Levenshtein distances to both DBAASP and AMPSphere databases, indicating generalised sequence space exploration without directional exploration. By contrast, fine-tuned candidates were notably closer to AMPSphere than DBAASP (Figure S44–S45), despite the fine-tuned model generating more globally divergent sequences overall (Section **??**). This is illustrated by two top candidates: VSVGVGVKVGVGVGVKVGVRESC showed Levenshtein distance d = 9 to its nearest AMPSphere match (AMP10.029_472, sharing GV/KV repeat motifs) versus d = 16 to its closest DBAASP match. Similarly, VQKVLKVHKVLKVHKVLKVGKV showed d = 8 to AMPSphere (AMP10.118_198), retaining comparable lysine enrichment, versus d = 15 to DBAASP. Therefore, fine-tuned candidates shared motif-level hallmarks with their AMPSphere neighbours while introducing sufficient divergence to constitute genuinely novel sequences.

The proximity to AMPSphere sequences correlated with more-active MIC, suggesting the genomic database contains highly active, uncharacterised, natural products (Figure S40). To determine whether specific sources were overrepresented, the biological origins of the closest-matching sequences to the fine-tuned candidates were examined. Substantial diversity was observed among the matched peptides, reinforcing the broad natural diversity of AMPs (Table S24), with no two candidates originating from the same family or source. Moreover, the wide range in gene counts suggests that the selected peptides captured both rare and recurrent AMP candidates, which may be useful for contrasting rare yet potentially novel sequences with more broadly distributed AMP-like motifs.

### E. Synthesisability Filtering and Final Candidate Library

To minimise sequence-level SPPS failure modes, synthesisability filtering retained only candidates passing all motif checks (Section II E). The final 100-peptide synthesisability-filtered library (Tables S25–S26) comprised sequences from all three generative models, with the standalone GPT contributing the largest share (67%), followed by the fine-tuned GPT (31%) and the LSTM (2%). This composition reflects the higher mean predicted activity of the standalone GPT across the full filtered set (Section **??**), resulting in a greater overall proportion of sequences meeting the activity and synthesisability thresholds for library inclusion.

Despite its smaller library share, the fine-tuned GPT contributed disproportionately among the most active candidates: it accounted for 20% of the top-10 sequences while constituting only 31% of the library, and its candidates spanned the full predicted activity range of the library. The single most active sequence in the synthesisability-filtered library, KFKVKAFKFKVGAGVFGVPVP, originated from the standalone GPT (log_10_(MIC) [µg mL^−1^] = −0.003). The LSTM contributed only two sequences, both ranking in the lower half of the library, consistent with its weaker generative performance observed throughout this study. Collectively, the synthesisability-filtered library confirms that the fine-tuned GPT offers a favourable balance of predicted potency and synthetic accessibility among the three generative models evaluated.

### F. Conclusion

The fine-tuned GPT model emerged as the most effective architecture for generating candidates with high predicted activity and plausible physicochemical profiles. Transfer learning from genomic data therefore seems a rich avenue for improving computational prediction of new peptides. The same techniques (and in fact, open-source codes and workflow) can be applied to other design problems in this space.

All trained-model checkpoints, training datasets, scaler parameters, and workflow code are publicly available at https://github.com/Frost-group/ ML-AMP-PIPELINE. The top candidates and full workflow have been submitted to the Szczurek Lab AMP Design Challenge^1^, which provides wet-lab MIC validation across 20 bacterial strains and HC50 measurements under a single standardised protocol. This offers an independent, blinded benchmark for the proposed pipeline that is not contingent on in-house synthesis capacity. Consequently, results from this challenge will constitute the primary experimental validation of the candidates identified here (Table S25–S26), and the open-access benchmarking dataset released under CC-BY 4.0 will enable direct comparison with other computational pipelines submitted to the challenge.

The principal bottleneck in the current workflow is the limited availability of large, standardised, and experimentally diverse peptide activity datasets. Existing AMP databases are small, noisy, and biased toward well-studied natural peptides, limiting generalisation and the construction of reliable negative datasets. Directing research effort toward coordinated functional screening of broader sequence libraries — including additional AMPSphere candidates — is therefore the most impactful near-term priority, as large-scale screening studies confirm this is feasible^49–51^. Architectural advances alone are unlikely to yield substantial improvements until this data bottleneck is resolved. Concurrently, continued advances in automated solid-phase peptide synthesis are reducing the cost and time required to experimentally validate computational candidates^52^, further lowering the barrier between *in silico* discovery and experimental confirmation.

The generation of standardised empirical datasets — encompassing both active and inactive sequences, as produced by the AMP Design Challenge^1^ — will be critical for the iterative refinement of MIC prediction models. Indeed, a key limitation encountered here when developing the MIC predictors was that inter-dataset experimental variability in measured MIC values exceeded the dynamic range of inter-peptide differences captured by the predictors. This hightlighted the need for consistent assay protocols. Transitioning to high-throughput synthesis and testing under standardised conditions, combined with open source development of machine-learning methods applied to shared datasets, will establish the feedback loop necessary for continued progress in computational AMP design.

## DATA AVAILABILITY

All trained model checkpoints, training datasets, scaler parameters, and workflow code are publicly available at https://github.com/Frost-group/ ML-AMP-PIPELINE under an open-source license.

The version submitted under team-name VINCI is archived on Zenodo^2^.

## IV. AUTHOR CONTRIBUTIONS

Contributor Role Taxonomy (CRediT). L.P.: Investigation (lead); Data curation (lead); Formal Analysis (lead); Methodology (equal); Validation (lead); Visualization (lead); Writing – original draft (lead); I.G.: Investigating (equal); Methodology (equal); Writing - original draft (supporting); K.B.: Investigating (equal); Methodology (equal); Writing - original draft (supporting); J.M.F.: Conceptualization (lead); Investigating (equal); Methodology (equal); Supervision (lead); Writing – original draft (equal); Writing – review and editing (lead).

## V. ACKNOWLEDGMENT

The authors are grateful for fruitful discussion with Anna Barnard, Andrew Edwards and Edward Douglas.

J.M.F. is supported by a Royal Society University Research Fellowship (URF-R1-191292), which also paid for consumables and synthesis. L.P., I.G. and K.B. undertook the research described herein in partial fulfilment of their degrees at the Department of Chemistry, Imperial College London.

## SUPPORTING INFORMATION

### S1. STATE OF THE ART

Computational approaches have accelerated progress in this field. Curated AMP databases, including the Database of Antimicrobial Activity and Structure of Peptides (DBAASP) and the Data Repository of Antimicrobial Peptides (DRAMP), have been compiled, providing structured access to experimentally validated AMPs and their minimum inhibitory concentrations (MICs)^S13,S14^. This enabled quantitative structure–activity relationship (QSAR) models, and, later on, deep learning classifiers for AMP identification: Veltri *et al.* (2018) demonstrated that convolutional and recurrent neural networks operating on numerical sequence encodings could discriminate AMPs from non-AMPs with high accuracy in AMPScanner, establishing a benchmark for sequence-based prediction^S53^. For regression-based activity prediction, Fjell *et al.* (2009) demonstrated that physicochemical descriptors derived from AMP databases could be used to quantitatively predict MIC from sequence alone, establishing the feasibility of QSAR-based potency estimation^S54^. Lately, Kolomeisky *et al.* (2023) showed that restricting feature selection to organism-specific descriptors substantially improves MIC prediction against *Escherichia coli* (*E. coli*), highlighting the importance of target-organism specialisation over broad-spectrum modelling^S55^.

Parallel advances in generative modelling followed, motivated by the need to explore sequence space rather than merely classify known peptides. Müller *et al.* (2018) and Nagarajan *et al.* (2018) independently applied recurrent neural networks (RNNs)—specifically long short-term memory (LSTM) architectures—to generate novel AMP-like sequences from curated databases^S56,S57^. Generative adversarial networks (GANs) and variational autoencoders (VAEs) subsequently extended this capability: Dean and Walper (2020) employed a VAE to design de novo peptides in a continuous latent space, while Szymczak *et al.*(2023) introduced HydrAMP, a conditional VAE (cVAE) that jointly models sequence and MIC, enabling guided generation of candidates with predicted activity against *E. coli*^S58,S59^. More recently, latent diffusion models operating on protein language model embeddings have shown promise for generating diverse AMP candidates with physicochemical properties consistent with the training corpus^S60,S61^.

The success of large language models (LLMs) in natural language processing has motivated their adaptation to protein sequences. A foundational insight is that, just as language derives meaning from the ordering and co-occurrence of words, protein sequences derive functional character from residue patterns, positional biases and long-range dependencies along the chain. These patterns arise from evolutionary constraints, which restrict viable residue combinations for stability and function, making them learnable from sequence data alone. Generative models such as ProtGPT2 (Ferruz *et al.*, 2022) and ProGen2 (Nijkamp *et al.*, 2023) pretrained autoregressive transformers on tens of millions of sequences from UniRef50 and UniRef90, learning broad protein sequence distributions and demonstrating the ability to generate sequences that fold into structurally plausible conformations^S62,S63^. AMP-Designer (Wang *et al.*, 2025) adopted this paradigm for AMP-specific design by pretraining a GPT model on UniProt and subsequently applying contrastive prompt tuning to bias generation towards antimicrobial activity, achieving a high experimental validation rate against Gram-negative bacteria^S64^. However, pretraining on generic protein databases introduces a fundamental domain mismatch: the vast majority of sequences in UniProt and UniRef correspond to structured proteins with lengths, amino acid compositions, and physicochemical properties that differ substantially from the short, cationic, amphipathic sequences that characterise AMPs. This may be an inefficient use of computational resources, as the model learns sequence patterns that are not directly relevant to AMP design. Critically, even after fine-tuning on curated AMP databases, models conditioned on generic protein priors have been observed to generate sequences that cluster within familiar regions of AMP sequence space, offering limited exploration of genuinely novel peptide chemistries. This represents a key unresolved challenge in computational AMP design: the tension between the scale required for effective language model pre-training and the domain specificity required for biologically plausible AMP generation.

To overcome this limitation, modern pipelines have shifted towards new input data for machine learning models: (meta)genomic datasets. These approaches identify small open reading frames (smORFs) that encode previously uncharacterised AMP-like sequences from microbiome data^S65,S66^. AMPSphere is a notable example, comprising 863, 498 non-redundant prokaryotic peptides uncovered through machine learning-based genome mining of 63,410 metagenomes and 87,920 microbial genomes^S15^. While experimentally unlabelled, their biological validity is supported by Santos-Júnior *et al.* (2024), who synthesised 100 AMPSphere candidates and found that 79 were active. Furthermore, the physicochemical properties of AMPSphere sequences—including mean positive charge (4.7 ± 2.6), high isoelectric point (10.9 ± 1.2), and amphiphilicity—are statistically consistent with experimentally validated AMPs, confirming that the database captures authentic AMP-like sequence grammar at ecologically relevant scale^S15^. This mirrors the established practice in protein language modelling of pretraining on large, computationally annotated databases (e.g. UniRef, BFD) to capture distributional sequence priors, before fine-tuning on smaller, experimentally validated datasets for task-specific adaptation^S62,S63^. Whereas traditional AMP databases are typically enriched for experimentally characterised peptides and variants of known natural products, AMPSphere extends coverage into less explored regions of natural peptide sequence space.

### S2. LSTM BASELINE

#### Peptide generation

Inspired by the NLP-to-peptide-sequence analogy established by Nagarajan^S57^, a character-level Long Short-Term Memory (LSTM) generative model was implemented in PyTorch. Each amino acid was embedded as a 128-dimensional continuous vector, and the embedded sequence was processed by a multi-layer LSTM followed by a fully connected output layer projecting onto a probability distribution over the 20 canonical amino acids plus a padding token. The model was trained on the DBAASP database (approximately 14,000 experimentally validated AMP sequences, truncated or padded to a fixed length of 15 residues), using cross-entropy loss and the Adam optimiser. Hyperparameters were tuned via random search with test accuracy as the selection objective. After training, new sequences were generated autoregressively: a three-residue seed sampled in proportion to the amino acid frequencies in DBAASP was fed to the model, which extended the sequence one residue at a time through temperature sampling until a termination token was predicted or the maximum length was reached. Duplicate sequences and those already present in the training set were discarded. From 40,000 generation calls, 39,846 unique novel sequences were obtained, demonstrating high generative novelty.

#### Physicochemical filtering

Because the generation model was not conditioned on activity, the raw output was filtered using five physicochemical criteria derived from the AMP literature. Peptides were retained only if they satisfied all of the following: hydrophobic residue percentage between 35% and 70%; net charge between +2 and +6 at physiological pH; amphiphilicity score H* > 0.33, calculated via helical-wheel projection with residues spaced 100° apart; Boman index between 0 and 8 kcal/mol; and isoelectric point above 10. Threshold values were established empirically by plotting each property against MIC or *IC*_50_ values for peptides in the DRAMP and RW-lexicon databases. This filtering step reduced the candidate pool from 39,846 to roughly 1,349 sequences. Analysis of physicochemical properties confirmed that the retained peptides were statistically consistent with known AMPs: their mean charge, amphiphilicity, hydrophobic residue percentage, and isoelectric point all aligned closely with the DBAASP and DRAMP training distributions. PCA and t-SNE visualisations further showed that the generated peptides occupied a region of physicochemical space broadly overlapping with known AMPs, while t-SNE indicated somewhat greater local diversity. Nevertheless, both analyses revealed that substantial regions of the AMP feature space covered by the training databases were not represented in the generated set, pointing to a limitation in sequence diversity.

#### MIC prediction

To rank the filtered candidates, a Bidirectional LSTM (Bi-LSTM) regression model was developed. The sequence branch processed tokenized, embedded amino acid sequences through a single bidirectional LSTM layer; in parallel, three sets of QSAR descriptors—amino acid composition, Composition–Transition–Distribution (CTD) features, and Quasi-Sequence Order (QSO) descriptors—were each processed through separate two-layer fully connected networks before being concatenated with the sequence representation. A sigmoid output activation was used to constrain predictions to the normalised [0, 1] range, which were subsequently mapped back to MIC values in µM via inverse log–min–max normalisation. The model was trained on 728 peptides from the DRAMP database carrying MIC values against *E. coli*, with an 80/20 train–test split. Hyperparameter optimisation was again performed by random search.

From the perspective of the present study, sequence-diversity analysis revealed that the generative model; trained exclusively on the approximately 14,000 sequences in DBAASP; produced candidates that clustered tightly around known AMP sequence motifs. The 3-mer cosine similarity between generated and training sets was 0.73, and the most enriched motifs (RRR, KKK, LKK, KKL, WRR) were drawn directly from the most frequent patterns in DBAASP. The generated peptides therefore explored only a shallow region of the broader AMP sequence space. These combined observations; adequate MIC predictive performance but insufficient generative diversity; directly motivated the transfer-learning strategy from the large-scale AMPSphere dataset described in the main text.

The second stage of the predecessor pipeline used a Martini 3 coarse-grained molecular dynamics workflow to rank the LSTM-generated candidates above by their predicted free energy of bilayer insertion before peptides were committed to synthesis. Coarse-grained bacterial (DOPE:DOPG, 7:3) and mammalian (DOPC:CHOL, 7:3) bilayers were built in COBY within a 20 × 20 × 20 nm box, solvated with Martini 3 water and 150 M NaCl, and equilibrated to a 2 *µ*s production run at 310 K and 1 bar in GROMACS. Each candidate sequence was modelled atomistically with PEP-FOLD3, mapped to a coarse-grained representation using Martinize2 and Vermouth, and inserted above the bacterial bilayer. A steered molecular dynamics pull along the membrane normal generated 256 evenly spaced umbrella windows, each simulated for 10.24 ns, and the biased histograms were unwrapped to a potential of mean force using the Weighted Histogram Analysis Method.

The workflow was validated against curated antimicrobial peptide datasets, with predicted insertion energies correlating with experimental potency for short, surface-acting peptides. The benchmarked filter was then applied to the top twenty LSTM candidates from the work described above. Four sequences spanning the low and high energy regions of the predicted PMF distribution were progressed to synthesis; three displayed activity against *E. coli* MG1655, with one (peptide 11, MIC = 0.125 *µ*g/mL) surpassing polymyxin B under identical assay conditions. The present work expands the chemical diversity of candidates entering this filter through genomic pre-training.

#### Weighted Levenshtein Distance

**Colour Key:** 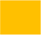 Substitution (cost = 1) 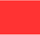 Deletion / 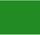 Insertion (cost = 2) □ Match (cost = 0)

*Note: Dashes (–) indicate gap characters introduced by the optimal alignment. Colour coding applied to query (generated) sequence; DBAASP reference shown in italics below. Scoring scheme: match = 0, substitution = 1, insertion/deletion = 2*.

**FIG. S1.**
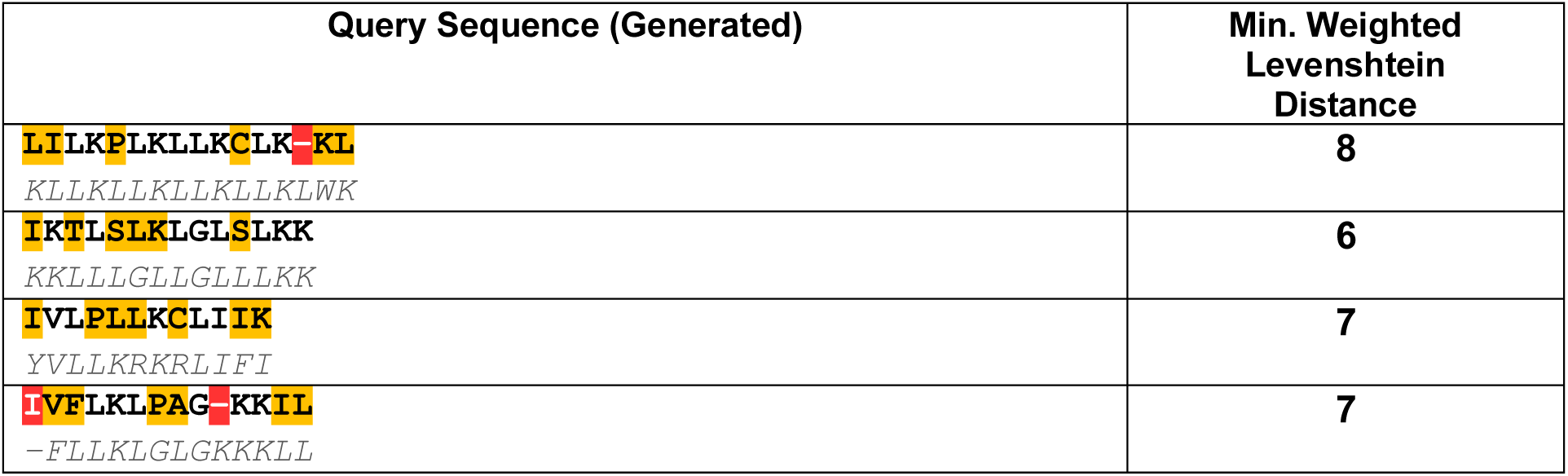
Weighted Levenshtein alignment of the top five highest predicted-activity peptide candidates reported by Gonteri. Coloured residues indicate the edit operations required to align each generated query sequence to its nearest DBAASP reference sequence, with substitution cost = 1 and insertion/deletion cost = 2.

### S3. MIC PREDICTION

#### A. Model Architecture Details

##### MLP

The input vector was passed through a series of fully connected layers with GELU activations:

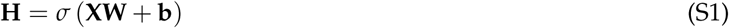

where **H** is the hidden layer output, **X** is the input, and **W**, **b** are the learned weight matrix and bias. The GELU activation is defined as:

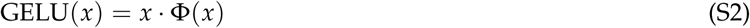

where Φ(*x*) is the cumulative distribution function of the standard normal distribution^S19^.

##### BiLSTM

At each sequence position *t*, the LSTM computes:

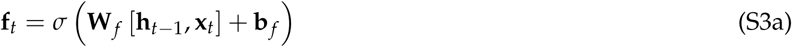

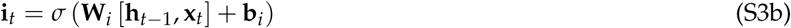

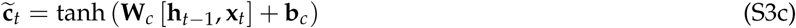

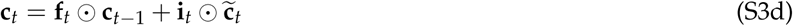

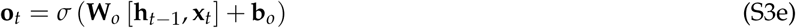

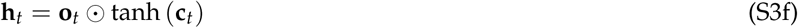

(S3f)

where **f***_t_*, **i***_t_*, and **o***_t_* are the forget, input, and output gates; **c***_t_*is the candidate cell state; **c***_t_* and **h***_t_*are the cell and hidden states at position *t*; ⊙ denotes element-wise multiplication; and **x***_t_* is the ESM-2 embedding of residue *t*. Two independent LSTM layers process the sequence in forward and reverse directions, with hidden states concatenated at each position:

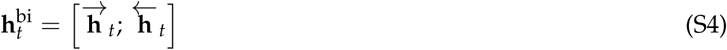

where 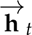 and 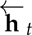 denote the forward and backward hidden states at position *t*, and [·; ·] denotes vector concatenation.

###### 1. MIC predictor model

SI*_m_ic_h_yperparametersEightcandidatearchitectures*(*TablesS*11*andS*12)*wereevaluatedontheE. colitestsetacrossregressionaccuracy*(R MAE, RMSE), rank-order performance (Spearman’s *ρ*), and drug-discovery enrichment metrics (*EF*@10%, and BEDROC, *α* = 20). Model selection was driven by a combined score comprising Spearman’s *ρ* and *EF*@10% (Eq. (**??**)) because virtual screening is fundamentally a ranking problem; correct prioritisation of potent peptides is more relevant for experimental follow-up compared to minimising mean absolute error. *R*^2^, MAE, RMSE, and BEDROC are reported for benchmarking against prior work. The results from 3 random seeds are shown in Table S1. Across all models, performance is consistent with independent-set benchmarks reported for *E. coli* MIC regression in the literature (Table S1), including those using ESM-2 embeddings^S67^ and larger held-out sets^S68^. Direct numerical comparison is approximate due to unit differences between studies. The MLP (ESM+QSAR) and the BiLSTM variant achieved the highest combined scores among all single models (Table S1). Despite its marginally higher mean value, the BiLSTM showed greater seed-to-seed variability than the MLP, indicating reduced reproducibility. Moreover, training speed differed markedly: the MLP converged within seconds, whereas BiLSTM variants required several minutes, reflecting the computational overhead of sequential hidden-state propagation inherent to recurrent architectures.

**Table S1.** Performance of MIC prediction models on the *E. coli* test set across three random seeds.

| Model | R<sup>2</sup> | MAE | RMSE | Spearman $\rho$ | EF@10% | BEDROC ($\alpha = 20$) | Combined Score |
| --- | --- | --- | --- | --- | --- | --- | --- |
| MLP (ESM+QSAR) | 0.365 ± 0.132 | 0.426 ± 0.024 | 0.599 ± 0.098 | 0.639 ± 0.050 | 4.82 ± 0.38 | 0.561 ± 0.077 | 6.095 ± 0.289 |
| MLP (ESM-only) | 0.413 ± 0.098 | 0.425 ± 0.012 | 0.574 ± 0.046 | 0.626 ± 0.067 | 4.33 ± 0.43 | 0.542 ± 0.093 | 5.578 ± 0.469 |
| BiLSTM (+QSAR) | 0.350 ± 0.144 | 0.431 ± 0.017 | 0.606 ± 0.105 | 0.643 ± 0.039 | 4.37 ± 0.46 | 0.544 ± 0.092 | 5.654 ± 0.464 |
| BiLSTM | 0.366 ± 0.110 | 0.438 ± 0.005 | 0.597 ± 0.057 | 0.611 ± 0.050 | 5.06 ± 0.78 | 0.597 ± 0.103 | 6.279 ± 0.728 |
| BiLSTM Seq | 0.310 ± 0.009 | 0.472 ± 0.033 | 0.625 ± 0.050 | 0.524 ± 0.024 | 4.01 ± 0.70 | 0.470 ± 0.100 | 5.055 ± 0.690 |
| BiLSTM Seq (+QSAR) | 0.334 ± 0.072 | 0.444 ± 0.012 | 0.614 ± 0.061 | 0.578 ± 0.046 | 4.58 ± 0.86 | 0.539 ± 0.118 | 5.733 ± 0.896 |
| BiLSTM Seq (+ESM) | 0.448 ± 0.113 | 0.415 ± 0.032 | 0.557 ± 0.070 | 0.654 ± 0.081 | 4.54 ± 0.60 | 0.526 ± 0.116 | 5.851 ± 0.646 |
| BiLSTM Seq (+ESM+QSAR) | 0.386 ± 0.113 | 0.439 ± 0.038 | 0.589 ± 0.079 | 0.603 ± 0.066 | 4.38 ± 0.46 | 0.559 ± 0.036 | 5.583 ± 0.329 |

All models performed worse on the *E. coli* dataset than on the broader Gram-negative (GN) dataset used for initial hyperparameter selection (Table S13), despite its apparent biological homogeneity. This likely reflects data limitations rather than biological simplicity: single-species datasets are typically smaller, making models more susceptible to overfitting and higher variance, whereas broader GN datasets benefit from larger sample sizes and improved generalisation. This is consistent with Bonifacio-Velez de Villa *et al.* who report that species-specific models outperform GN-level models only when species subset is sufficiently large^S69^.

###### 2. Calibration and Out-of-Distribution Robustness

To assess calibration, log_10_(MIC) [µg mL^−1^] distributions on the *E. coli* test set were compared across architectures (Figure S2). All models produced distributions centred close to the *E. coli* test set mean, consistent with well-calibrated central tendency across architectures. MLPs displayed markedly narrower distributions than BiLSTM models. Notably, BiLSTMs predicted more negative log_10_(MIC) values, corresponding to greater antimicrobial activity — a property directly relevant to downstream candidate selection. Despite this, no single model captured the full distribution of the training dataset.

**FIG. S2.**
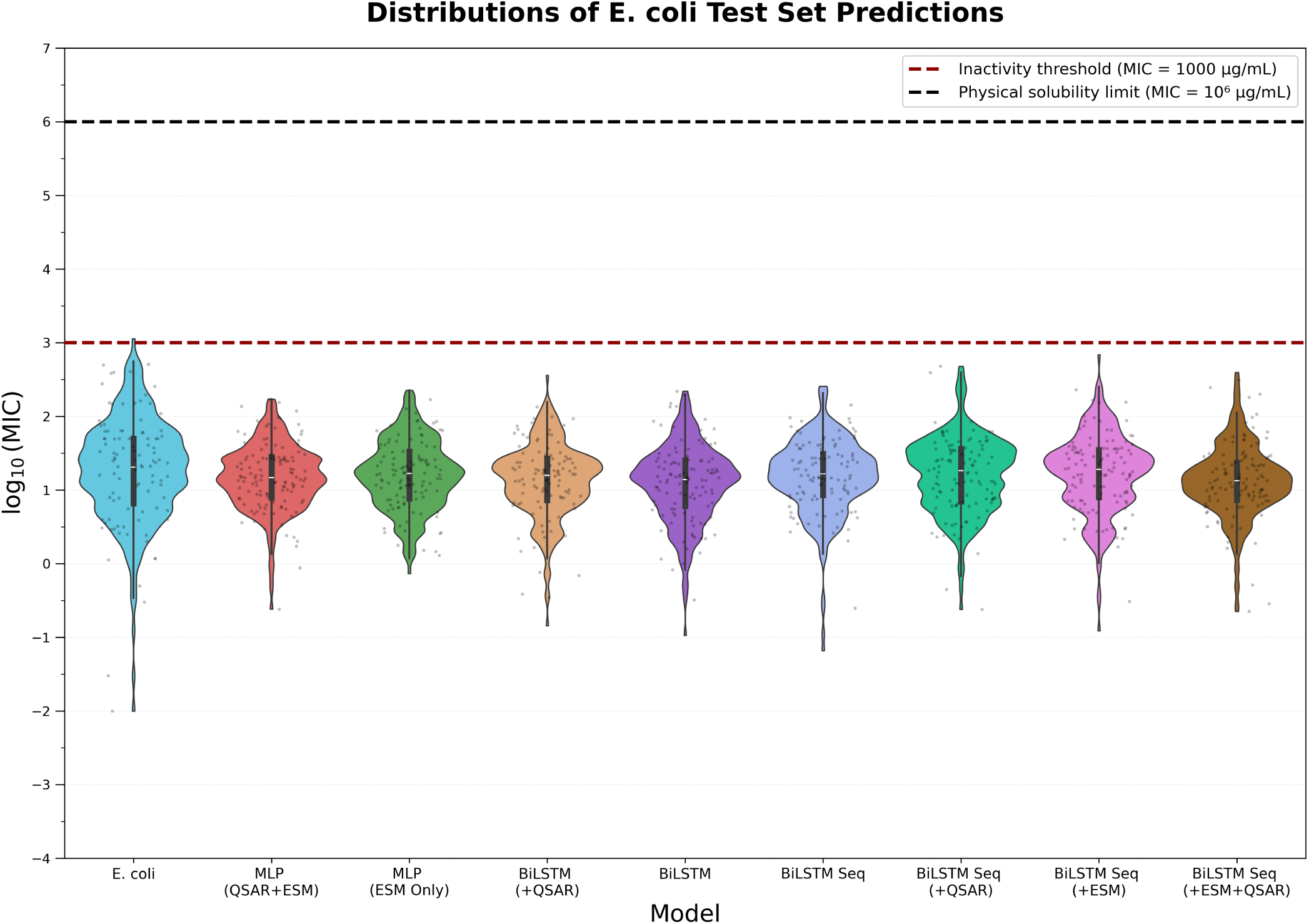
Distribution of predicted log_10_(MIC) values (µg mL^−1^) on the *E. coli* test set for all evaluated model architectures, alongside the experimental reference distribution. The experimental *E. coli* test set mean is *µ* = 1.27 ± 0.70 (*n* = 250). Predicted means and standard deviations are: MLP (ESM+QSAR) 1.17 ± 0.44; MLP (ESM-only) 1.20 ± 0.47; BiLSTM (+QSAR) 1.13 ± 0.49; BiLSTM 1.08 ± 0.52; BiLSTM Seq 1.18 ± 0.51; BiLSTM Seq (+QSAR) 1.21 ± 0.51; BiLSTM Seq (+ESM) 1.23 ± 0.53; BiLSTM Seq (+ESM+QSAR) 1.12 ± 0.48. All architectures show well-calibrated central tendency; BiLSTM models predict a broader range of low-MIC values than MLPs, which is relevant for downstream candidate selection. Individual points represent a random sample of 500 peptides per model for visualisation clarity. The red dashed line denotes the inactivity threshold (log_10_(MIC) = 3 µg mL^−1^), above which activity is considered practically irrelevant. The black dashed line indicates the physical solubility limit (log_10_(MIC) = 6 µg mL^−1^), beyond which the required peptide mass exceeds the solution volume.

Following this, robustness to out-of-distribution sequences was assessed by applying all eight predictors to 20,000 randomly generated peptides (convergence confirmed at ∼10,000 samples; Figures S14 and S15). Substantially higher predicted MICs to random peptides were predicted across all models than to the *E. coli* test set, demonstrating that models have learned AMP-discriminating features rather than regressing to the training mean.

Analysis of extreme predictions revealed a marked divergence in model behaviour (Tables S17 and S18). The five sequences assigned the highest predicted MICs were short pentapeptides dominated by hydrophobic residues (FIFMI, MVFMI, MFIYI). As shown in Figure S16, these sequences fell outside DRAMP *E. coli* physicochemical bounds across instability index, net charge, and Boman index and the majority were flagged by QSAR-containing MLP models with large inter-seed standard deviations. Conversely, the five sequences receiving the lowest predicted MICs (Figure S17) were identified exclusively by BiLSTM models with substantially smaller inter-seed standard deviations. Such sequences consisted of an N-terminal cysteine, and predominately remained within *E. coli* physicochemical bounds across all six descriptors. This asymmetry—QSAR-containing models excelling at flagging implausible sequences, BiLSTMs producing credible low-MIC predictions for structurally plausible candidates—reflected complementary feature capture. Physicochemical descriptors may provide a global viability filter, while ESM-2 per-residue context captures subtle activity-associated motifs. This motivated ensemble construction to improve MIC predictions.

**FIG. S3.**
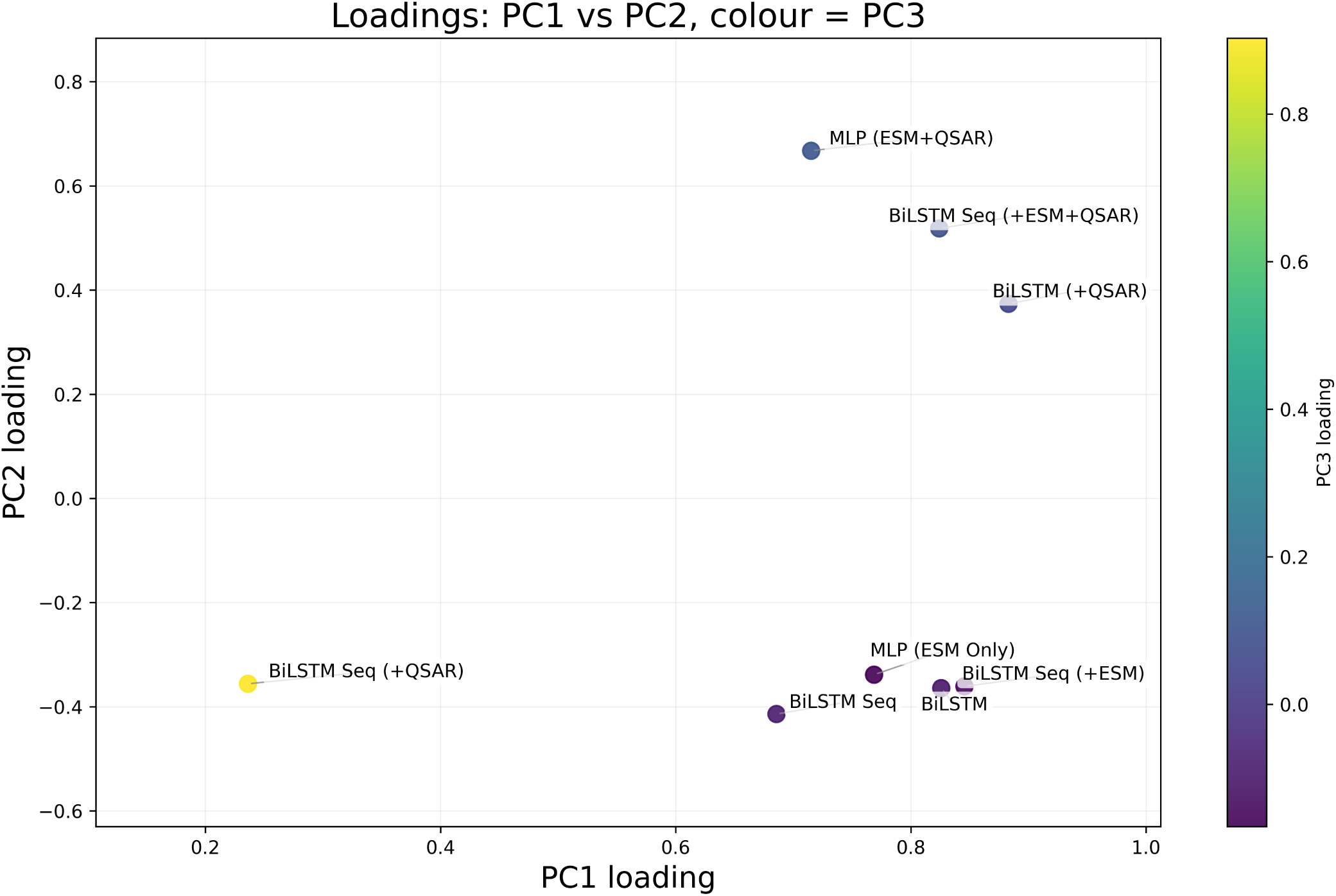
PCA of MIC predictor loadings across all evaluated models. PCA of the loading vectors reveals two main features: which models are correlated, as reflected by clustering, and which contribute most strongly to the predictions, as indicated by higher loadings.

###### 3. Ensemble Construction and Final Predictor

PCA of model prediction profiles confirmed the idea mentioned above (Figures S3 and S18–S20): loadings separated into QSAR-containing and non-QSAR clusters, indicating partial orthogonality. All ensemble strategies improved combined score relative to the best single model (Table S21). Since the full NNLS ensemble showed the highest test-set variability (*σ >* 1.0), the pairwise MLP (ESM+QSAR) + BiLSTM was selected for its balance of performance and reproducibility. The weights for the linear combination of model predictions were estimated using NNLS independently across three random seeds and then averaged. Averaging the NNLS weights across seeds yielded the final ensemble predictor (Eq. (S5); Table S19):

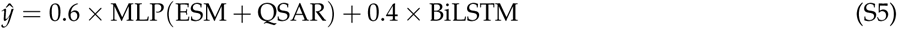

#### B. Model Hyperparameters

**Table S2.**
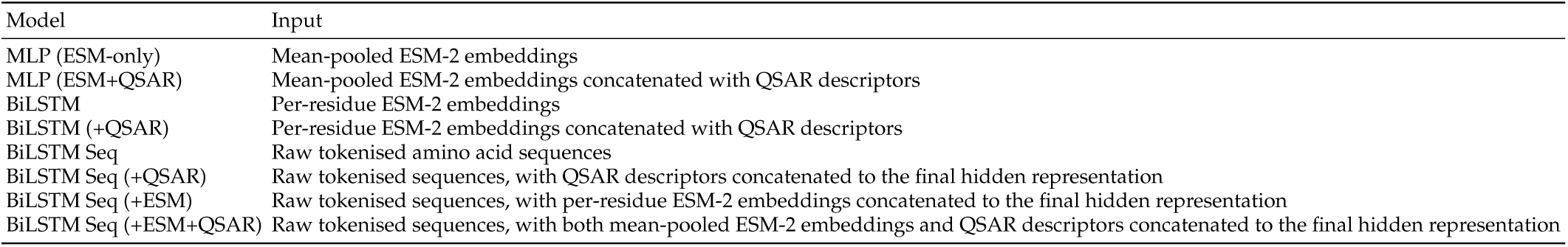
Nomenclature of Models.

| Model | Input |
| --- | --- |
| MLP (ESM-only) | Mean-pooled ESM-2 embeddings |
| MLP (ESM+QSAR) | Mean-pooled ESM-2 embeddings concatenated with QSAR descriptors |
| BiLSTM | Per-residue ESM-2 embeddings |
| BiLSTM (+QSAR) | Per-residue ESM-2 embeddings concatenated with QSAR descriptors |
| BiLSTM Seq | Raw tokenised amino acid sequences |
| BiLSTM Seq (+QSAR) | Raw tokenised sequences, with QSAR descriptors concatenated to the final hidden representation |
| BiLSTM Seq (+ESM) | Raw tokenised sequences, with per-residue ESM-2 embeddings concatenated to the final hidden representation |
| BiLSTM Seq (+ESM+QSAR) | Raw tokenised sequences, with both mean-pooled ESM-2 embeddings and QSAR descriptors concatenated to the final hidden representation |

This section presents the results of the hyperparameter grid searches for all MIC prediction models, together with additional optimisation analyses for the selected MLP (ESM+QSAR) model. Initial hyperparameter sweeps were performed on the broader Gram-negative (GN) dataset and are summarised in the tables below. The corresponding figures show the top GN-ranked configurations retrained and evaluated on the *E. coli* dataset used for final model selection.

Standard evaluation plots. Unless otherwise stated, all training figures consist of three panels: (left) training and validation loss curves with validation Spearman rank correlation (*ρ*), (middle) predicted versus actual log_10_(MIC) values with the identity line, and (right) enrichment curves assessing early retrieval performance (BEDROC). These panels are used consistently across all model sweeps to enable direct comparison.

The full BEDROC equations we used are,

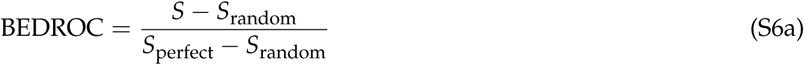

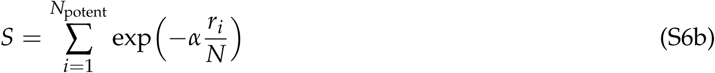

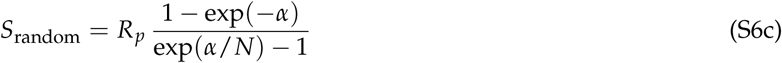

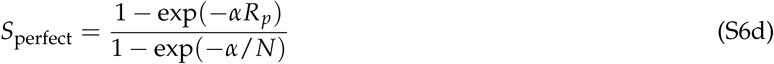

where:

- *N* is the total number of sequences in the ranked list;
- *r_i_* is the rank of the *i*-th potent sequence, ordered from most to least potent (rank 1 = most potent);
- *R_p_* = *N*_potent_/*N* is the fraction of potent sequences in the dataset;
- *α* controls how strongly early retrieval is prioritised, with larger values assigning greater Boltzmann weight to top-ranked peptides.

*α* = 20 was adopted following the recommendation of Truchon and Bayly^S24^, who show that for *α* = 20, more than 80%of the BEDROC score is concentrated within the first 10% of the ranked list. This makes it well-suited to practical screening settings in which only a small fraction of candidates can be taken forward for experimental validation.

**Table S3.**
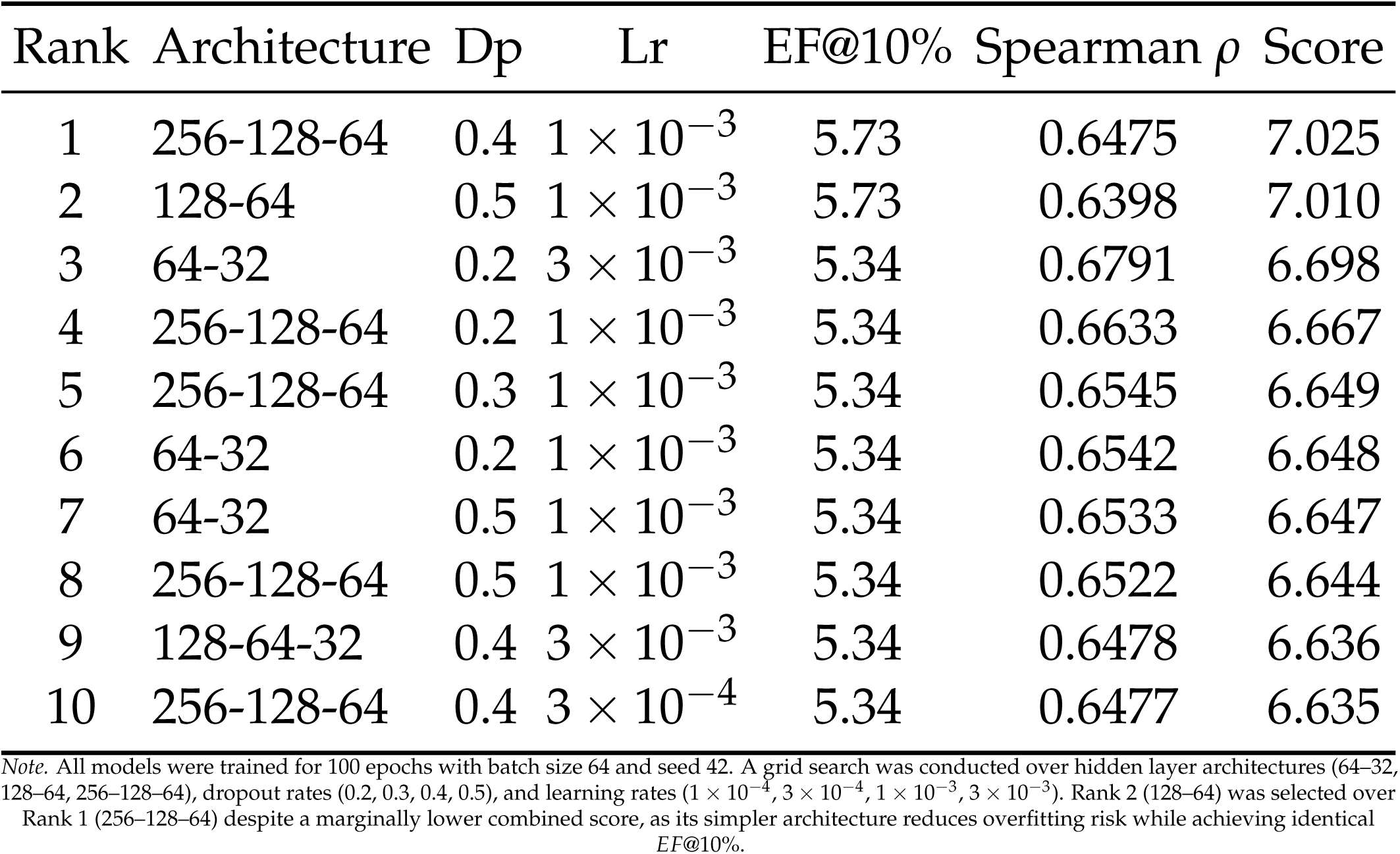
Top-10 MLP (ESM+QSAR) hyperparameter configurations ranked by combined score on the Gram-negative test set. Dp = dropout; Lr = learning rate; Score = EF@10% + 2*ρ*.

**FIG. S4.**
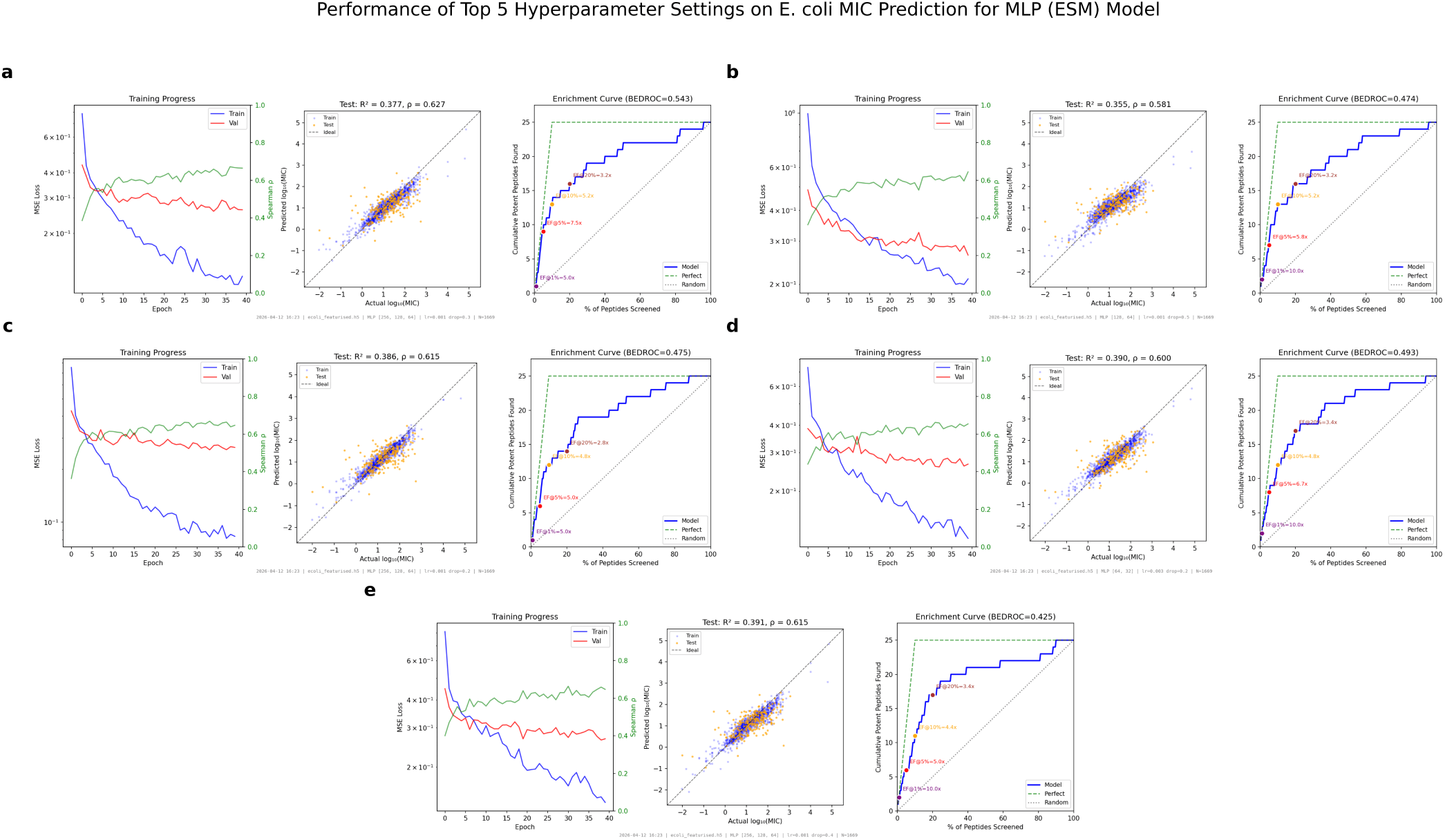
Performance of the top-5 MLP (ESM+QSAR) configurations — selected by combined score on the Gram-negative hyperparameter sweep — when retrained and evaluated on the *E. coli* peptide subset. Model (b), corresponding to rank 2 in the GN dataset, was selected as the final model using the evaluation framework mentioned in II A 5.

**Table S4.**
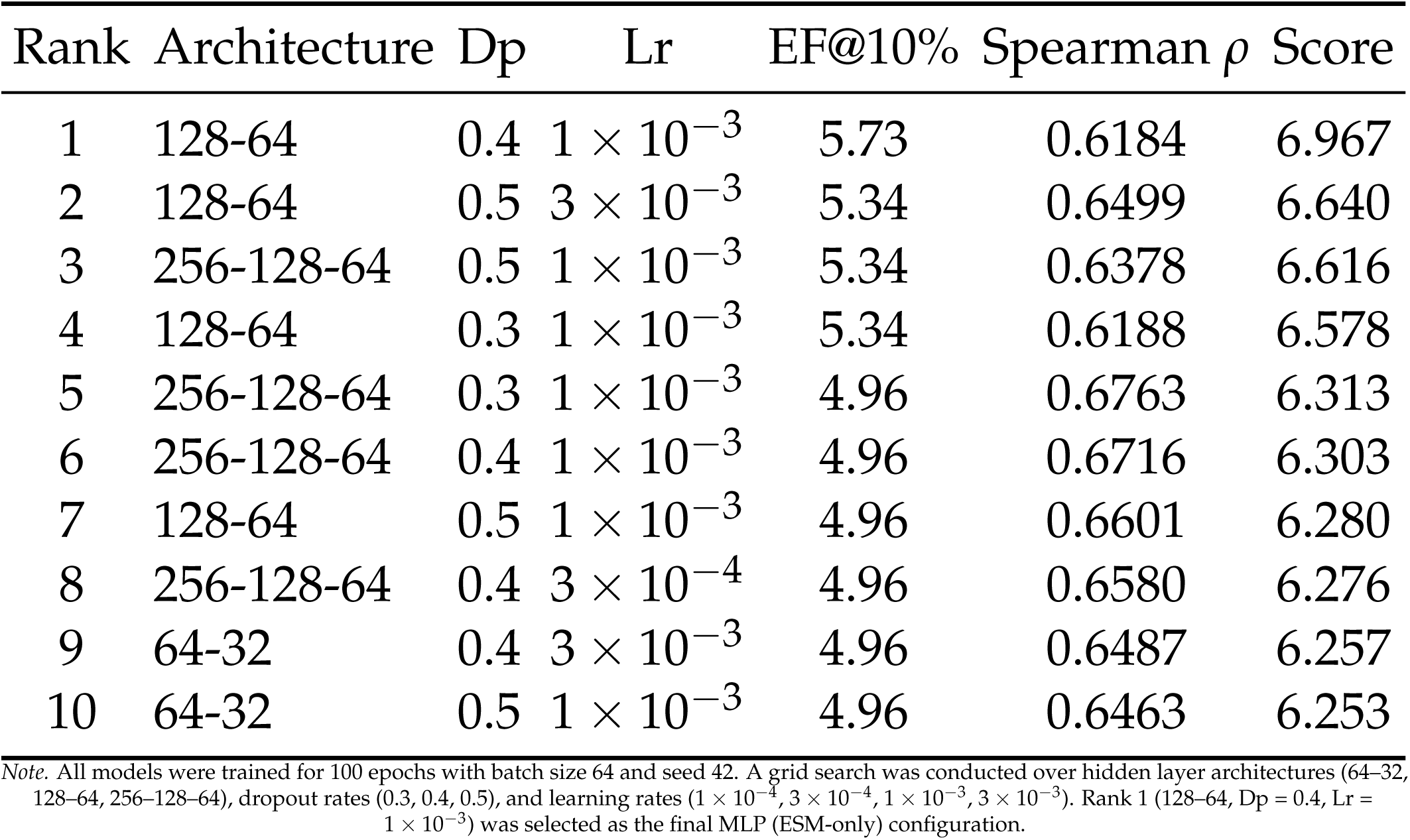
Performance of the top-10 MLP (ESM-only) hyperparameter configurations ranked by combined score on the Gram-negative test set. Dp = dropout; Lr = learning rate; Score = EF@10% + 2*ρ*.

**Table S5.** Performance of the top-10 BiLSTM hyperparameter configurations ranked by combined score on the Gram-negative test set. Dp = dropout; Lr = learning rate; Score = EF@10% + 2*ρ*.

| Rank | MLP Arch. | LSTM Layers | LSTM Hidden | Dp | Lr | EF@10% | Spearman $\rho$ | Score |
| --- | --- | --- | --- | --- | --- | --- | --- | --- |
| 1 | 64-32 | 1 | 256 | 0.5 | $3 \times 10^{-4}$ | 5.73 | 0.6177 | 6.965 |
| 2 | 128-64 | 1 | 256 | 0.4 | $3 \times 10^{-4}$ | 5.73 | 0.5823 | 6.895 |
| 3 | 64-32 | 1 | 256 | 0.5 | $1 \times 10^{-3}$ | 5.34 | 0.6065 | 6.553 |
| 4 | 128-64 | 1 | 128 | 0.5 | $1 \times 10^{-3}$ | 5.34 | 0.5981 | 6.536 |
| 5 | 128-64 | 1 | 256 | 0.3 | $1 \times 10^{-3}$ | 5.34 | 0.5901 | 6.520 |
| 6 | 128-64 | 1 | 128 | 0.3 | $3 \times 10^{-4}$ | 4.96 | 0.6267 | 6.213 |
| 7 | 64-32 | 1 | 128 | 0.3 | $3 \times 10^{-4}$ | 4.96 | 0.6194 | 6.199 |
| 8 | 64-32 | 1 | 64 | 0.5 | $3 \times 10^{-4}$ | 4.96 | 0.6161 | 6.192 |
| 9 | 64-32 | 1 | 256 | 0.3 | $3 \times 10^{-4}$ | 4.96 | 0.6156 | 6.191 |
| 10 | 128-64 | 1 | 64 | 0.3 | $3 \times 10^{-4}$ | 4.96 | 0.6138 | 6.188 |

*Note.* Training was limited to 40 epochs due to increased computational cost relative to MLP models (batch size = 64, seed = 42). A grid search was conducted over LSTM layers (1), LSTM hidden size (64, 128, 256), MLP head architectures (64–32, 128–64, 256–128–64), dropout rates (0.3, 0.4, 0.5), and learning rates ( $1 \times 10^{-4}$ , $3 \times 10^{-4}$ , $1 \times 10^{-3}$ ). Rank 1 (MLP 64–32, LSTM hidden 256, Dp = 0.5, Lr = $3 \times 10^{-4}$ ) was selected as the final BiLSTM configuration.

**FIG. S5.**
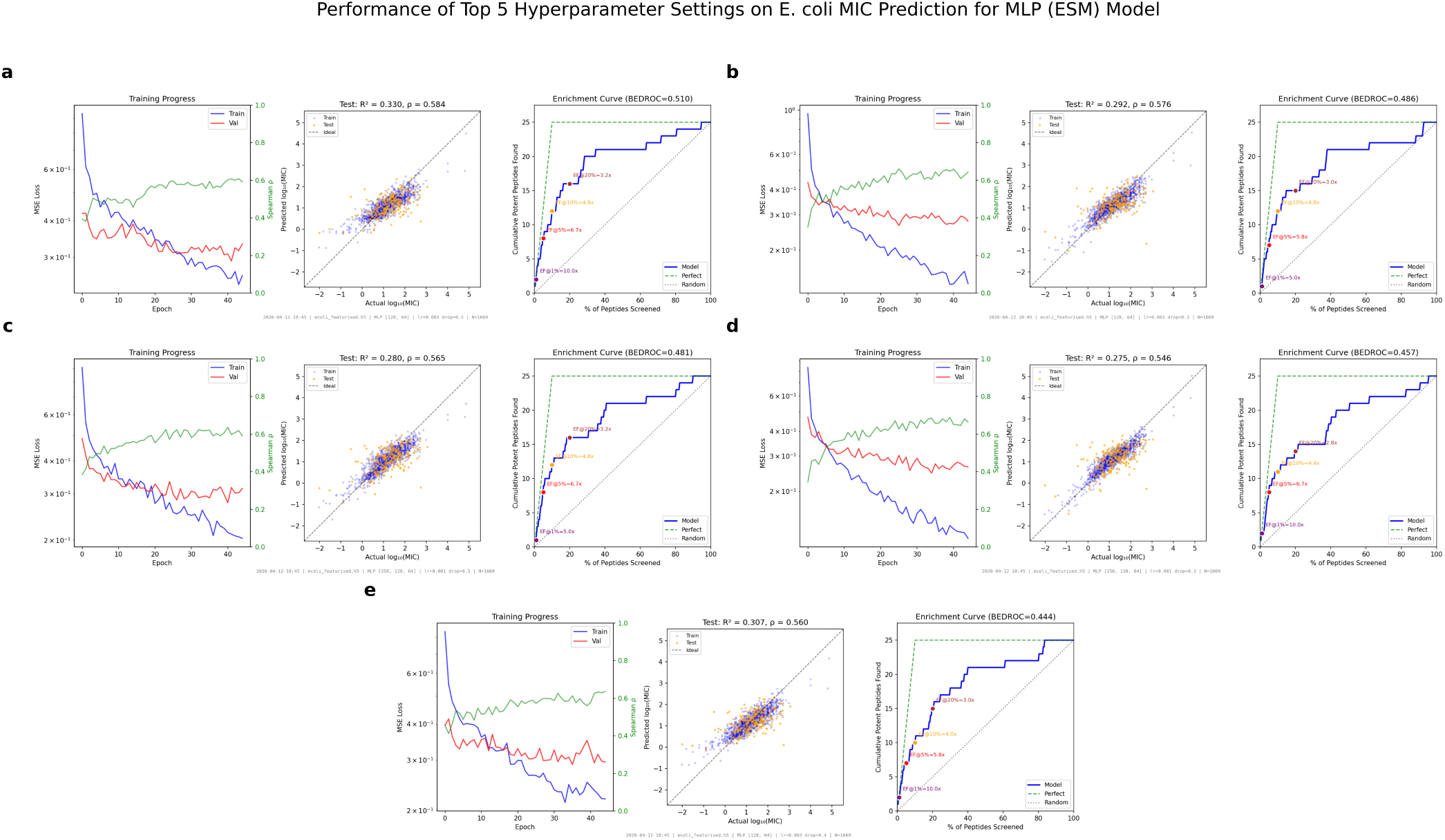
Performance of the top-5 MLP (ESM-only) configurations — selected by combined score on the Gram-negative hyperparameter sweep — when retrained and evaluated on the *E. coli* peptide subset. Model (e), corresponding to rank 1 in the GN dataset, was selected as the final model using the evaluation framework mentioned in II A 5.

**Table S6.** Performance of the top-10 BiLSTM (+QSAR) hyperparameter configurations ranked by combined score on the Gram-negative test set. Dp = dropout; Lr = learning rate; Score = EF@10% + 2*ρ*.

| Rank | MLP Arch. | LSTM Layers | LSTM Hidden | Dp | Lr | EF@10% | Spearman $\rho$ | Score |
| --- | --- | --- | --- | --- | --- | --- | --- | --- |
| 1 | 64-32 | 1 | 128 | 0.4 | $3 \times 10^{-3}$ | 5.73 | 0.6313 | 6.993 |
| 2 | 64-32 | 1 | 128 | 0.3 | $3 \times 10^{-4}$ | 5.34 | 0.6475 | 6.635 |
| 3 | 256-128-64 | 2 | 64 | 0.3 | $3 \times 10^{-4}$ | 5.34 | 0.6401 | 6.620 |
| 4 | 128-64 | 1 | 256 | 0.5 | $3 \times 10^{-3}$ | 5.34 | 0.6304 | 6.601 |
| 5 | 64-32 | 1 | 256 | 0.5 | $1 \times 10^{-3}$ | 5.34 | 0.6272 | 6.594 |
| 6 | 64-32 | 1 | 64 | 0.3 | $3 \times 10^{-3}$ | 5.34 | 0.6135 | 6.567 |
| 7 | 64-32 | 1 | 128 | 0.5 | $3 \times 10^{-3}$ | 5.34 | 0.6072 | 6.554 |
| 8 | 256-128-64 | 2 | 64 | 0.4 | $3 \times 10^{-3}$ | 5.34 | 0.6028 | 6.546 |
| 9 | 64-32 | 1 | 128 | 0.3 | $3 \times 10^{-3}$ | 5.34 | 0.5969 | 6.534 |
| 10 | 256-128-64 | 1 | 256 | 0.3 | $3 \times 10^{-3}$ | 5.34 | 0.5893 | 6.519 |

Note. All models were trained for 40 epochs with batch size 64 and seed 42. A grid search was conducted over LSTM layers (1, 2), LSTM hidden size (64, 128, 256), MLP head architectures (64-32, 128-64, 256-128-64), dropout rates (0.3, 0.4, 0.5), and learning rates ( $1 \times 10^{-4}$ , $3 \times 10^{-4}$ , $1 \times 10^{-3}$ , $3 \times 10^{-3}$ ). Rank 1 (MLP 64-32, LSTM layers 1, LSTM hidden 128, Dp = 0.4, Lr = $3 \times 10^{-3}$ ) was selected as the final BiLSTM (+QSAR) configuration.

**FIG. S6.**
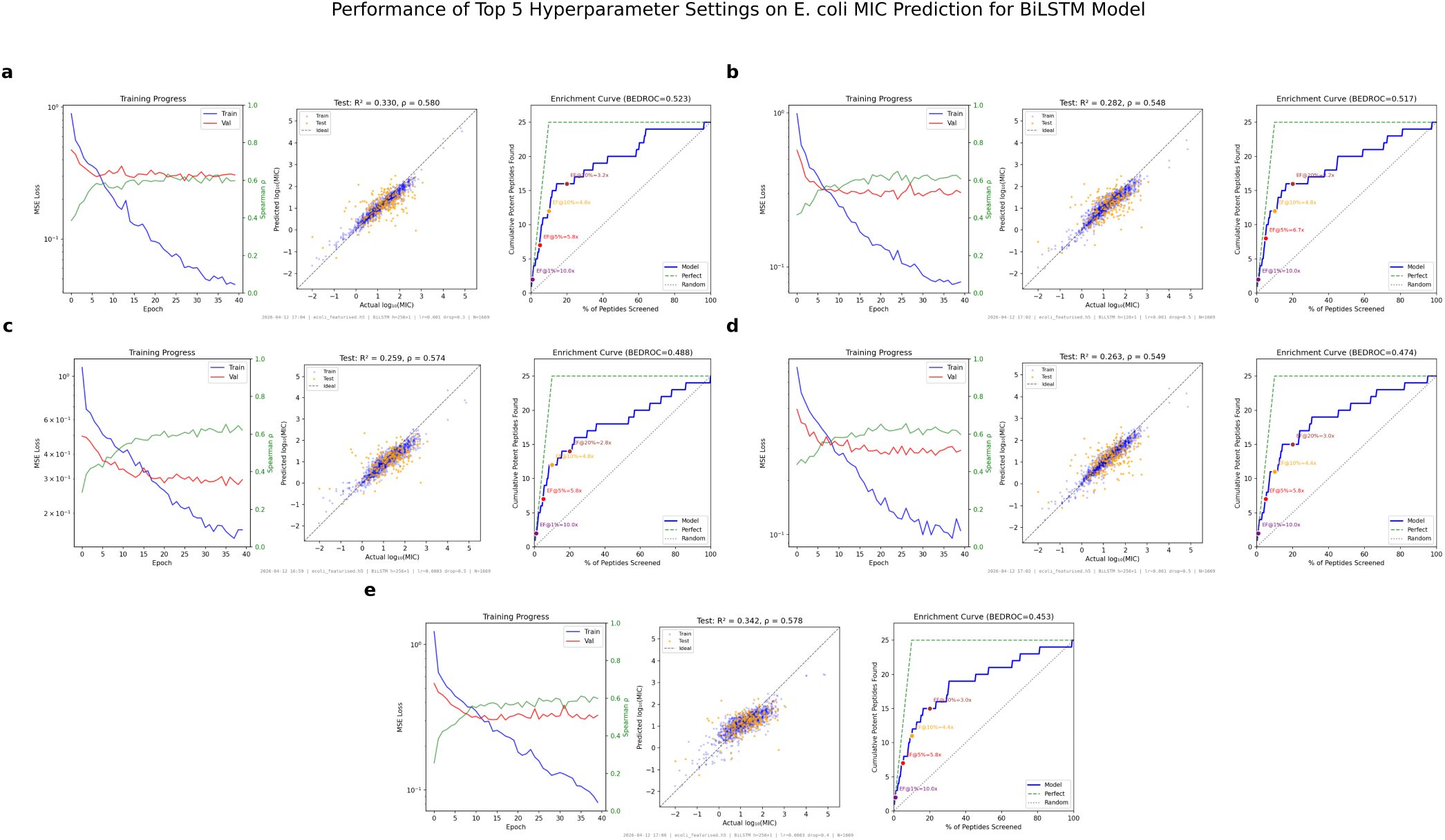
Performance of the top-5 BiLSTM configurations — selected by combined score on the Gram-negative hyperparameter sweep — when retrained and evaluated on the *E. coli* peptide subset. Model (c), corresponding to rank 1 in the GN dataset, was selected as the final model using the evaluation framework mentioned in II A 5.

**Table S7.** Performance of the top-10 BiLSTM Seq configurations hyperparameter configurations ranked by combined score on the Gram-negative test set. Dp = dropout; Lr = learning rate; Ed = embedding dimension; Score = EF@10% + 2*ρ*.

| Rank | MLP Arch. | LSTM Layers | LSTM Hidden | Dp | Lr | Ed | EF@10% | Spearman $\rho$ | Score |
| --- | --- | --- | --- | --- | --- | --- | --- | --- | --- |
| 1 | 64-32 | 1 | 256 | 0.3 | $3 \times 10^{-4}$ | 128 | 6.49 | 0.5578 | 7.606 |
| 2 | 128-64 | 1 | 256 | 0.5 | $1 \times 10^{-3}$ | 32 | 6.49 | 0.5523 | 7.595 |
| 3 | 64-32 | 1 | 256 | 0.5 | $3 \times 10^{-4}$ | 64 | 6.11 | 0.5467 | 7.203 |
| 4 | 64-32 | 1 | 256 | 0.5 | $3 \times 10^{-4}$ | 32 | 6.11 | 0.5458 | 7.202 |
| 5 | 128-64 | 1 | 64 | 0.3 | $1 \times 10^{-3}$ | 32 | 6.11 | 0.5430 | 7.196 |
| 6 | 128-64 | 1 | 128 | 0.4 | $1 \times 10^{-3}$ | 32 | 6.11 | 0.5345 | 7.179 |
| 7 | 128-64 | 1 | 128 | 0.4 | $3 \times 10^{-4}$ | 256 | 6.11 | 0.5146 | 7.139 |
| 8 | 64-32 | 1 | 128 | 0.3 | $1 \times 10^{-3}$ | 32 | 6.11 | 0.5130 | 7.136 |
| 9 | 64-32 | 1 | 128 | 0.5 | $3 \times 10^{-4}$ | 256 | 5.73 | 0.5858 | 6.902 |
| 10 | 128-64 | 1 | 256 | 0.3 | $1 \times 10^{-4}$ | 256 | 5.73 | 0.5627 | 6.855 |

Note. All models were trained for 40 epochs with batch size 64 and seed 42. A grid search was conducted over LSTM layers (1), LSTM hidden size (64, 128, 256), MLP head architectures (64–32, 128–64), dropout rates (0.3, 0.4, 0.5), learning rates ( $1 \times 10^{-4}$ , $3 \times 10^{-4}$ , $1 \times 10^{-3}$ ), and embedding dimensions (32, 64, 128, 256). Rank 3 (MLP 64–32, LSTM hidden 256, Dp = 0.5, Lr = $3 \times 10^{-4}$ , Ed = 64) was selected over Ranks 1 and 2 for its more favourable train–validation loss profile (S8), indicating better generalisation despite a marginally lower combined score.

**FIG. S7.**
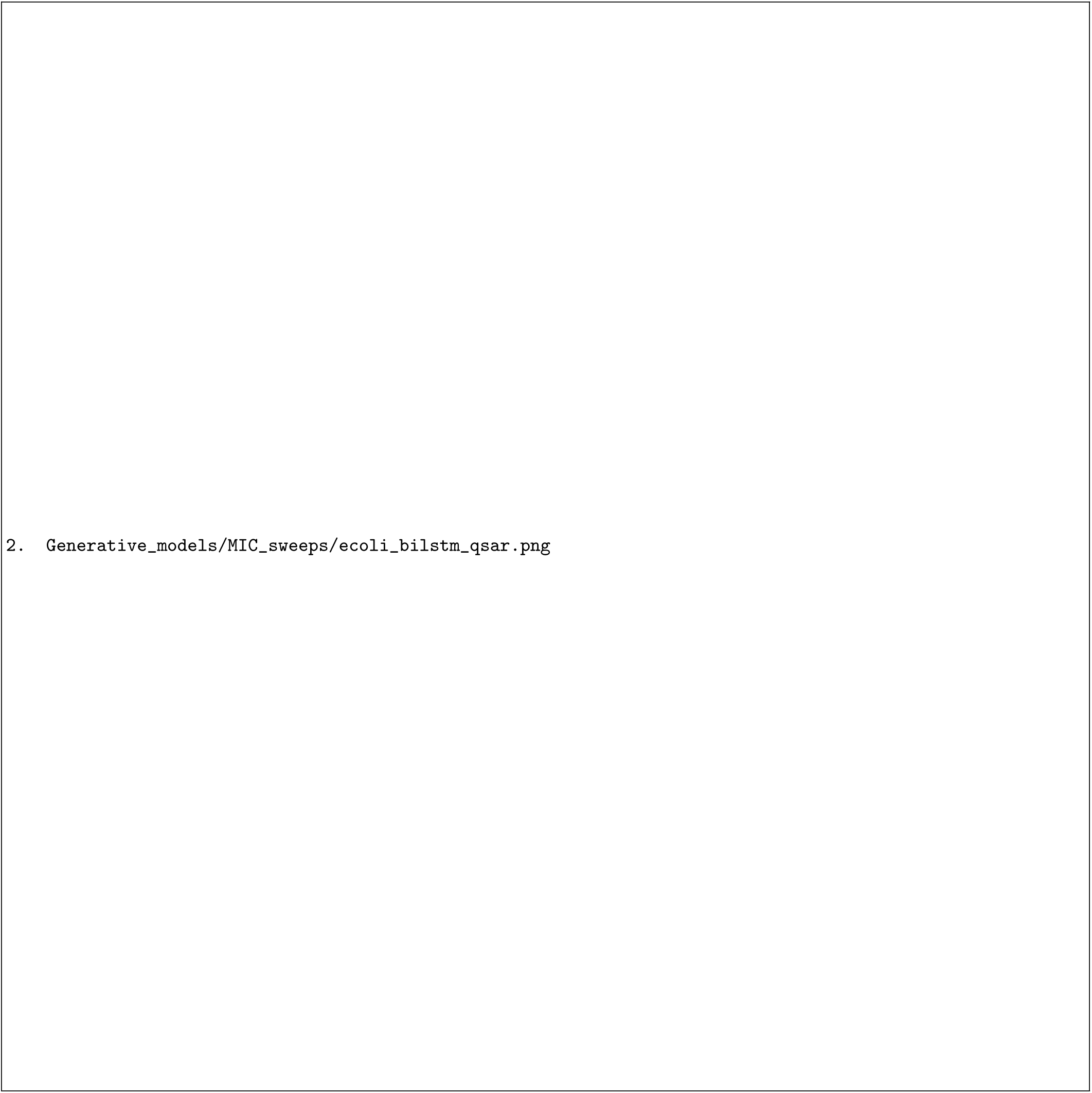
Performance of the top-5 BiLSTM (+QSAR) configurations — selected by combined score on the Gram-negative hyperparameter sweep — when retrained and evaluated on the *E. coli* peptide subset. Model (d), corresponding to rank 1 in the GN dataset, was selected as the final model using the evaluation framework mentioned in II A 5.

**FIG. S8.**
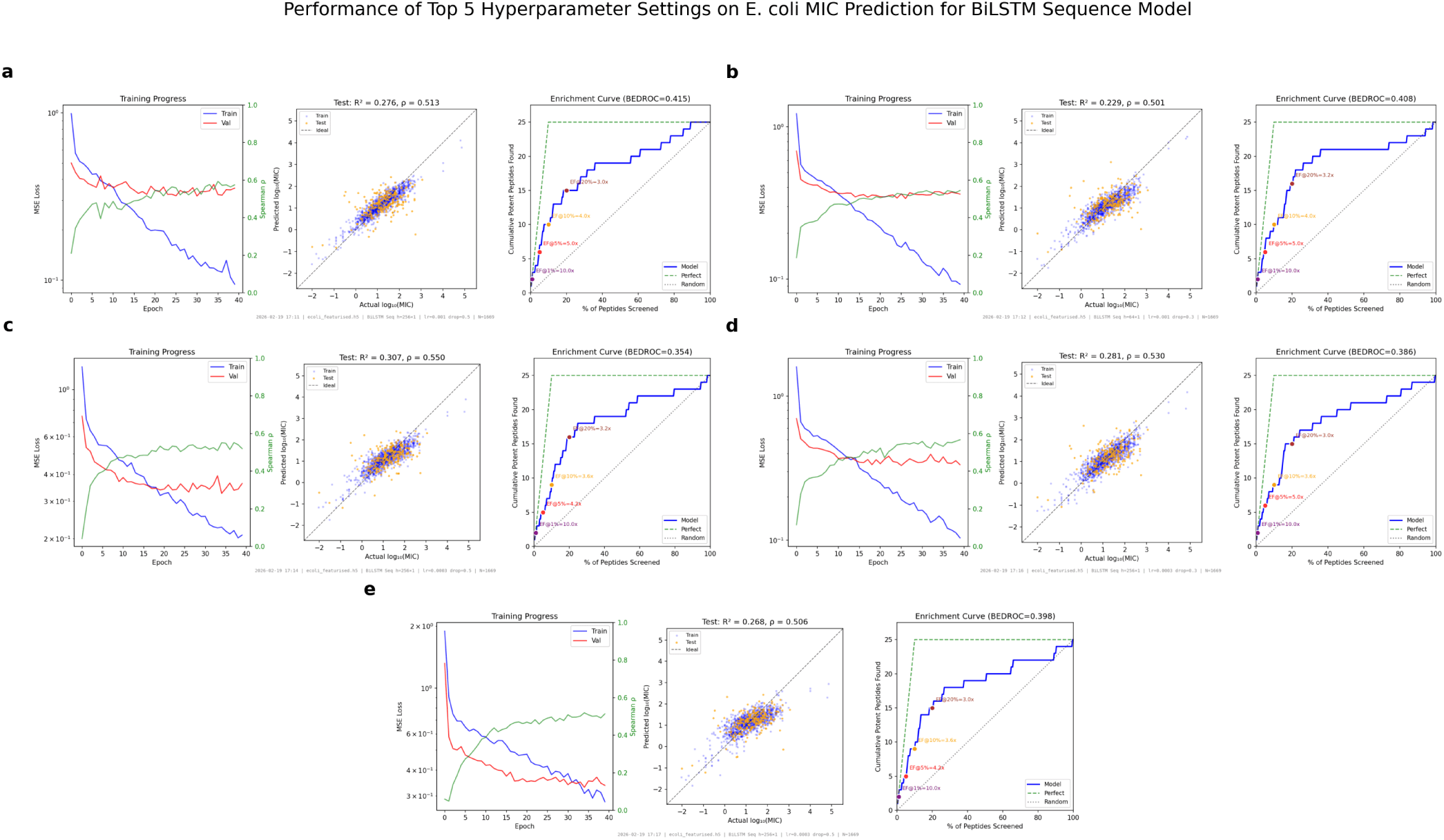
Performance of the top-5 BiLSTM Seq configurations — selected by combined score on the Gram-negative hyperparameter sweep — when retrained and evaluated on the *E. coli* peptide subset. Model (c), corresponding to rank 3 in the GN dataset, was selected as the final model using the evaluation framework mentioned in II A 5.

**Table S8.** Performance of the top-10 BiLSTM Seq (+QSAR) hyperparameter configurations ranked by combined score on the Gram-negative test set. Dp = dropout; Lr = learning rate; Ed = embedding dimension; Score = EF@10% + 2*ρ*.

| Rank | MLP Arch. | LSTM Layers | LSTM Hidden | Dp | Lr | Ed | EF@10% | Spearman $\rho$ | Score |
| --- | --- | --- | --- | --- | --- | --- | --- | --- | --- |
| 1 | 128-64 | 1 | 64 | 0.5 | $3 \times 10^{-3}$ | 64 | 5.73 | 0.6575 | 7.045 |
| 2 | 128-64 | 1 | 128 | 0.3 | $3 \times 10^{-3}$ | 64 | 5.73 | 0.6342 | 6.998 |
| 3 | 64-32 | 1 | 128 | 0.5 | $3 \times 10^{-3}$ | 64 | 5.73 | 0.6187 | 6.967 |
| 4 | 64-32 | 1 | 128 | 0.5 | $3 \times 10^{-3}$ | 32 | 5.73 | 0.6141 | 6.958 |
| 5 | 128-64 | 1 | 128 | 0.4 | $1 \times 10^{-3}$ | 32 | 5.34 | 0.6401 | 6.620 |
| 6 | 128-64 | 1 | 128 | 0.5 | $1 \times 10^{-3}$ | 32 | 5.34 | 0.6313 | 6.603 |
| 7 | 64-32 | 1 | 64 | 0.5 | $3 \times 10^{-3}$ | 64 | 5.34 | 0.6176 | 6.575 |
| 8 | 64-32 | 1 | 128 | 0.4 | $3 \times 10^{-3}$ | 32 | 5.34 | 0.6175 | 6.575 |
| 9 | 64-32 | 1 | 128 | 0.3 | $3 \times 10^{-3}$ | 32 | 5.34 | 0.6077 | 6.555 |
| 10 | 128-64 | 1 | 64 | 0.4 | $3 \times 10^{-3}$ | 64 | 5.34 | 0.6053 | 6.551 |

Note. All models were trained for 40 epochs with batch size 64 and seed 42. A grid search was conducted over LSTM layers (1), LSTM hidden size (64, 128, 256), MLP head architectures (64–32, 128–64), dropout rates (0.3, 0.4, 0.5), learning rates ( $1 \times 10^{-4}$ , $3 \times 10^{-4}$ , $1 \times 10^{-3}$ , $3 \times 10^{-3}$ ), and embedding dimensions (32, 64, 128, 256). Rank 3 (MLP 64–32, LSTM hidden 128, Dp = 0.5, Lr = $3 \times 10^{-3}$ , Ed = 64) was selected over Ranks 1 and 2 for its more favourable train-validation loss profile (S9), indicating better generalisation despite a marginally lower combined score.

**FIG. S9.**
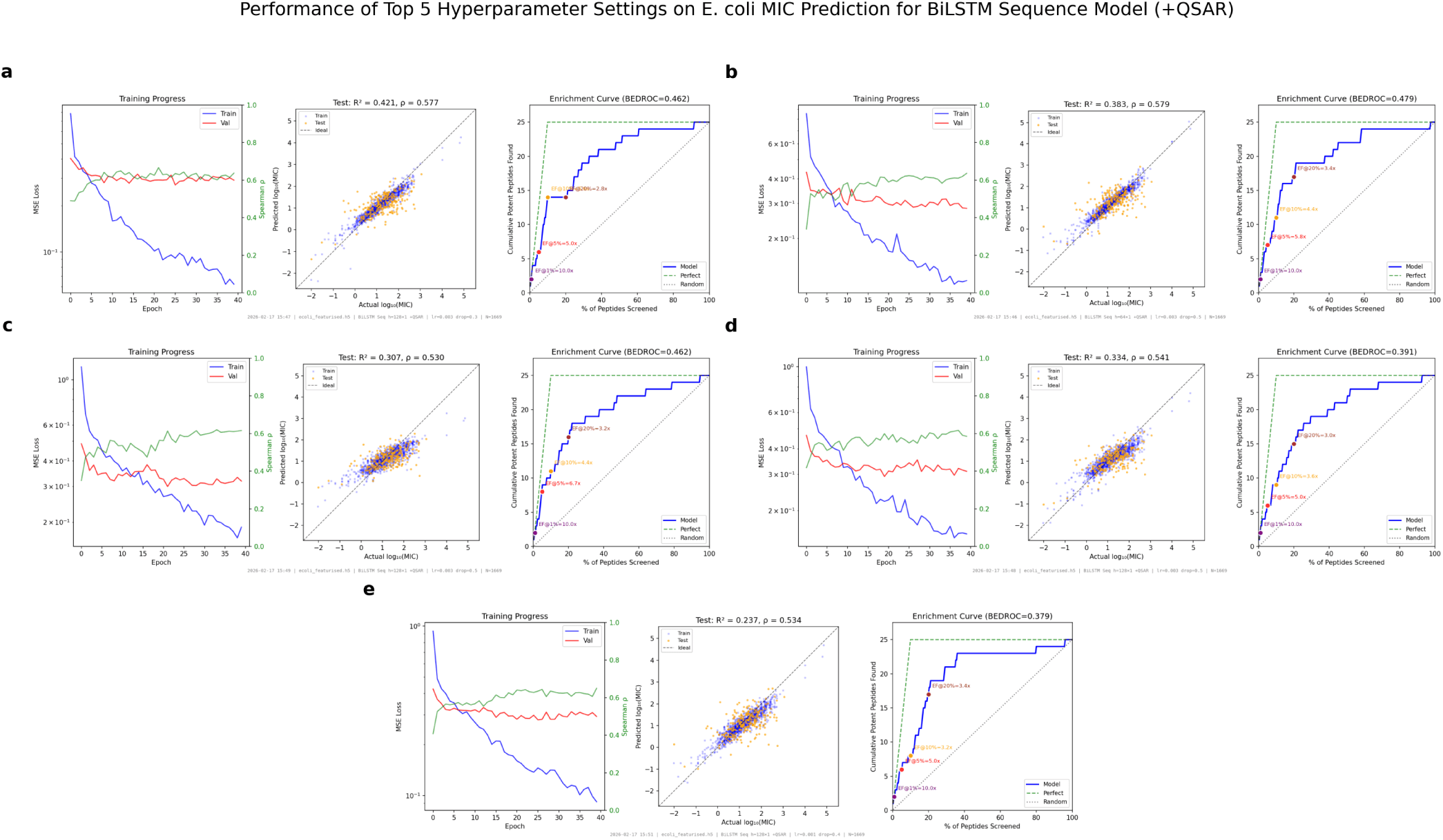
Performance of the top-5 BiLSTM Seq (+QSAR) configurations — selected by combined score on the Gram-negative hyperparameter sweep — when retrained and evaluated on the *E. coli* peptide subset. Model (c), corresponding to rank 3 in the GN dataset, was selected as the final model using the evaluation framework mentioned in II A 5.

**Table S9.** Performance of the top-10 BiLSTM Seq (+ESM) hyperparameter configurations ranked by combined score on the Gram-negative test set. Dp = dropout; Lr = learning rate; Ed = embedding dimension; Score = EF@10% + 2*ρ*.

| Rank | MLP Arch. | LSTM Layers | LSTM Hidden | Dp | Lr | Ed | EF@10% | Spearman $\rho$ | Score |
| --- | --- | --- | --- | --- | --- | --- | --- | --- | --- |
| 1 | 128-64 | 1 | 256 | 0.3 | $3 \times 10^{-3}$ | 64 | 6.11 | 0.5977 | 7.305 |
| 2 | 64-32 | 1 | 64 | 0.3 | $3 \times 10^{-3}$ | 64 | 5.73 | 0.6263 | 6.983 |
| 3 | 64-32 | 1 | 128 | 0.3 | $1 \times 10^{-3}$ | 128 | 5.73 | 0.5767 | 6.883 |
| 4 | 128-64 | 1 | 64 | 0.4 | $1 \times 10^{-3}$ | 32 | 5.34 | 0.6735 | 6.687 |
| 5 | 128-64 | 1 | 256 | 0.3 | $1 \times 10^{-3}$ | 32 | 5.34 | 0.6527 | 6.645 |
| 6 | 64-32 | 1 | 256 | 0.3 | $1 \times 10^{-3}$ | 32 | 5.34 | 0.6457 | 6.631 |
| 7 | 64-32 | 1 | 128 | 0.3 | $3 \times 10^{-3}$ | 128 | 5.34 | 0.6432 | 6.626 |
| 8 | 128-64 | 1 | 64 | 0.3 | $3 \times 10^{-3}$ | 64 | 5.34 | 0.6306 | 6.601 |
| 9 | 128-64 | 1 | 128 | 0.4 | $3 \times 10^{-3}$ | 128 | 5.34 | 0.6276 | 6.595 |
| 10 | 128-64 | 1 | 128 | 0.5 | $1 \times 10^{-3}$ | 64 | 5.34 | 0.6256 | 6.591 |

Note. All models were trained for 40 epochs with batch size 64 and seed 42. A grid search was conducted over LSTM layers (1), LSTM hidden size (64, 128, 256), MLP head architectures (64–32, 128–64), dropout rates (0.3, 0.4, 0.5), learning rates ( $1 \times 10^{-4}$ , $3 \times 10^{-4}$ , $1 \times 10^{-3}$ , $3 \times 10^{-3}$ ), and embedding dimensions (32, 64, 128, 256). Rank 4 (MLP 128–64, LSTM hidden 64, Dp = 0.4, Lr = $1 \times 10^{-3}$ , Ed = 32) was selected over Ranks 1–3 for its more favourable train-validation loss profile (S10), indicating better generalisation despite a lower combined score on the Gram-negative sweep.

**FIG. S10.**
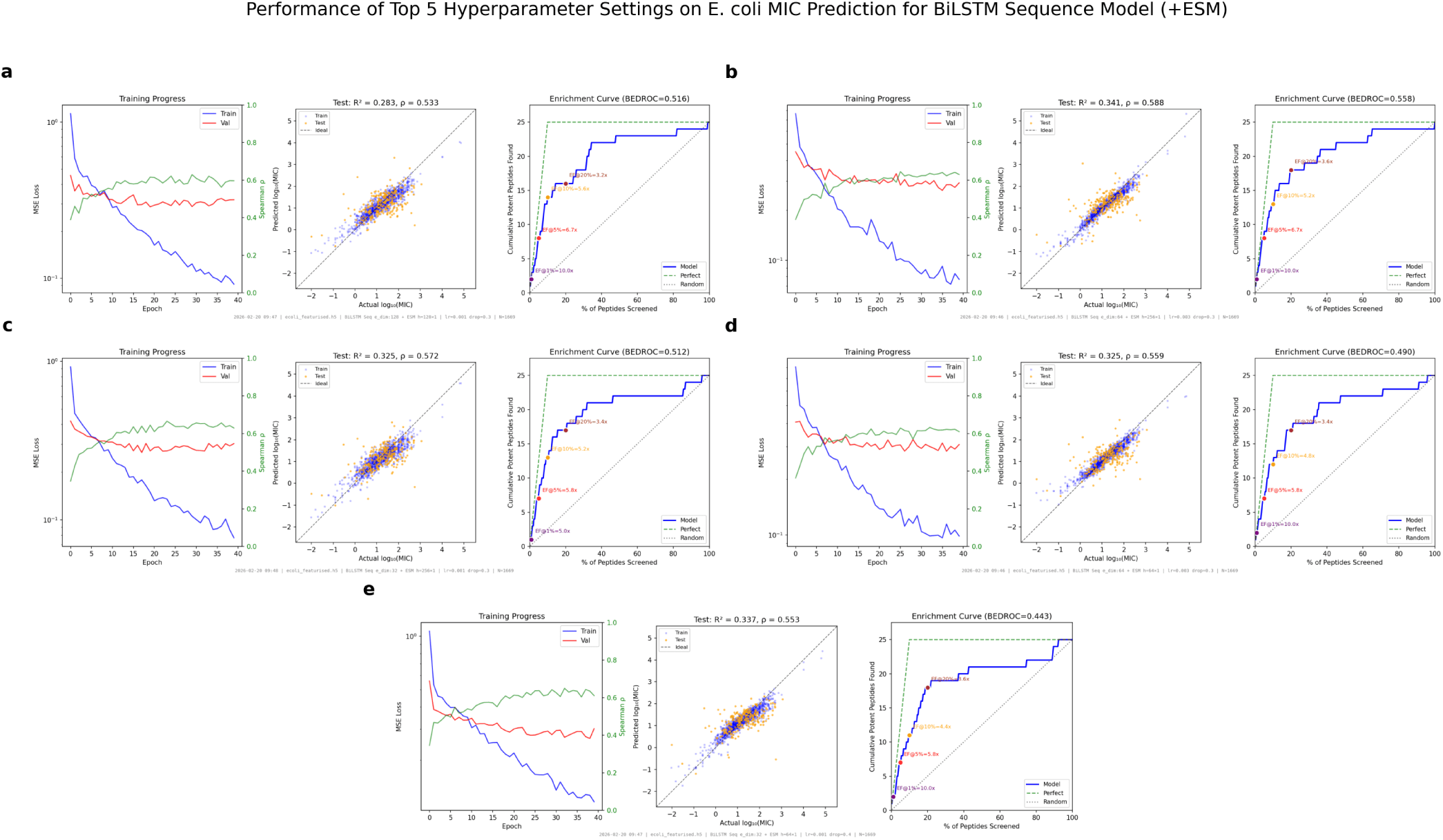
Performance of the top-5 BiLSTM Seq (+ESM) configurations — selected by combined score on the Gram-negative hyperparameter sweep — when retrained and evaluated on the *E. coli* peptide subset. Model (d), corresponding to rank 4 in the GN dataset, was selected as the final model using the evaluation framework mentioned in II A 5.

**Table S10.** Performance of the top-10 BiLSTM Seq (+ESM+QSAR) configurations hyperparameter configurations ranked by combined score on the Gram-negative test set. Dp = dropout; Lr = learning rate; Ed = embedding dimension; Score = EF@10% + 2*ρ*.

| Rank | MLP Arch. | LSTM Layers | LSTM Hidden | Dp | Lr | Ed | EF@10% | Spearman $\rho$ | Score |
| --- | --- | --- | --- | --- | --- | --- | --- | --- | --- |
| 1 | 128-64 | 1 | 128 | 0.4 | $3 \times 10^{-3}$ | 32 | 5.73 | 0.6394 | 7.009 |
| 2 | 64-32 | 1 | 256 | 0.5 | $3 \times 10^{-3}$ | 64 | 5.73 | 0.6242 | 6.978 |
| 3 | 128-64 | 1 | 256 | 0.5 | $3 \times 10^{-3}$ | 64 | 5.73 | 0.6107 | 6.951 |
| 4 | 64-32 | 1 | 256 | 0.3 | $3 \times 10^{-3}$ | 64 | 5.34 | 0.6772 | 6.694 |
| 5 | 128-64 | 1 | 256 | 0.4 | $3 \times 10^{-3}$ | 64 | 5.34 | 0.6675 | 6.675 |
| 6 | 128-64 | 1 | 256 | 0.3 | $3 \times 10^{-3}$ | 128 | 5.34 | 0.6392 | 6.618 |
| 7 | 128-64 | 1 | 256 | 0.3 | $3 \times 10^{-4}$ | 128 | 5.34 | 0.6382 | 6.616 |
| 8 | 64-32 | 1 | 128 | 0.3 | $3 \times 10^{-3}$ | 32 | 5.34 | 0.5934 | 6.527 |
| 9 | 128-64 | 1 | 64 | 0.5 | $3 \times 10^{-3}$ | 64 | 4.96 | 0.6674 | 6.295 |
| 10 | 64-32 | 1 | 64 | 0.4 | $3 \times 10^{-3}$ | 64 | 4.96 | 0.6642 | 6.288 |

Note. All models were trained for 40 epochs with batch size 64 and seed 42. A grid search was conducted over LSTM layers (1), LSTM hidden size (64, 128, 256), MLP head architectures (64–32, 128–64), dropout rates (0.3, 0.4, 0.5), learning rates ( $1 \times 10^{-4}$ , $3 \times 10^{-4}$ , $1 \times 10^{-3}$ , $3 \times 10^{-3}$ ), and embedding dimensions (32, 64, 128, 256). Rank 1 (MLP 128–64, LSTM hidden 128, Dp = 0.4, Lr = $3 \times 10^{-3}$ , Ed = 32) was selected as the final BiLSTM Seq (+ESM+QSAR) configuration.

**FIG. S11.**
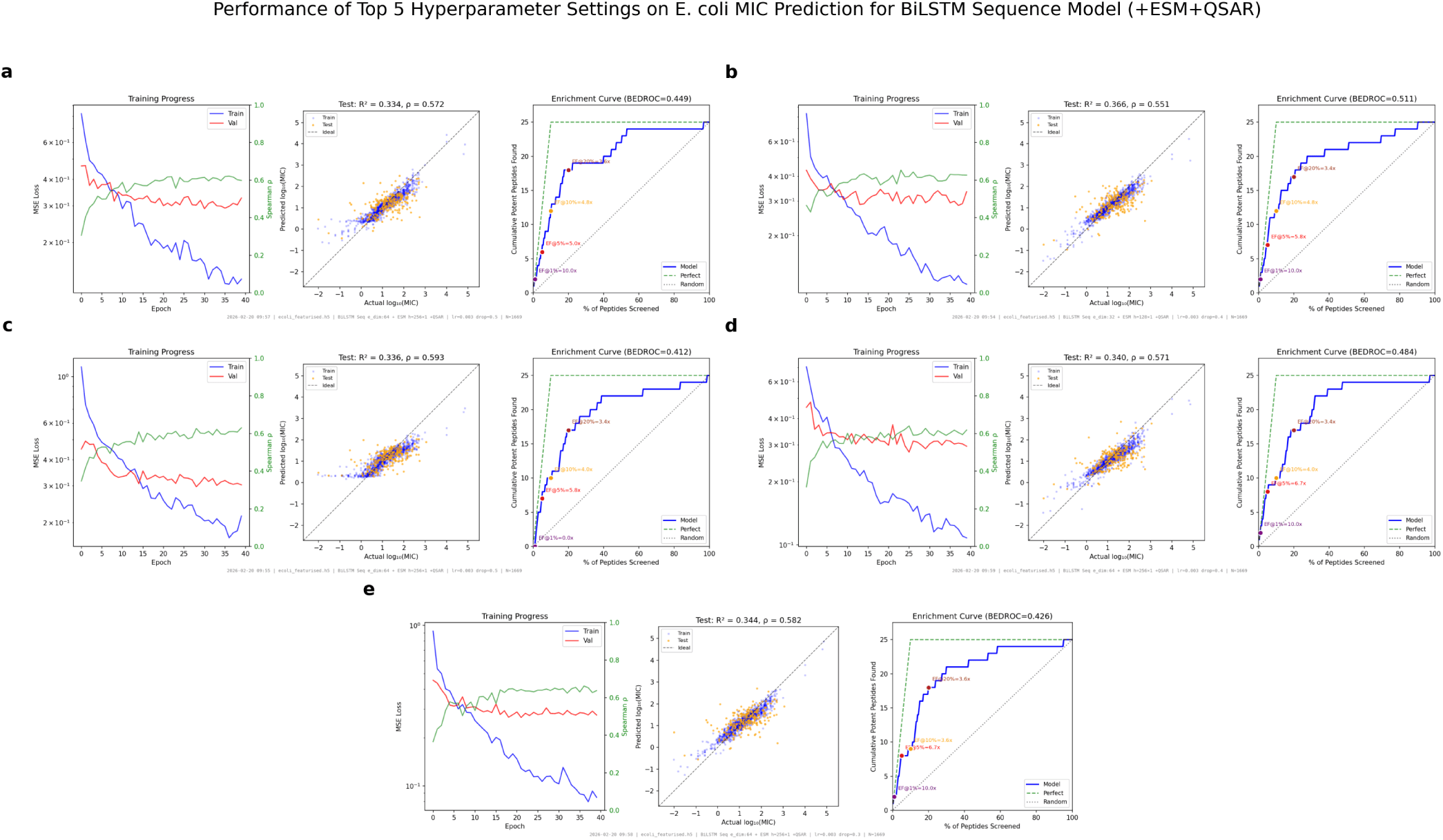
Performance of the top-5 BiLSTM Seq (+ESM+QSAR) hyperparameter configurations ranked by combined score on the Gram-negative test set. Model (d), corresponding to rank 4 in the GN dataset, was selected as the final model using the evaluation framework mentioned in II A 5.

**Table S11.** Hyperparameters used for the selected MLP models in MIC prediction.

| Model | Architecture | Dropout | Learning Rate | Epochs |
| --- | --- | --- | --- | --- |
| MLP (ESM+QSAR) | 128-64 | 0.5 | 0.001 | 40 |
| MLP (ESM-only) | 128-64 | 0.4 | 0.003 | 45 |

**Table S12.** Hyperparameters used for the selected BiLSTM models in MIC prediction.

| Model | MLP Architecture | LSTM layers | LSTM Hidden Layers | Dropout | Learning Rate | Embedding Dimension | Epochs |
| --- | --- | --- | --- | --- | --- | --- | --- |
| BiLSTM (+QSAR) | 64-32 | 1 | 128 | 0.3 | 0.0003 | - | 40 |
| BiLSTM | 64-32 | 1 | 256 | 0.5 | 0.0003 | - | 40 |
| BiLSTM Seq | 64-32 | 1 | 256 | 0.5 | 0.0003 | 64 | 40 |
| BiLSTM Seq (+QSAR) | 64-32 | 1 | 128 | 0.5 | 0.003 | 64 | 40 |
| BiLSTM Seq (+ESM) | 128-64 | 1 | 64 | 0.4 | 0.001 | 32 | 40 |
| BiLSTM Seq (+ESM+ QSAR) | 128-64 | 1 | 128 | 0.4 | 0.003 | 32 | 40 |

**Table S13.** Comparison of selected model performance across Gram-negative (GN) and *E. coli* datasets using multiple evaluation metrics. Score = EF@10% + 2*ρ*. Rankings are within-dataset.

| Model | Rank | | EF@10% | | Spearman $\rho$ | | Score | |
| --- | --- | --- | --- | --- | --- | --- | --- | --- |
|  | GN | *E. coli* | GN | *E. coli* | GN | *E. coli* | GN | *E. coli* |
| MLP (ESM+QSAR) | 2 | 2 | 5.73 | 5.20 | 0.6398 | 0.5812 | 7.010 | 6.362 |
| MLP (ESM-only) | 1 | 5 | 5.73 | 4.00 | 0.6184 | 0.5602 | 6.967 | 5.120 |
| BiLSTM | 1 | 3 | 5.73 | 4.80 | 0.6177 | 0.5739 | 6.965 | 5.948 |
| BiLSTM (+QSAR) | 1 | 4 | 5.34 | 4.80 | 0.6475 | 0.5569 | 6.635 | 5.914 |
| BiLSTM Seq | 3 | 3 | 6.11 | 3.60 | 0.5467 | 0.5504 | 7.203 | 4.701 |
| BiLSTM Seq (+QSAR) | 3 | 4 | 5.73 | 3.60 | 0.6187 | 0.5413 | 6.967 | 4.683 |
| BiLSTM Seq (+ESM) | 4 | 4 | 5.34 | 4.00 | 0.6735 | 0.5856 | 6.687 | 5.171 |
| BiLSTM Seq (+ESM+QSAR) | 2 | 1 | 5.73 | 4.80 | 0.6394 | 0.5489 | 7.009 | 5.898 |

*Note.* Each model's selected configuration was determined by hyperparameter sweep on the GN dataset and subsequently retrained and evaluated on the *E. coli* subset. The consistent drop in EF@10% and Spearman $\rho$ from GN to *E. coli* reflects the reduced dataset size of the species-specific subset rather than a failure of generalisation, as discussed in Section S3 A 1.

**Table S14.** Comparison of EF@10%, Spearman *ρ*, and combined score for Huber loss with *δ* = 0.5 and *δ* = 1.0, evaluated on the *E. coli* test set using the fixed MLP (ESM+QSAR) configuration (architecture 128–64, dropout = 0.5, lr = 1 × 10^−3^).

| Huber $\delta$ | EF@10% | Spearman $\rho$ | Score |
| --- | --- | --- | --- |
| 0.5 | 5.34 | 0.6273 | 6.595 |
| 1.0 | 5.34 | 0.6345 | 6.609 |

*Note.* Neither Huber loss variant consistently outperformed MSE (Score = $6.095 \pm 0.289$ ; Table S1), and MSE was therefore retained as the training objective given its stronger penalisation of large deviations, which is directly relevant to identifying highly active peptides at the extremes of the MIC distribution.

**Table S15.** Effect of weight decay on MLP (ESM+QSAR) performance evaluated on the *E. coli* test set. Results are reported for the fixed configuration (architecture 128–64, dropout = 0.5, lr = 1 × 10^−3^). Score = EF@10% + 2*ρ*.

| Weight Decay | EF@10% | Spearman $\rho$ | Score |
| --- | --- | --- | --- |
| $1 \times 10^{-4}$ | 5.73 | 0.6398 | 7.010 |
| 0 | 5.73 | 0.6294 | 6.989 |
| $1 \times 10^{-5}$ | 5.73 | 0.6294 | 6.989 |
| $5 \times 10^{-4}$ | 5.34 | 0.6328 | 6.606 |
| $1 \times 10^{-3}$ | 5.34 | 0.6308 | 6.602 |
| $5 \times 10^{-3}$ | 4.96 | 0.6255 | 6.211 |

*Note.* A weight decay of $1 \times 10^{-4}$ yielded the best overall performance, achieving the highest Spearman rank correlation and combined score while maintaining strong early enrichment. This value corresponds to the standard default for the AdamW optimiser and provides an effective balance between regularisation and model capacity for the given dataset and architecture<sup>S22</sup>.

**FIG. S12.**
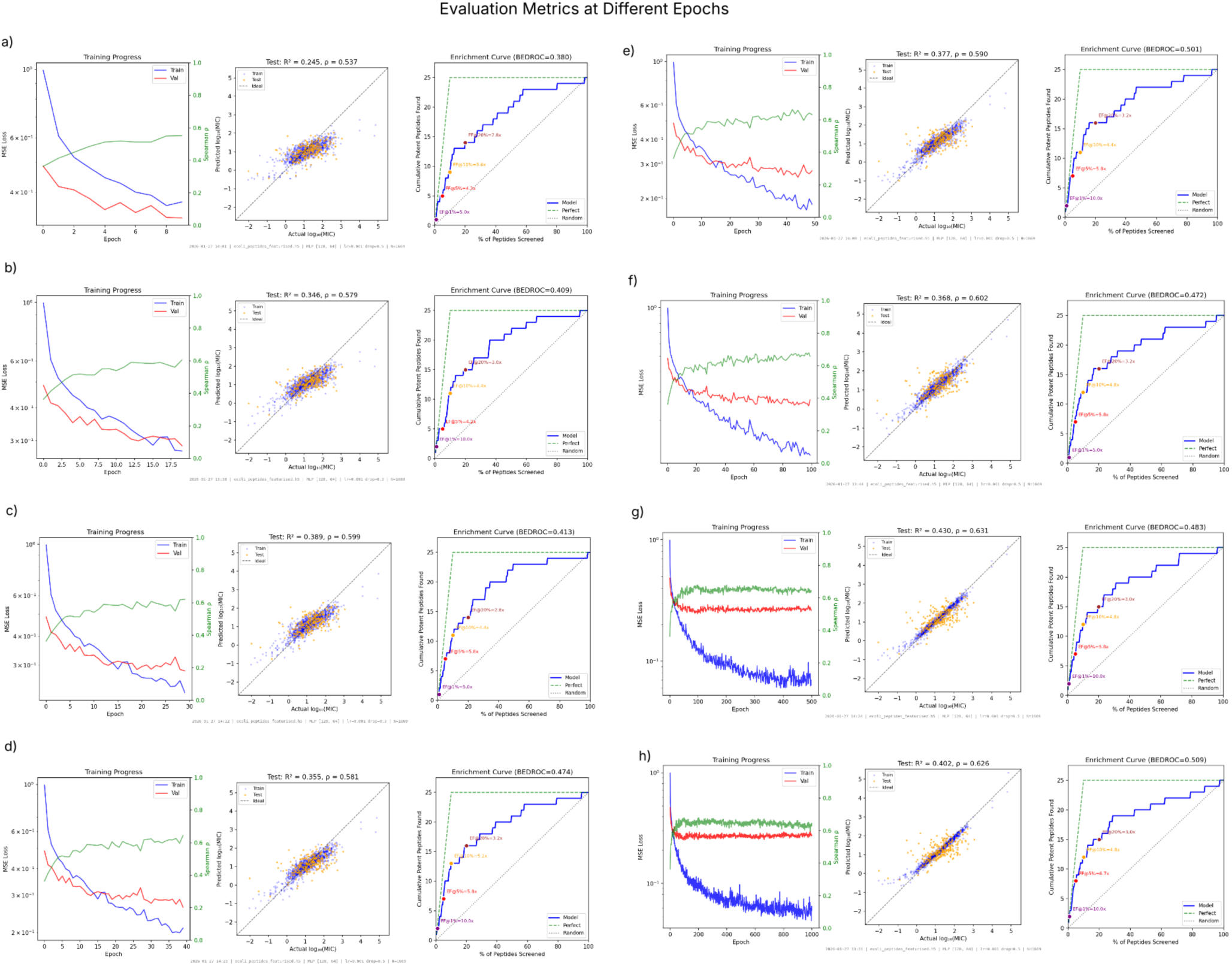
Evolution of evaluation metrics across training epochs for the MLP (ESM+QSAR) model (architecture 128–64, dropout = 0.5, lr = 1 × 10^−3^) trained on the *E. coli* dataset. Panels (a)–(h) correspond to epochs 10, 20, 30, 40, 50, 100, 500, and 1000, respectively. Each panel shows training and validation loss curves, predicted versus true log_10_(MIC) values, and cumulative enrichment, illustrating convergence behaviour, generalisation trends, and ranking stability over extended training.

**Table S16.** Comparison of EF@10%, Spearman *ρ*, and combined score across training epochs for the selected MLP (ESM+QSAR) configuration (architecture 128–64, dropout = 0.5, lr = 1 × 10^−3^) on the *E. coli* dataset. Early stopping was triggered at epoch 12. Score = EF@10% + 2*ρ*.

| Epochs | EF@10% | Spearman $\rho$ | Score |
| --- | --- | --- | --- |
| 10 | 3.60 | 0.537 | 4.674 |
| 20 | 4.40 | 0.579 | 5.558 |
| 30 | 4.40 | 0.599 | 5.599 |
| 40 | 5.20 | 0.581 | 6.362 |
| 50 | 4.40 | 0.590 | 5.580 |
| 100 | 4.80 | 0.601 | 6.003 |
| 500 | 4.80 | 0.631 | 6.063 |
| 1000 | 4.80 | 0.626 | 6.052 |

*Note.* Controlled extension of training beyond the early stopping trigger at epoch 12 yielded improved test set performance, with epoch 40 achieving the highest combined score (6.362). These findings suggest that strict reliance on early stopping may lead to undertraining on small, noisy biological datasets, and that limited additional training can improve model robustness and ranking performance. Performance stabilises beyond epoch 100, with diminishing returns at epochs 500 and 1000, confirming that extended training does not introduce progressive overfitting under this configuration.

#### C. Distribution Analysis

**FIG. S13.**
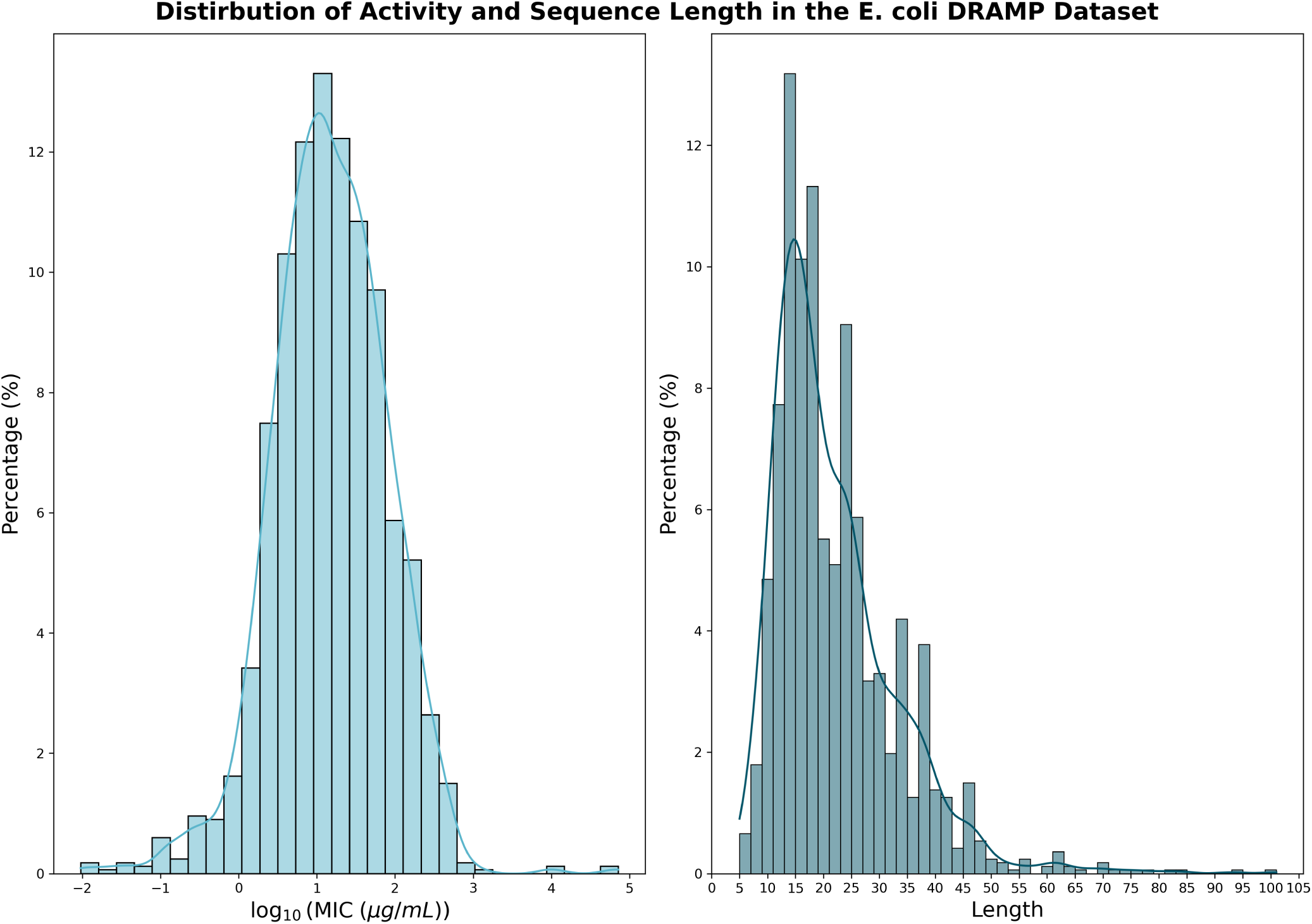
Distribution of antimicrobial activity and sequence length in the *E. coli* subset of the DRAMP dataset. Left: Histogram of log_10_-transformed minimum inhibitory concentration (MIC, [µg mL^−1^]), showing an approximately normal distribution centred around moderate activity values. Right: Distribution of peptide sequence lengths, which is right-skewed, with most peptides between 10 and 30 residues but with a long tail extending to longer sequences. Kernel density estimates (KDE) are overlaid to highlight the underlying distribution shapes.

**FIG. S14.**
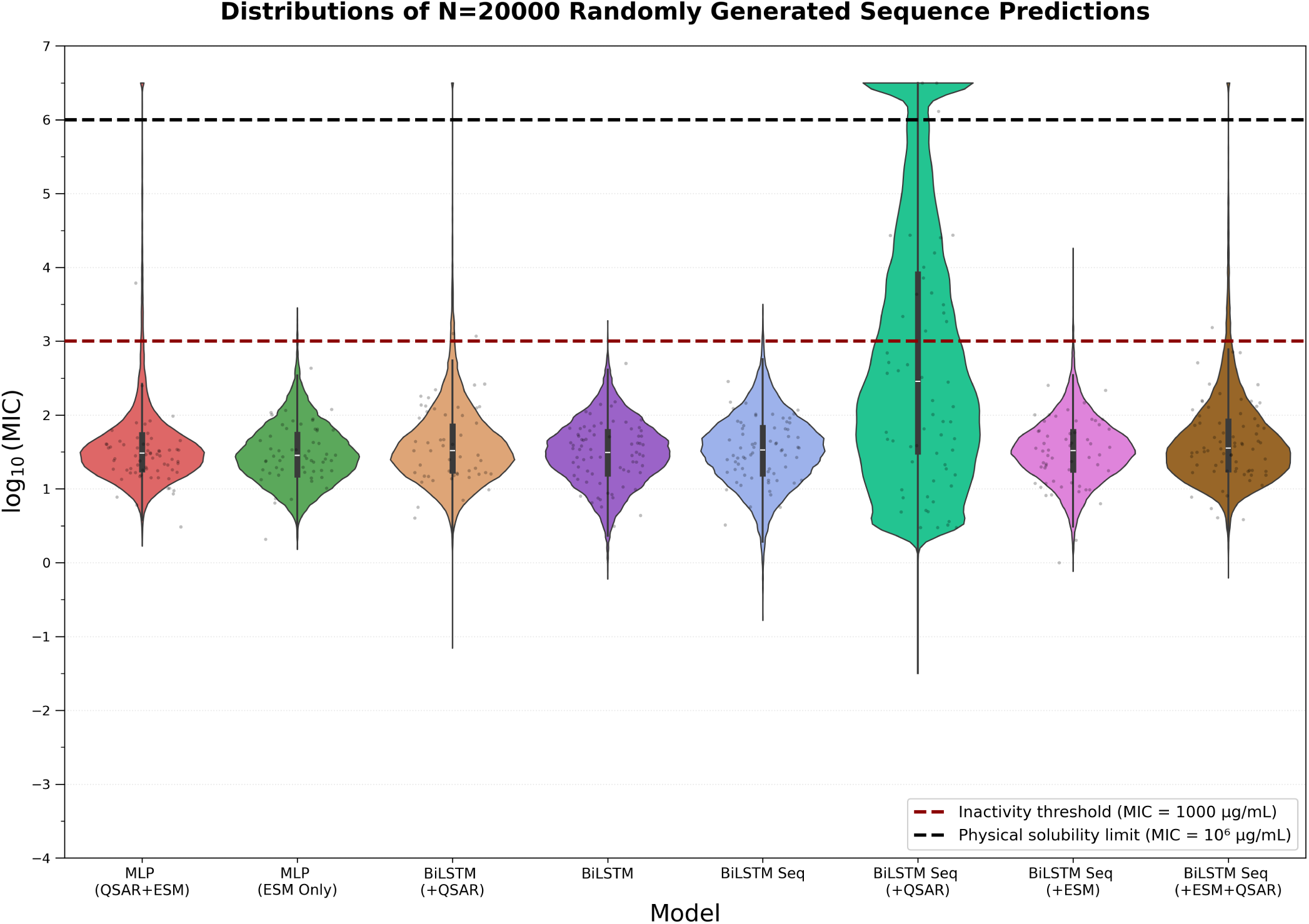
Predicted log_10_(MIC) [µg mL^−1^] distribution for 20,000 randomly generated peptides. Individual points represent a random sample of 500 selected peptides per model for visualisation clarity. The red dashed line denotes the inactivity threshold (log_10_(MIC) = 3 µg mL^−1^), above which antimicrobial activity is considered practically irrelevant. The black dashed line indicates the physical solubility limit (log_10_(MIC) = 6 µg mL^−1^), beyond which the required peptide mass exceeds the solution volume.

**FIG. S15.**
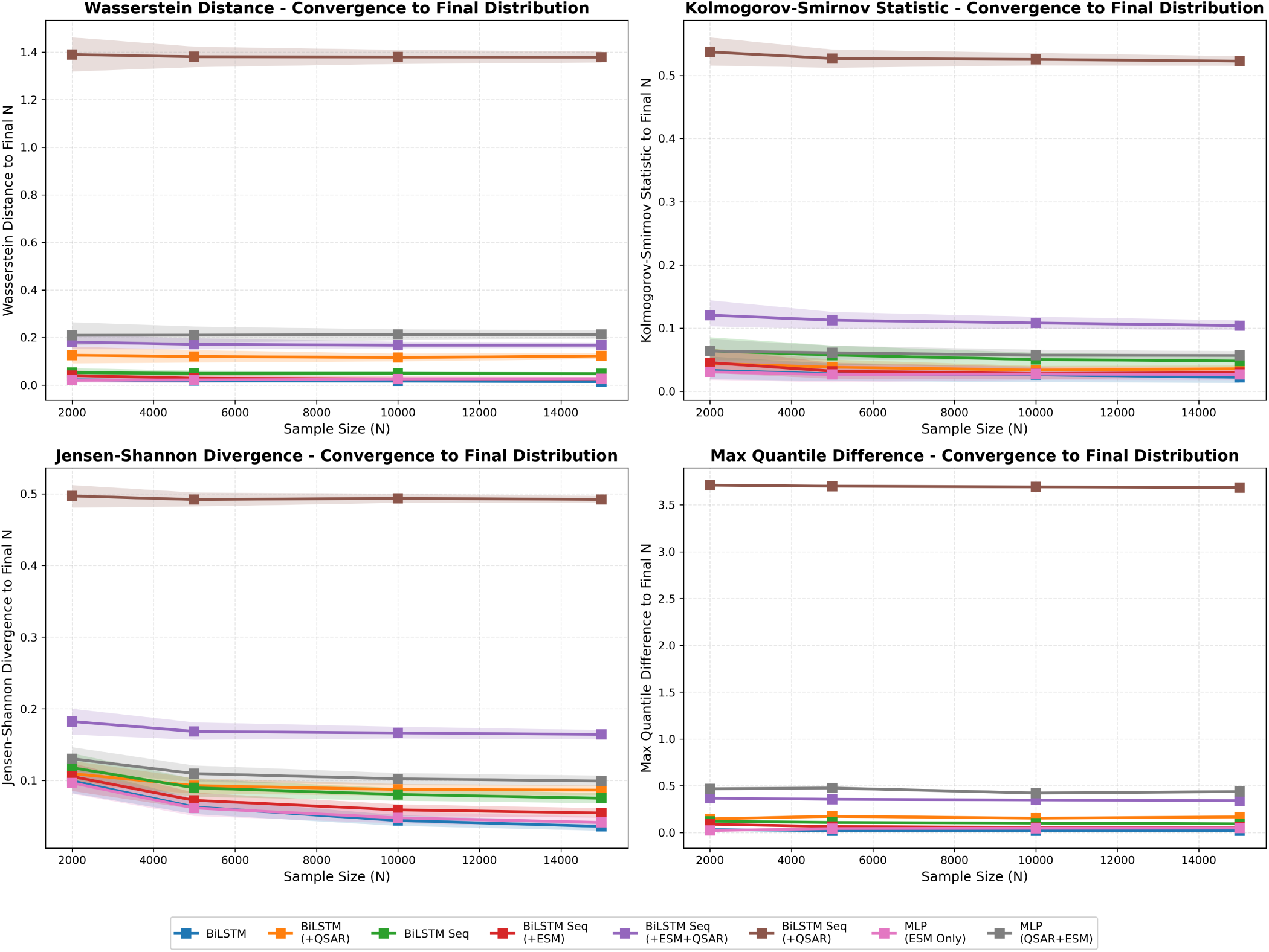
Step-wise convergence analysis of ensemble MIC predictions over randomly generated peptide pools of increasing size (2,000–20,000 samples). Four distributional distance metrics — Wasserstein distance, Kolmogorov–Smirnov statistic, Jensen–Shannon divergence, and maximum quantile difference — were computed between successive sample sizes for each model. The BiLSTM Seq model was not included as it was identified as an outlier. Shaded bands denote 95% bootstrap confidence intervals. Red dashed lines indicate predefined stabilisation thresholds. All models converge below threshold by approximately 10,000 samples, confirming that a generative pool of 40,000 peptides lies well within the stable regime.

**Table S17.** Top 5 highest predicted log_10_(MIC) [µg mL^−1^] values among 20,000 randomly generated peptides.

| Rank | Sequence | Model | $\log_{10}(\text{MIC})$ | Std |
| --- | --- | --- | --- | --- |
| 1 | FIFMI | MLP (ESM+QSAR) | 60.99 | 2.827 |
| 2 | MVFMI | MLP (ESM+QSAR) | 57.00 | 2.830 |
| 3 | MFIYI | MLP (ESM+QSAR) | 47.17 | 21.02 |
| 4 | HNQEE | MLP (ESM+QSAR) | 42.46 | 12.83 |
| 5 | MVFMI | BiLSTM Seq (+ESM+QSAR) | 38.40 | 4.773 |

**Table S18.**
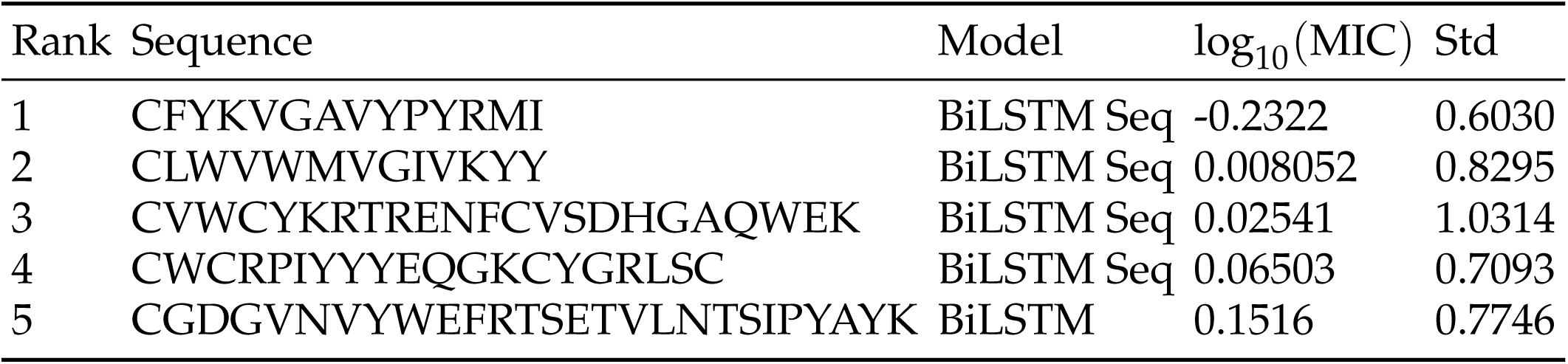
Top 5 lowest predicted log_10_(MIC) [µg mL^−1^] values among 20,000 randomly generated peptides.

**FIG. S16.**
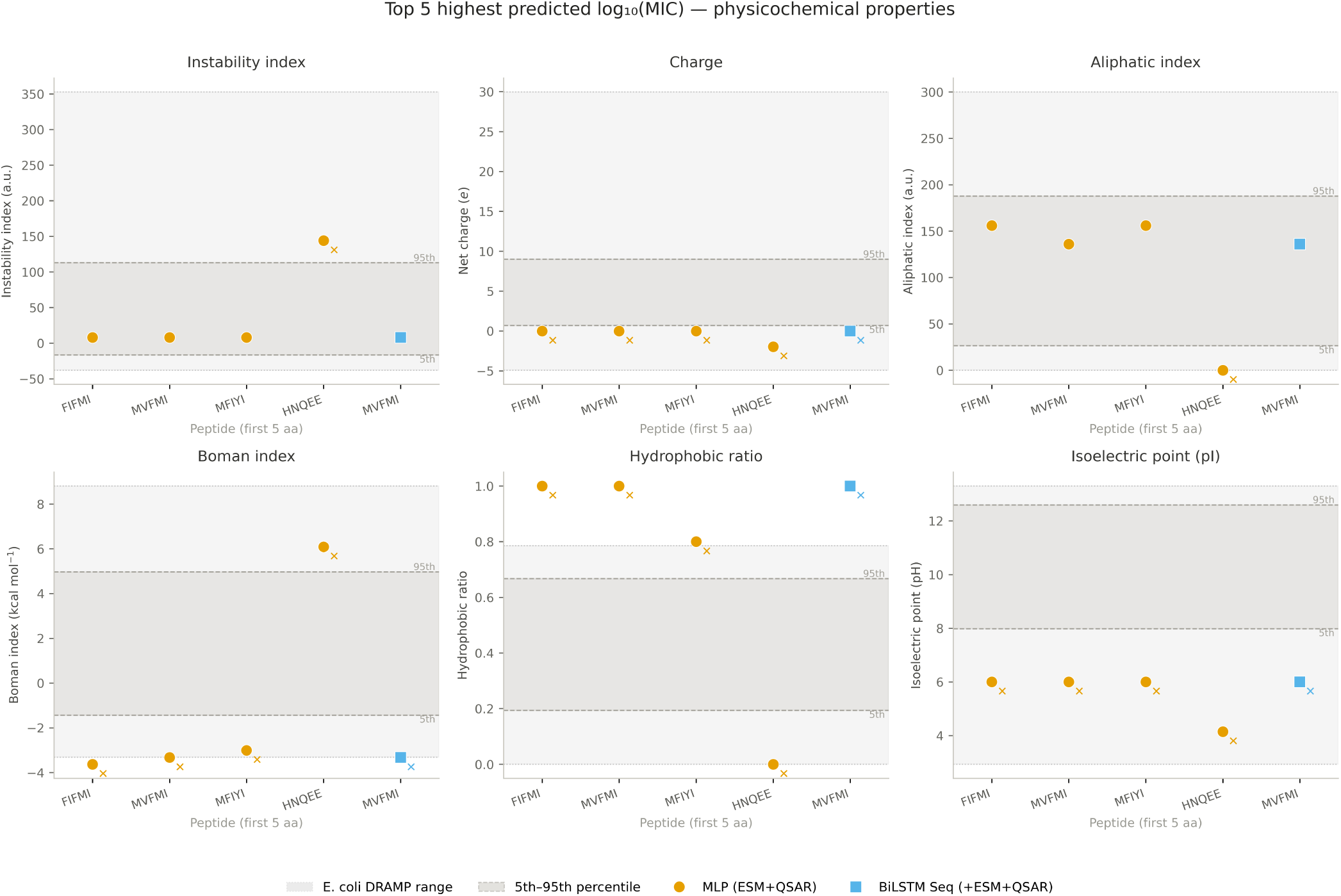
Physicochemical properties of the five peptides with the highest predicted log_10_(MIC) [µg mL^−1^] values. For each property (instability index, charge, aliphatic index, Boman index, hydrophobic ratio, and isoelectric point), individual peptide values are shown. Orange circles denote peptide predictions made by the MLP (ESM+QSAR) model, and blue squares denote peptide predictions made by the BiLSTM Seq (+ESM+QSAR) model. The shaded region indicates the *E. coli* DRAMP property range, while dashed horizontal lines mark the *E. coli* DRAMP 5th–95th percentile bounds. Crosses indicate values that fall outside the *E. coli* DRAMP 5th–95th percentile range.

**FIG. S17.**
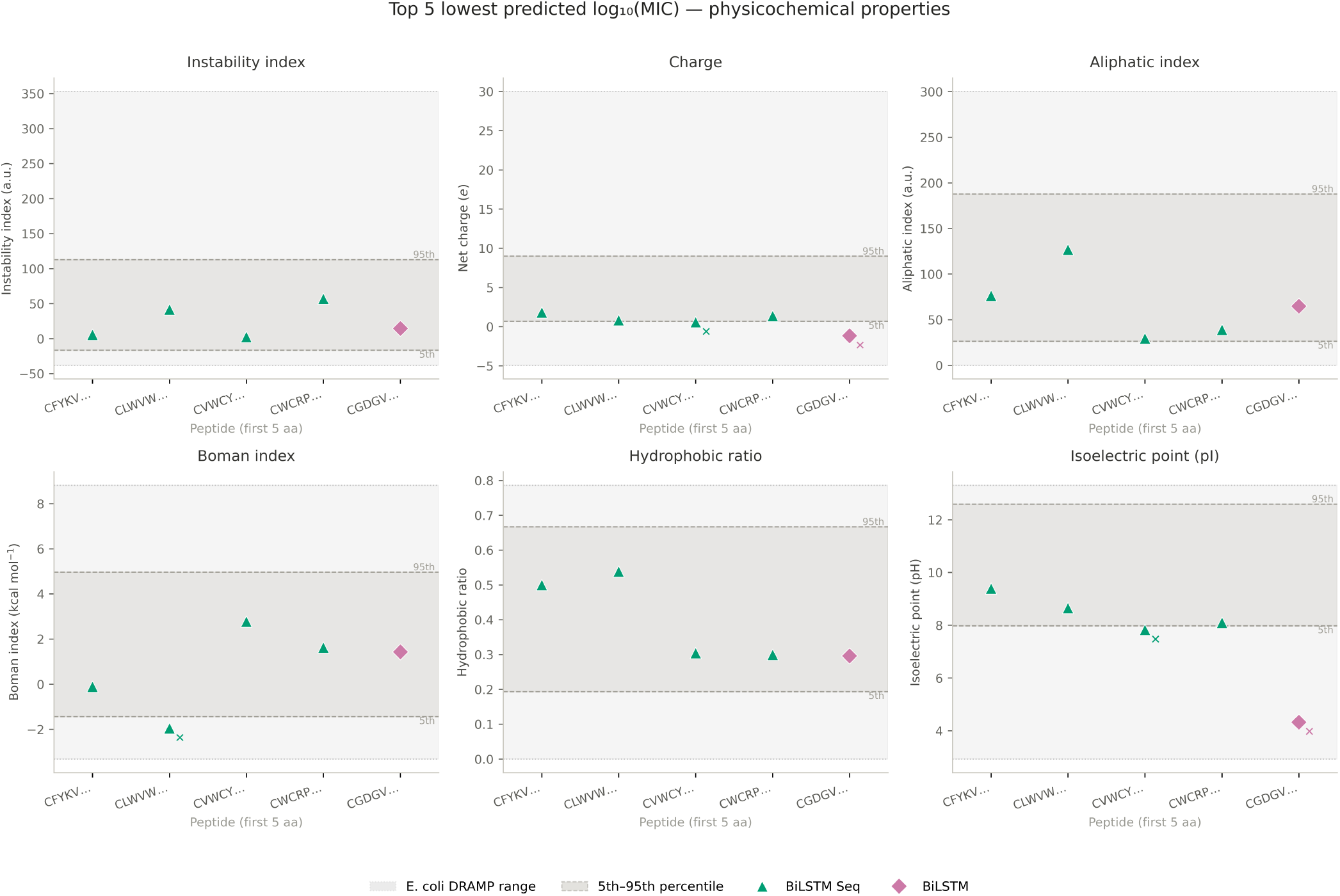
Physicochemical properties of the five peptides with the lowest predicted log_10_(MIC) [µg mL^−1^] values. For each property (instability index, charge, aliphatic index, Boman index, hydrophobic ratio, and isoelectric point), individual peptide values are shown. Green triangles denote peptide predictions made by BiLSTM Seq model, and pink diamonds denote peptide predictions made by the BiLSTM model. The shaded region indicates the *E. coli* DRAMP property range, while dashed horizontal lines mark the *E. coli* DRAMP 5th–95th percentile bounds. Crosses indicate values that fall outside the *E. coli* DRAMP 5th–95th percentile range.

**FIG. S18.**
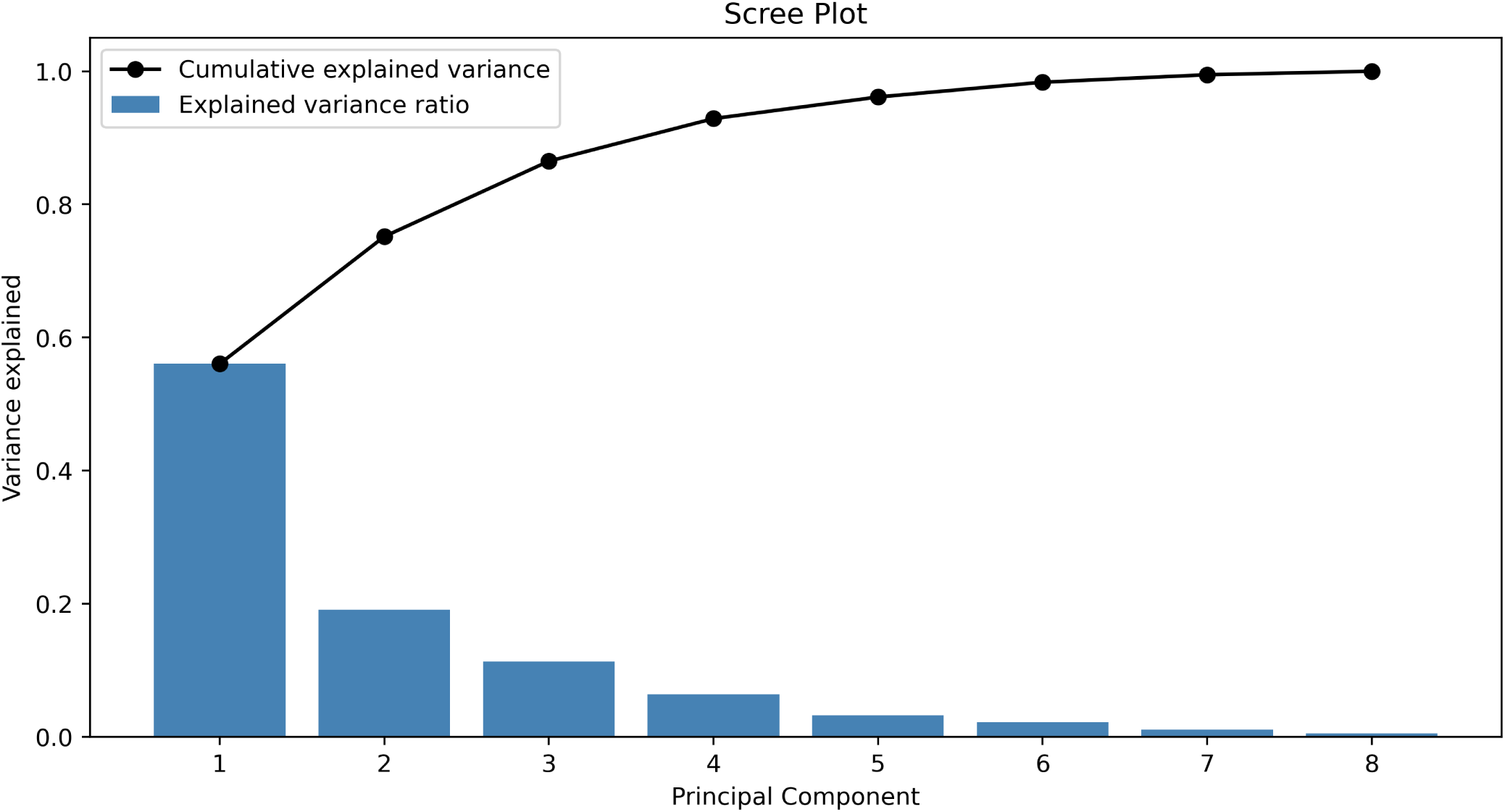
Scree plot, showing the proportion of variance explained by each principal component (blue bars) and the cumulative explained variance (black line) for the eight model prediction outputs for the *E. coli* test and 20k randomly-generated peptide dataset. PC1 explains approximately 54% of the total variance, with the first three components accounting for over 80% of the variance. The gradual flattening of the curve after PC3 indicates diminishing marginal contribution of additional components, suggesting that most of the variability in model predictions is captured by the first few principal components.

#### D. PCA Analysis

**FIG. S19.**
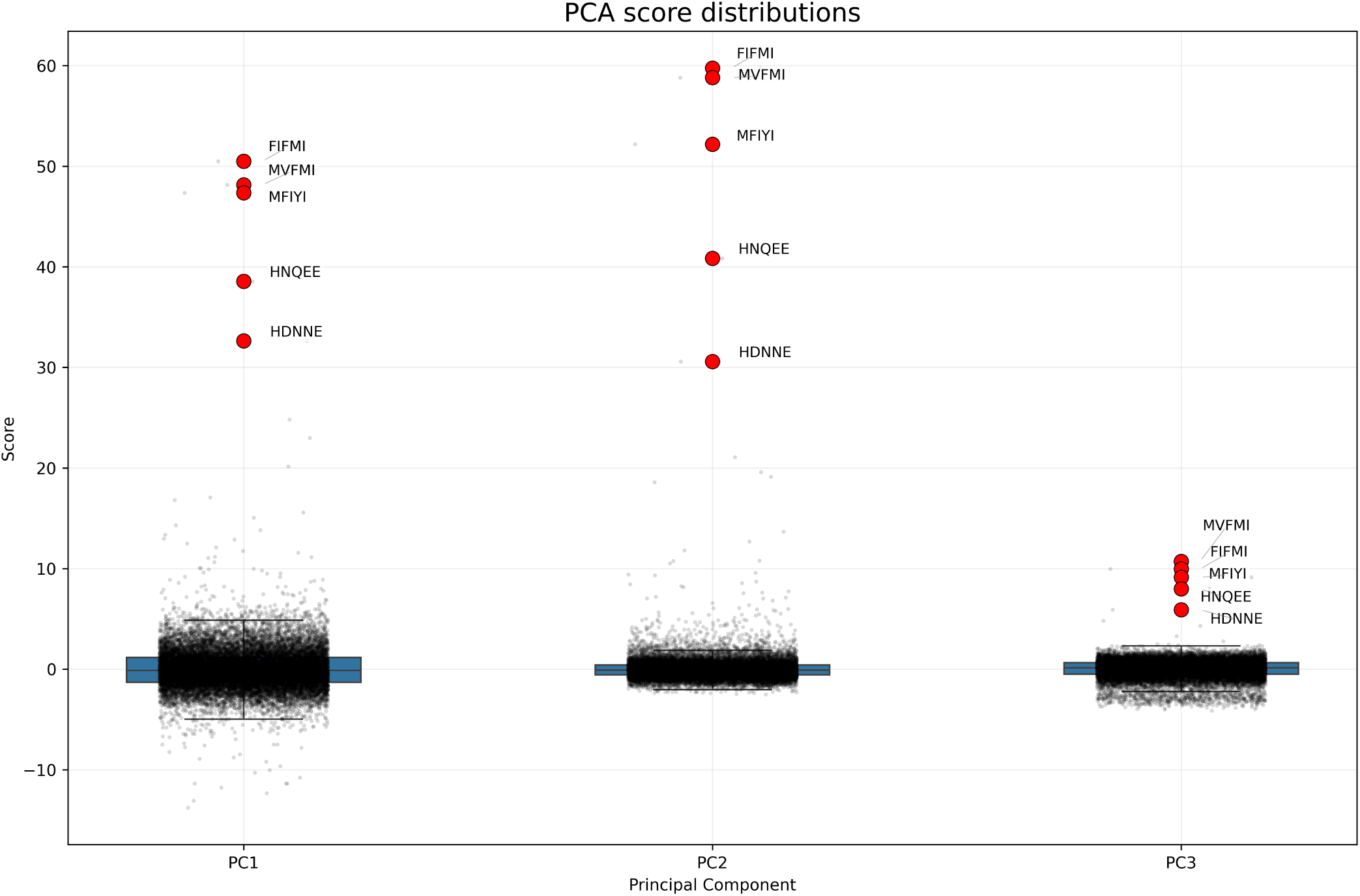
Distribution of principal component (PC) scores for generated peptides.Boxplots show the distribution of scores along PC1, PC2, and PC3 obtained from PCA of model prediction outputs. Individual points denote outlier peptides, with the top 5 extreme outliers for each PC highlighted in red and annotated by sequence. Extreme outliers were selected by ranking the PC of each peptide by its absolute score and these were selected to be marked on the PCA plot (Figure S20). The majority of peptides cluster tightly around zero across all components, while a small subset exhibits markedly high positive PC scores, indicating distinct prediction patterns relative to the main population

**FIG. S20.**
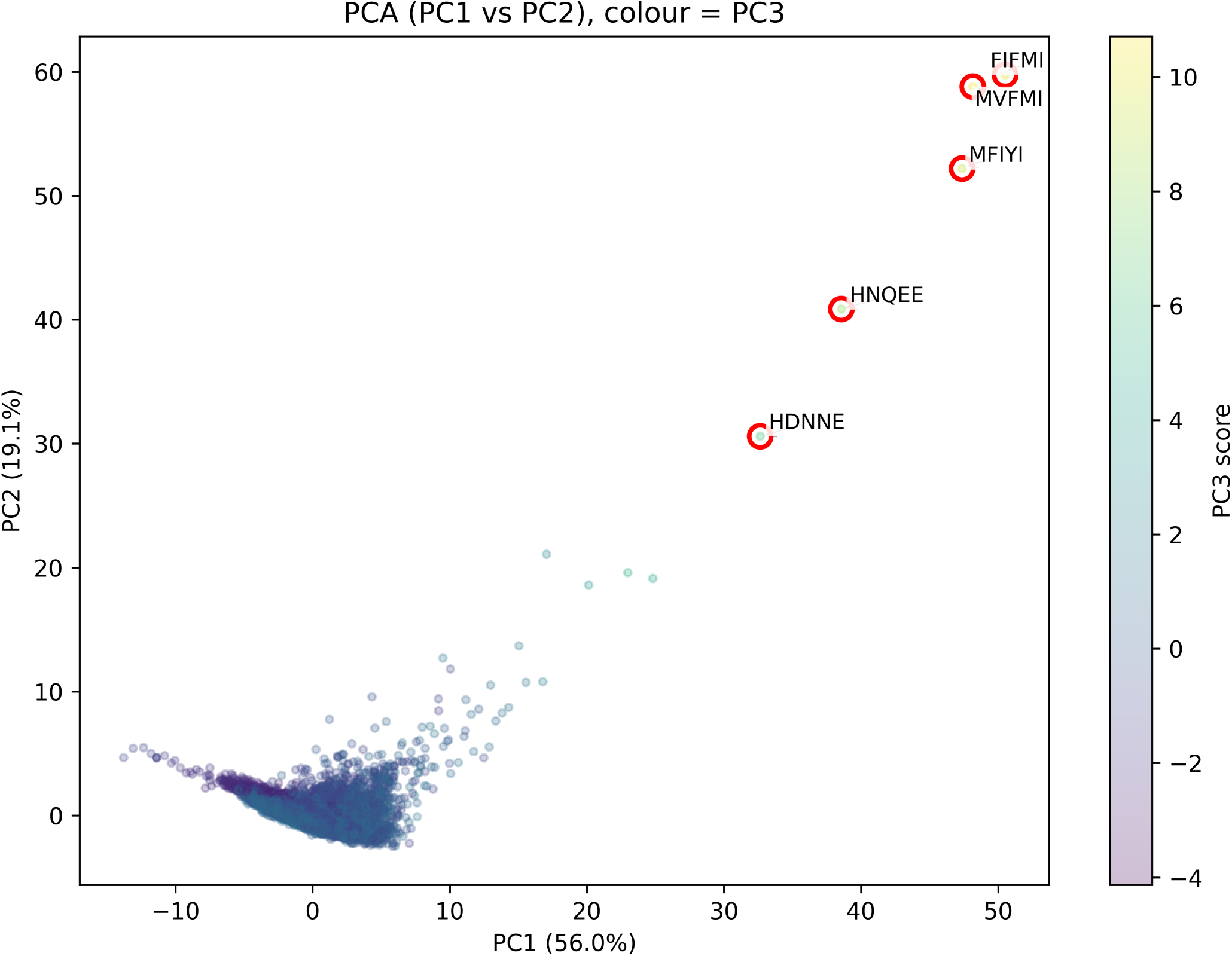
Principal component analysis (PCA) of MIC prediction profiles across peptides. PC1 and PC2 explain 50.0% and 19.3% of the variance, respectively, and points are coloured by PC3 score. Red-circled points indicate peptides with extreme prediction profiles relative to the main cluster.

#### E. Ensemble Models

**Table S19.** Pairwise ensemble weights across random seeds for the MLP (ESM+QSAR) + BiLSTM combination. Weights correspond to the contribution of MLP (ESM+QSAR) in the pairwise linear ensemble optimised on the validation set; the complementary weight (1 − *w*) represents the BiLSTM contribution.

| Condition | MLP (ESM+QSAR) Weight |
| --- | --- |
| Seed 42 | 0.5663 |
| Seed 77 | 0.8139 |
| Seed 123 | 0.6044 |
| Mean ± std | $0.6616 \pm 0.1335$ |

**Table S20.** NNLS ensemble weights across three independent random seeds. Non-negative least squares (NNLS) weights were fitted on the validation set for each seed and normalised to sum to one. Zero weights indicate that a model did not contribute to the optimal validation solution for that seed.

| Model | Seed 42 | Seed 77 | Seed 123 |
| --- | --- | --- | --- |
| MLP (ESM+QSAR) | 0.2622 | 0.4435 | 0.1707 |
| BiLSTM Seq (+ESM) | 0.2351 | 0.5217 | 0.0000 |
| BiLSTM | 0.2295 | 0.0000 | 0.2264 |
| BiLSTM Seq (+ESM+QSAR) | 0.1438 | 0.0000 | 0.0000 |
| BiLSTM Seq (+QSAR) | 0.1294 | 0.0000 | 0.0000 |
| MLP (ESM Only) | 0.0000 | 0.0347 | 0.3390 |
| BiLSTM (+QSAR) | 0.0000 | 0.0000 | 0.0977 |
| BiLSTM Seq | 0.0000 | 0.0000 | 0.1243 |
| **Sum** | **1.0000** | **1.0000** | **1.0000** |

*Note.* The composition of models in the NNLS ensemble varied across random seeds. However, MLP (ESM+QSAR) was consistently assigned a non-zero weight across all three seeds, indicating its importance as a key contributor to predictive performance. Each ensemble relied on only a small subset of the available models, reflecting the sparsity-inducing property of NNLS optimisation.

**Table S21.** Comparison of ensemble methods to the MLP (ESM+QSAR).

| Method | Dataset | $R^2$ | MAE | RMSE | Spearman $\rho$ | BEDROC ($\alpha = 20$) | EF@10% | Combined Score |
| --- | --- | --- | --- | --- | --- | --- | --- | --- |
| MLP (ESM+QSAR) | Val | $0.441 \pm 0.038$ | $0.401 \pm 0.012$ | $0.529 \pm 0.017$ | $0.641 \pm 0.004$ | $0.556 \pm 0.056$ | $4.76 \pm 0.22$ | $6.040 \pm 0.218$ |
| | Test | $0.365 \pm 0.132$ | $0.426 \pm 0.024$ | $0.599 \pm 0.098$ | $0.639 \pm 0.050$ | $0.561 \pm 0.077$ | $4.82 \pm 0.38$ | $6.095 \pm 0.289$ |
| Equal weights | Val | $0.460 \pm 0.046$ | $0.388 \pm 0.010$ | $0.520 \pm 0.020$ | $0.661 \pm 0.020$ | $0.605 \pm 0.057$ | $5.38 \pm 0.36$ | $6.705 \pm 0.369$ |
| | Test | $0.461 \pm 0.084$ | $0.403 \pm 0.024$ | $0.553 \pm 0.064$ | $0.672 \pm 0.049$ | $0.595 \pm 0.096$ | $5.14 \pm 0.54$ | $6.487 \pm 0.541$ |
| Pairwise MLP (ESM+QSAR) + BiLSTM | Val | $0.460 \pm 0.032$ | $0.389 \pm 0.011$ | $0.520 \pm 0.014$ | $0.657 \pm 0.013$ | $0.608 \pm 0.056$ | $5.46 \pm 0.47$ | $6.779 \pm 0.492$ |
| | Test | $0.416 \pm 0.073$ | $0.417 \pm 0.014$ | $0.575 \pm 0.054$ | $0.653 \pm 0.035$ | $0.604 \pm 0.082$ | $5.27 \pm 0.62$ | $6.577 \pm 0.628$ |
| NNLS ensemble | Val | $0.484 \pm 0.041$ | $0.380 \pm 0.007$ | $0.508 \pm 0.019$ | $0.678 \pm 0.008$ | $0.620 \pm 0.058$ | $5.60 \pm 0.28$ | $6.953 \pm 0.294$ |
| | Test | $0.470 \pm 0.091$ | $0.398 \pm 0.021$ | $0.547 \pm 0.063$ | $0.676 \pm 0.061$ | $0.613 \pm 0.092$ | $5.30 \pm 0.87$ | $6.652 \pm 1.053$ |

*Note.* The MLP (ESM+QSAR) was selected for comparison as it was the highest-performing single model. Mean ± standard deviation performance across three random seeds is reported for individual and ensemble MIC prediction models on the *E. coli* validation and test sets.

### S4. GENERATIVE MODELS

#### A. Early development

##### 1. Shakespeare Benchmark: LSTM vs. GPT

The Shakespeare corpus benchmark confirmed that both models trained stably under the shared configuration, reaching a clear held-out optimum (Figures S22 and S23). This validated the transferred hyperparameter settings were viable rather than poorly calibrated, before exposure to peptide data. Under matched conditions, the LSTM achieved lower best training loss and reached its best test loss far earlier, but its best test loss was substantially worse than the GPT’s (Figures S30). The widening train–test divergence beyond the LSTM’s optimal epoch indicated aggressive memorisation rather than generalisation. The GPT continued to improve for longer and converged to a lower held-out optimum, consistent with transformer-based models handling longer-range dependencies more robustly under consistent conditions. Per-sequence perplexity analysis—quantified by minimum Levenshtein distance to the Shakespeare vocabulary—was conducted (Figure S27). With the near-distribution British English dataset a gradual increase in the model’s uncertainty was observed, demonstrating that the model learned transferable orthographic and morphological patterns rather than simply memorising the training text. Instead, model uncertainty for the randomly-generated words was substantially higher, with median log-perplexity of 460 compared to 12–17 for the Shakespeare and British English datasets (Figure S26). The analogy to peptide modelling is direct: an effective generative model must capture residue co-occurrence patterns, motif-like subsequences, and valid termination behaviour while still assigning higher uncertainty to implausible out-of-distribution sequences.

##### 2. DBAASP Training and Hyperparameter Adaptation

Once it was confirmed on the Shakespeare corpus that the models produced coherent outputs and learned meaningful sequence structure, architecture-level hyperparameters were retained for training on DBAASP. By contrast, optimisation settings that are more sensitive to dataset size and noise, such as learning rate, dropout, and batch size, were re-evaluated. Specifically, dropout regularisation was introduced to prevent overfitting, particularly relevant in low-data regimes. A moderate dropout value of 0.1 was selected as it is commonly used in Transformer-based models to provide regularisation without significantly impairing learning dynamics^S70^. This adjustment reduced the train–test loss gap, indicating improved generalisation (Figure S28), but exploring higher dropout values remains an avenue for future work.

Furthermore, a learning rate sweep was conducted (Figure S29), as the learning rate is highly sensitive to dataset size and noise, with smaller datasets typically requiring lower values for stable convergence. This confirmed learning rate of 1 × 10^−3^ as optimal. Notably, this is the same learning rate used for the Shakespeare corpus, and was therefore retained, further supporting its robustness across datasets. The loss and perplexity curves are shown in Figure S31.

On DBAASP, metric differences narrowed considerably relative to the Shakespeare results (Figure S30): the GPT’s best test loss was only marginally worse than the LSTM’s, and both models converged at comparable epochs. Despite this convergence in headline metrics, the train–test gap remained markedly asymmetric: the LSTM retained a substantial gap compared to near-complete convergence in the GPT, indicating continued overfitting in the recurrent model. This is mechanistically consistent—the LSTM’s lower training loss on Shakespeare was driven by memorising its lexical repetitiveness, a property sparse in biologically diverse AMP sequences. Across both corpora, the GPT maintained a more balanced train–test loss profile, the criterion most directly relevant to generating novel, out-of-distribution peptide sequences.

#### B. Shakespeare Benchmark: LSTM vs. GPT

**FIG. S21.**
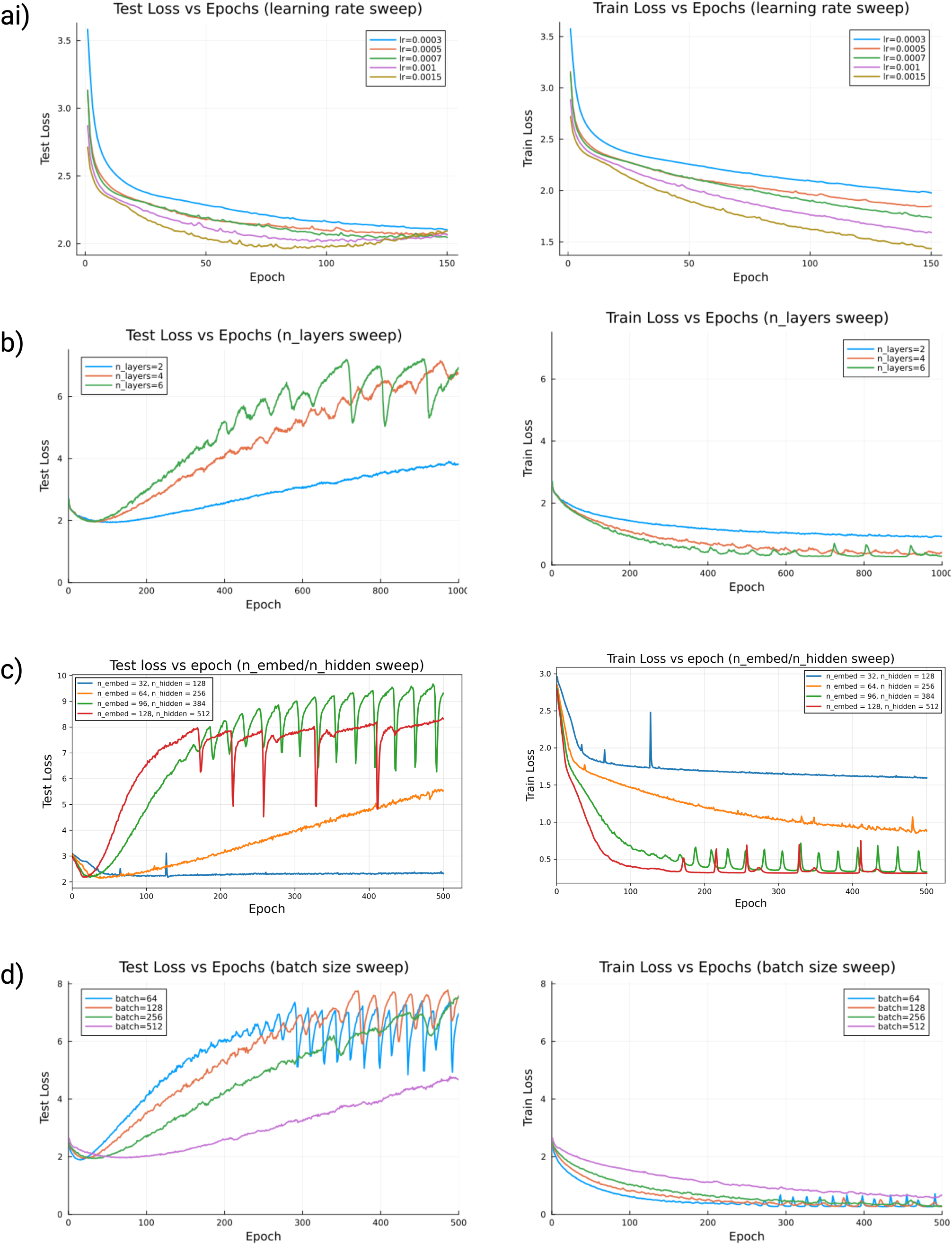
Evolution of training and test cross-entropy loss across epochs for GPT model under varying hyperparameters: (a) learning rate, (b) number of layers, (c) embedding and hidden dimensions, and (d) batch size. Results illustrate the trade-off between model capacity, optimisation dynamics, and generalisation.

**FIG. S22.**
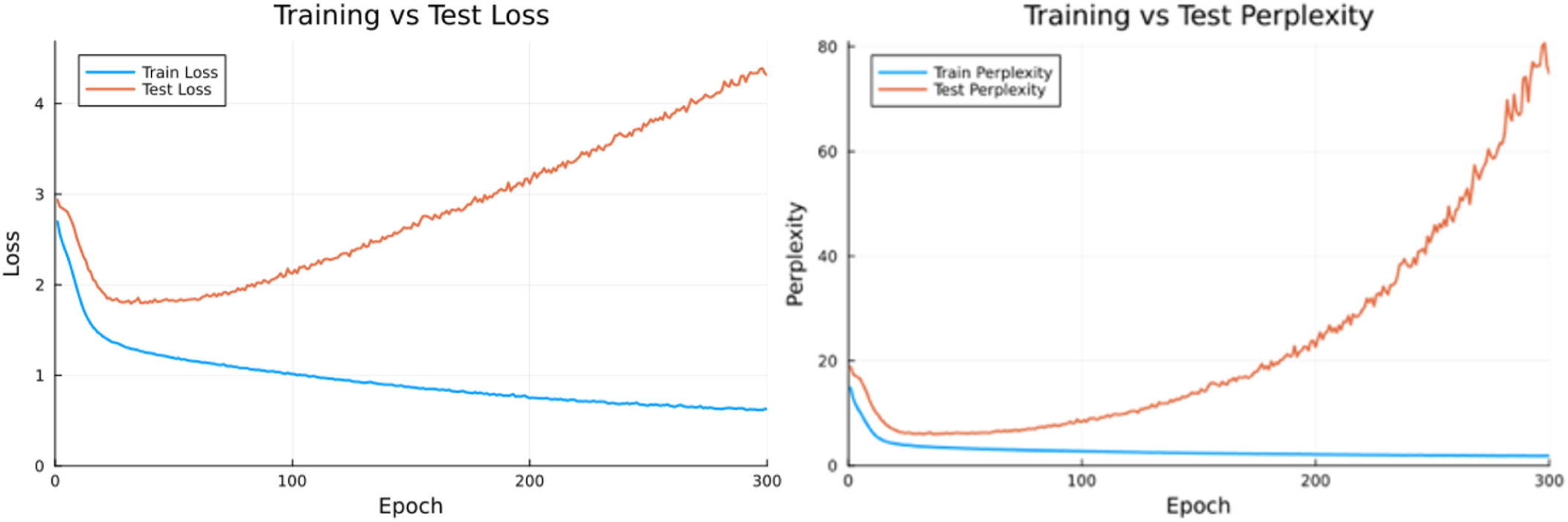
Cross-entropy loss and perplexity curves for a standalone GPT model trained on the Shakespeare corpus. Model hyperparameters were: embedding size 64, hidden size 256, 4 attention heads, query/key and value dimensions of 16, 4 layers, sequence length 32, batch size 256, dropout 0.0, seed 42, and learning rate 1 × 10^−3^. The model was trained for 300 epochs using a 90/10 train-test split.

**FIG. S23.**
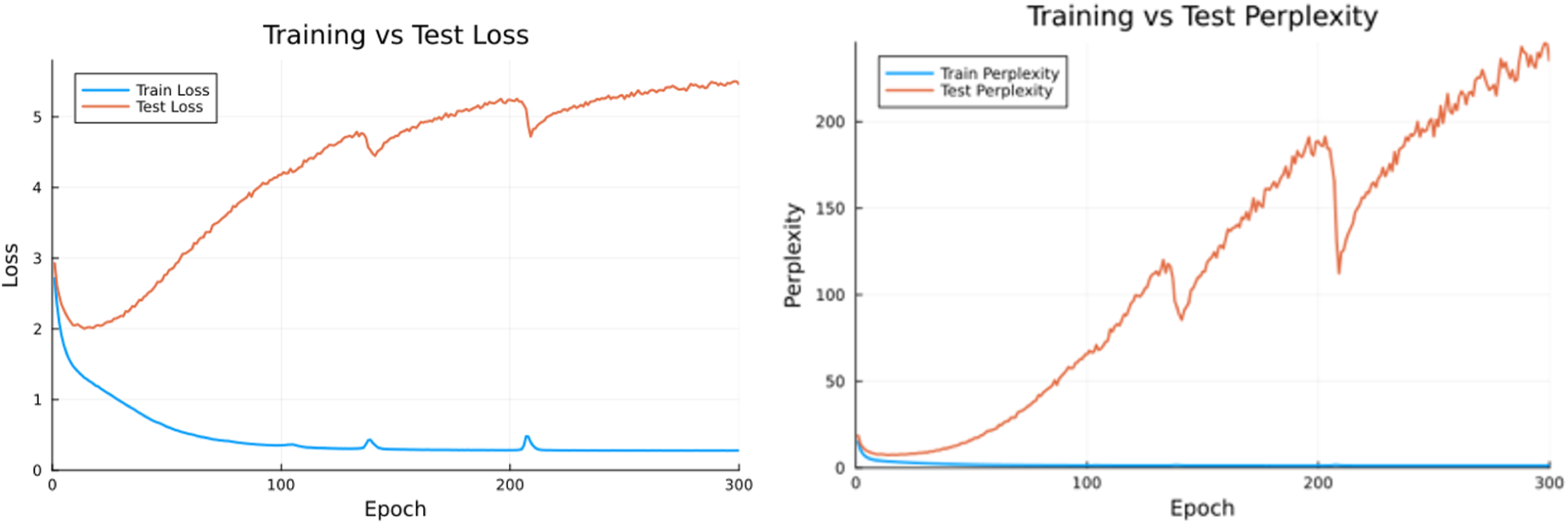
Cross-entropy loss and perplexity curves for a LSTM model trained on the Shakespeare corpus. Model hyperparameters were: embedding size 64, hidden size 256, 2 layers, seed 42, sequence length 32, batch size 256, learning rate 1 × 10^−3^, and 300 training epochs, using a 90/10 train-test split.

**FIG. S24.**
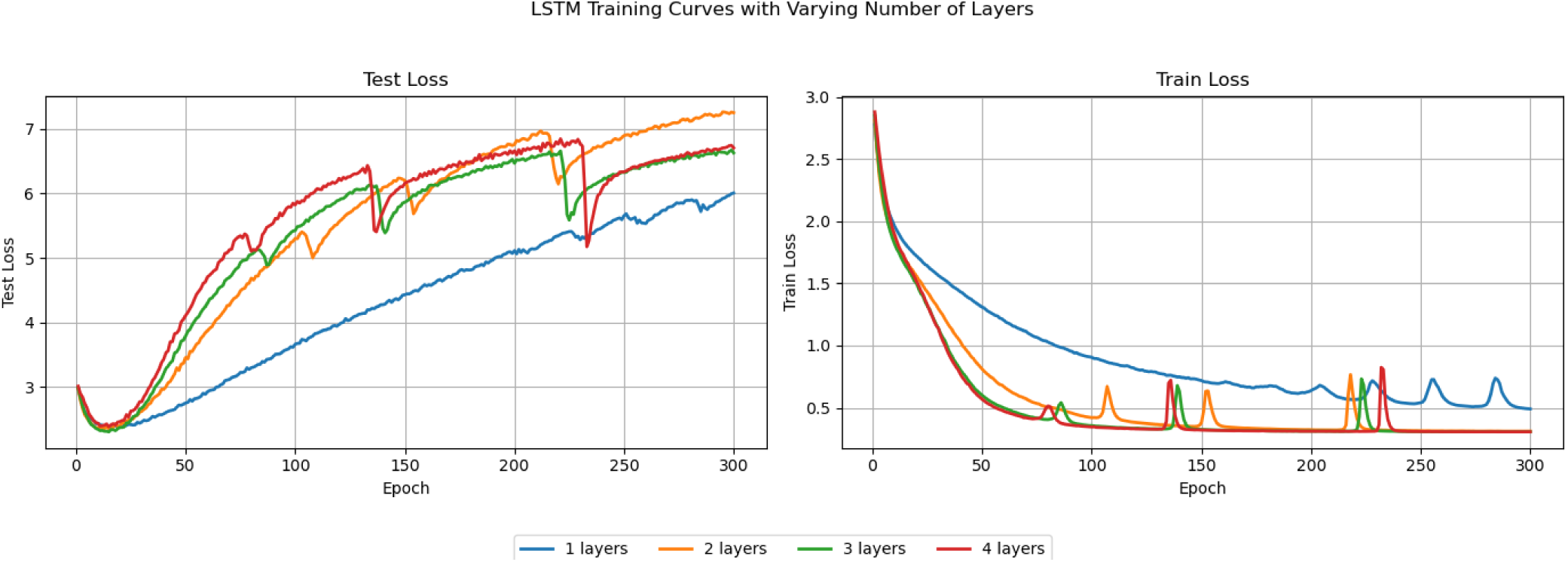
Evolution of training and test cross-entropy loss across epochs for LSTM model under varying number of layers. Model hyperparameters were: embedding size 64, hidden size 256, sequence length 32, batch size 256, learning rate 1 × 10^−3^, seed 1337, and 300 training epochs, using a 90/10 train-test split. Minimum test loss values were 2.3765 (1 layer), 2.3496 (2 layers), 2.3010 (3 layers), and 2.3804 (4 layers). Corresponding minimum training loss values were 0.4908, 0.3109, 0.3061, and 0.3060, respectively.

##### Word-Level Token Recovery — GPT vs LSTM across Epochs

*Input sequence: NERS TRUNCHEONS TRUNDLE TRUNK TR · Character positions 0–31 · Spaces shown as_*

###### Legend

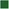 Exact match — all characters correct (100%)

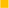 Partial match — ≥50% correct, or correct 3-character prefix

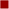 Mostly wrong — <50% correct characters

**FIG. S25.**
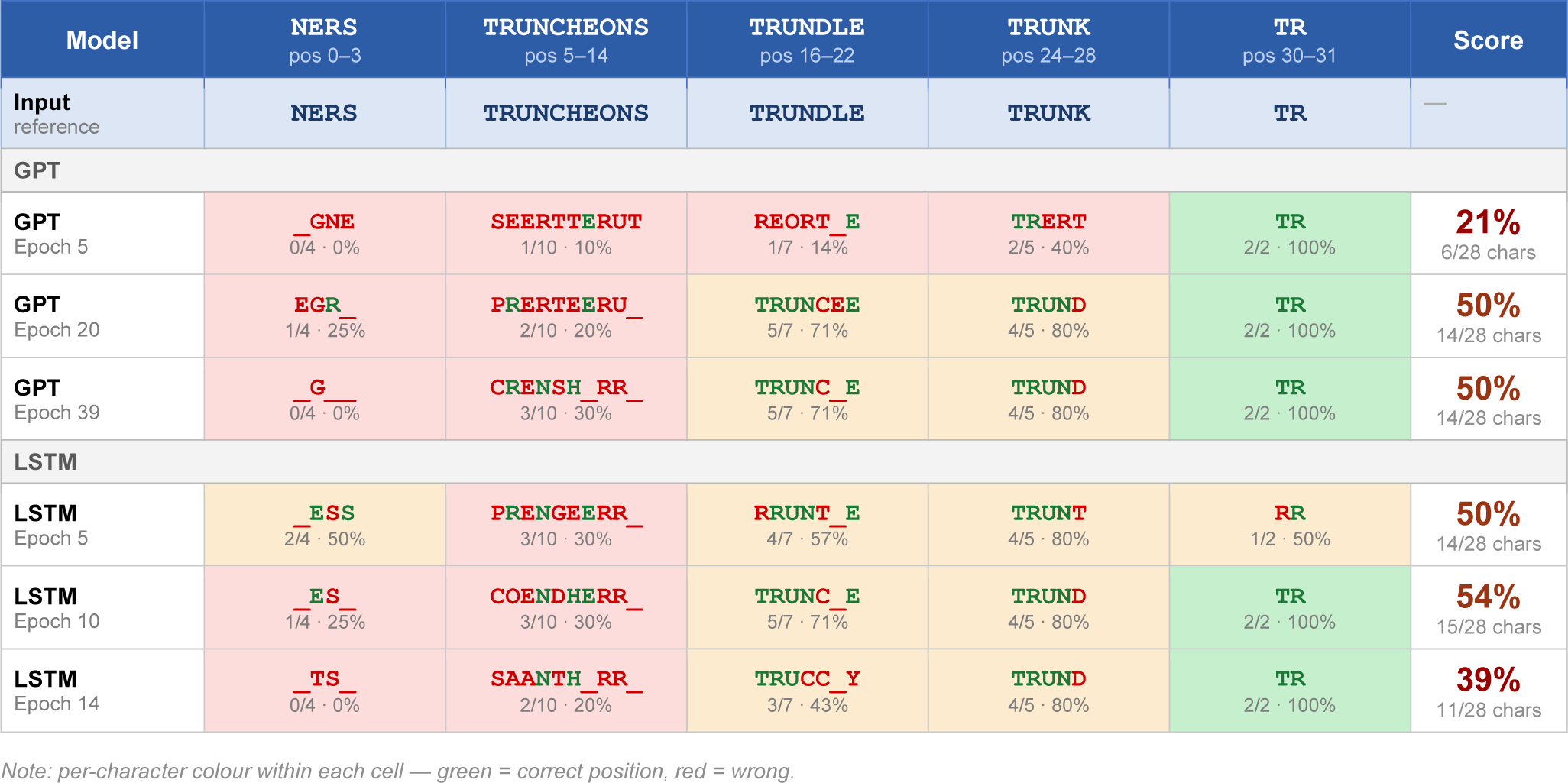
Probe analysis of GPT model output at selected training epochs for one input batch. Each row shows the model-generated sequence conditioned on the probe input. For each model, three samples were taken at evenly spaced intervals between epochs 5 and each model’s best test loss epoch. *Note.* Although at different speeds, both the GPT and LSTM recover key substructures such as the stems “TRUNC–” and “TRUND–”, and both acquire word-boundary placement (indicated by underscore positions) relatively early in training, suggesting that character-level segmentation cues are more readily learned than internal character order. Despite this, the LSTM output stagnates and regresses by epoch 14—reintroducing errors such as “TRUCC” in a previously correct stem region—suggesting susceptibility to overfitting and limited capacity to refine representations beyond a local optimum. The GPT, by contrast, continues to improve monotonically across all checkpoints. This is further reflected in sequence completeness: the LSTM’s outputs are shorter and more fragmented, whereas the GPT produces increasingly full-length, coherent sequences, demonstrating that its architecture scales more effectively with additional training and is better suited to the compositional structure of the target sequences.

#### C. Shakespeare Auxiliary Analysis

**FIG. S26.**
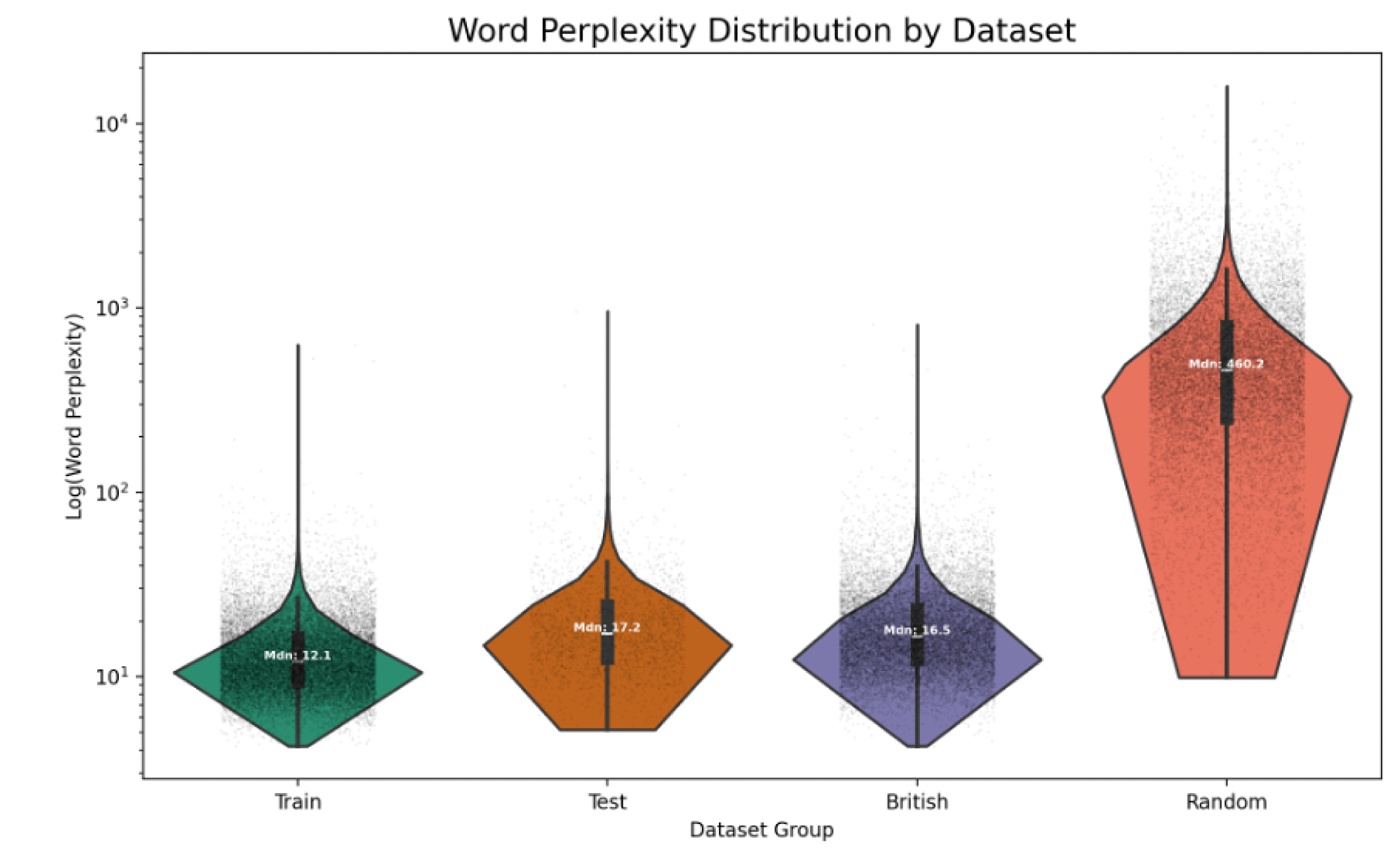
Distribution of GPT word-level perplexity across dataset groups (Train, Test, British, Random) shown on a logarithmic scale. Violin plots illustrate the density and spread of perplexity values, with overlaid scatter points representing individual observations. Median values are annotated for each group. The in-sample dataset ( 24,500 words) consists of words drawn directly from the Shakespeare corpus used for training and evaluation. The near-sample dataset ( 18,100 words) was constructed from the Corpus of Contemporary American English (COCA), with all overlaps with the Shakespeare corpus removed^S71^. This set contains words that are morphologically similar but not identical to those seen during training, as well as modern or technical vocabulary. The out-of-sample dataset ( 18,000 words) was created using a random generator to combine alphabet characters into sequences.

**FIG. S27.**
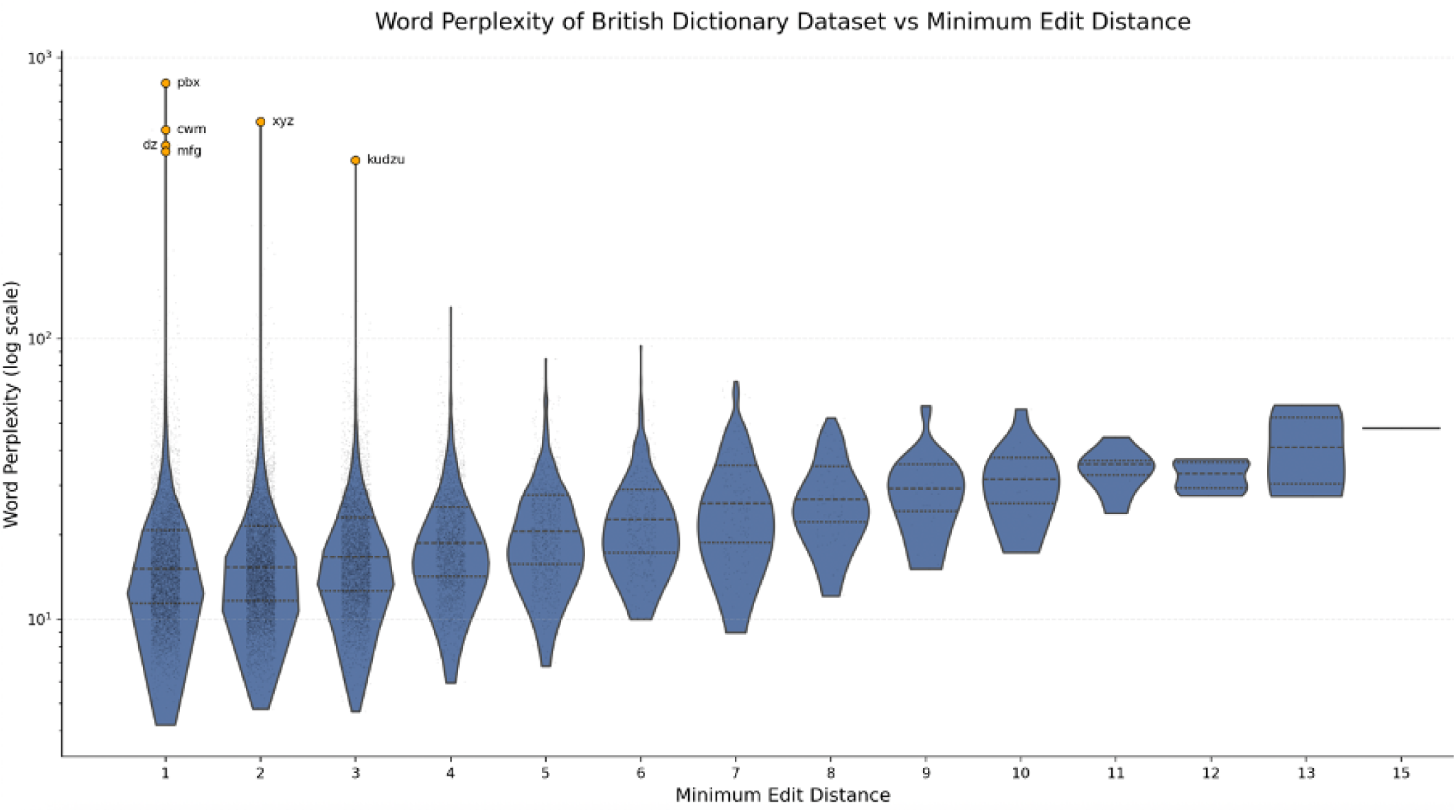
GPT word-level perplexities of 100 highest perplexity words from the test (orange) and ‘British’(purple) datasets. The mean for the in-sample (black dashed line) and the British (green dashed line) dataset are also marked. *Note.* Perplexity gradually increases with distance from the training distribution, indicating that the model has learned transferable orthographic and morphological patterns rather than simply memorising words.

**FIG. S28.**
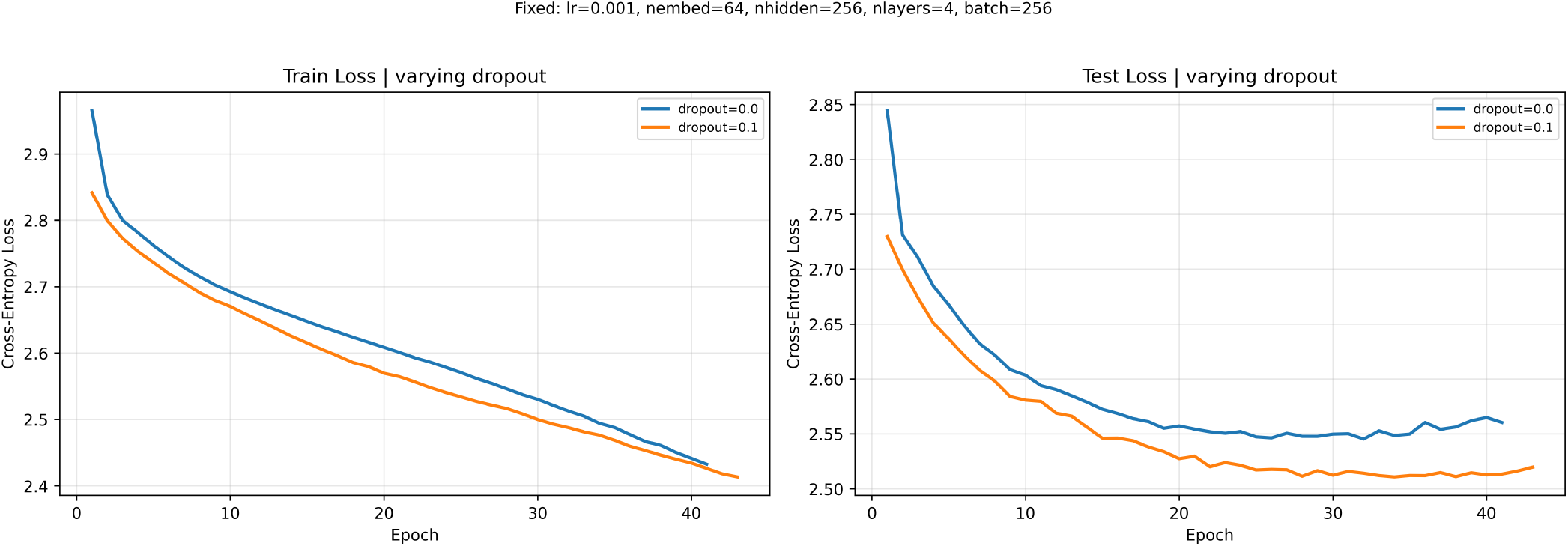
Effect of dropout (dp=0.1) addition to the GPT train and test cross-entropy loss, training on DBAASP dataset. Adding dropout helps to lower the train – test gap, improving generalisation.

**FIG. S29.**
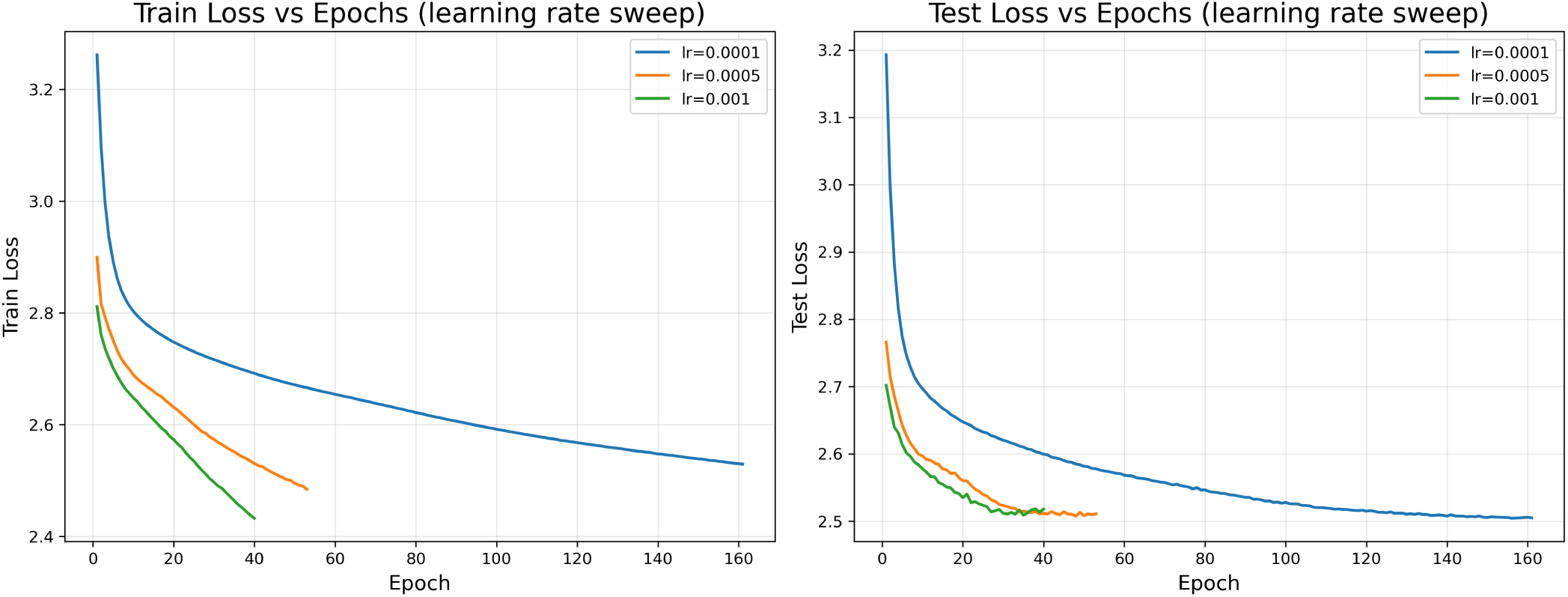
GPT learning rate (lr) sweep, training on DBAASP dataset with dropout 0.1, seed 42. Minimum train loss values were 2.529 (lr = 1 × 10^−4^), 2.485 (lr = 5 × 10^−4^), and 2.433 (lr = 1 × 10^−3^). Corresponding minimum test loss values were 2.504, 2.508, 2.509, respectively.

#### D. DBAASP Training and Hyperparameter Adaption

**FIG. S30.**
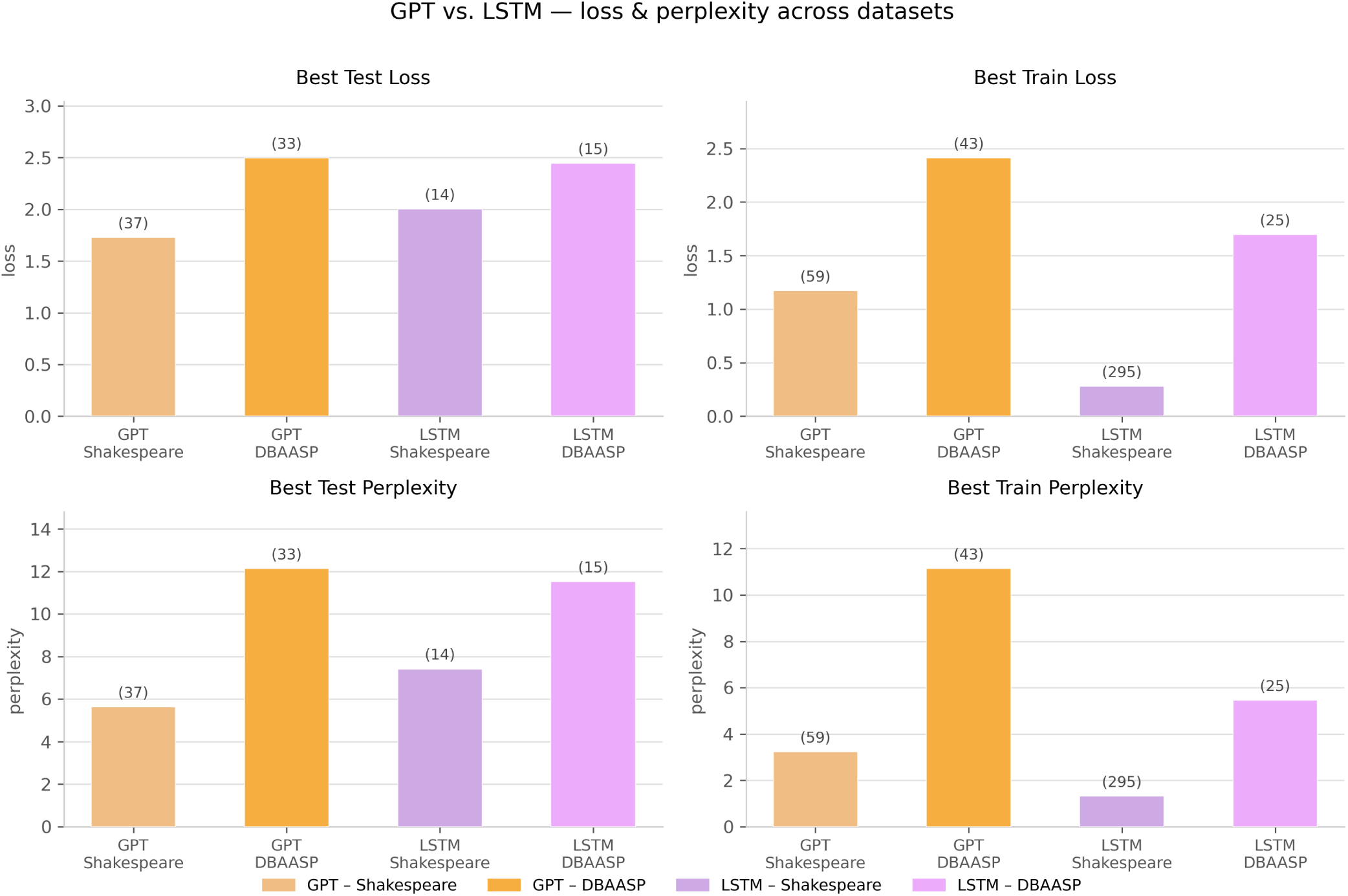
Comparison of GPT and LSTM performance on the Shakespeare and DBAASP datasets, showing best test loss, best training loss, best test perplexity, and best training perplexity. On Shakespeare, the LSTM achieved a lower best training loss (0.279 vs. 1.173) but a substantially worse best test loss (2.003 vs. 1.728) and perplexity (7.41 vs. 5.63), with its optimal epoch reached far earlier (14 vs. 37), indicating memorisation rather than generalisation. On DBAASP, headline metrics converged between architectures, but the train–test gap remained markedly larger for the LSTM (Δ = 0.75) than the GPT (Δ = 0.09). The results highlight dataset-dependent differences, with both models achieving substantially lower loss and perplexity on Shakespeare than on DBAASP.

**FIG. S31.**
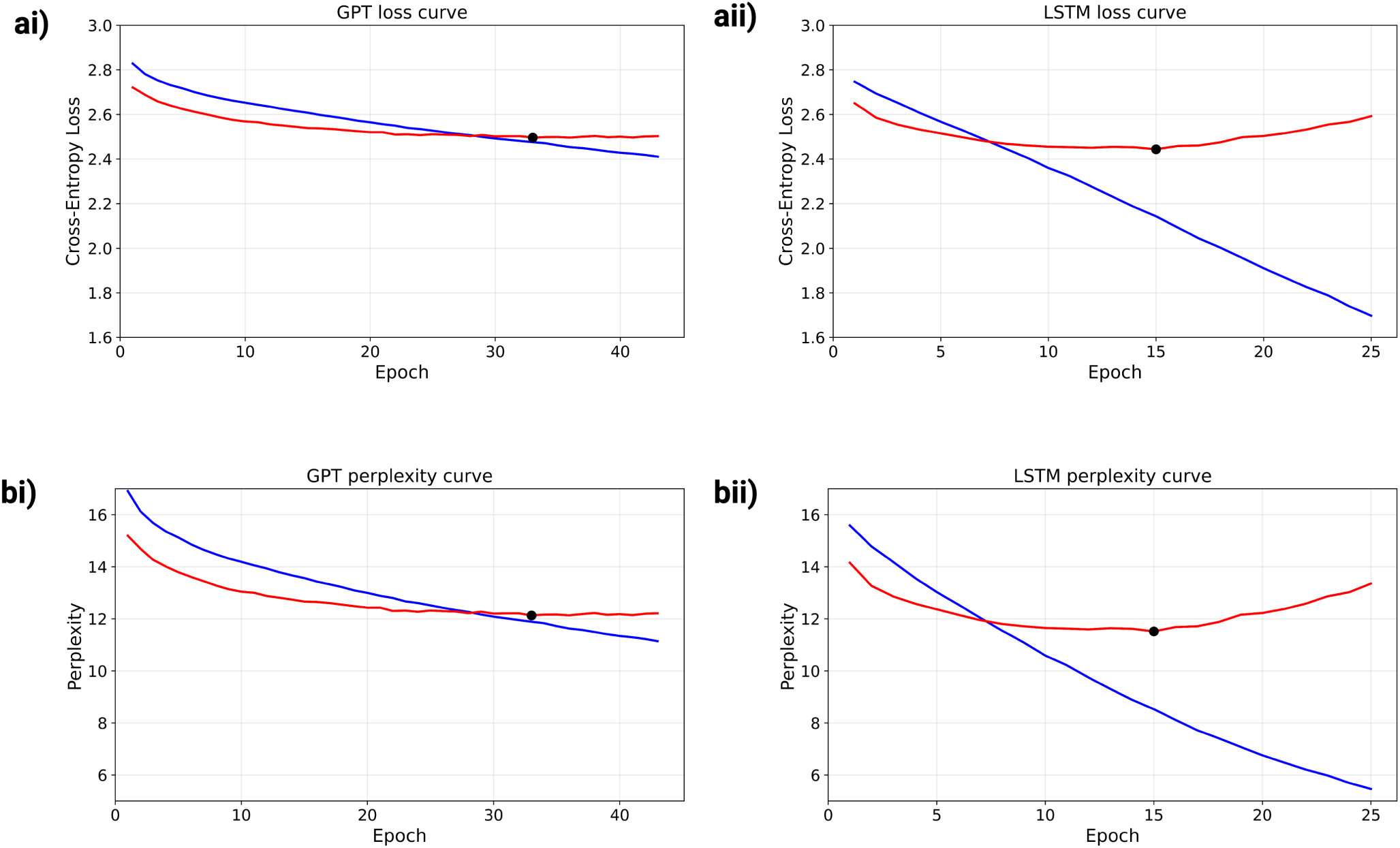
DBAASP LSTM and GPT training curves. Panels ai) and aii) show the training and test loss curves for the GPT and LSTM models, respectively, while panels bi) and bii) show the corresponding training and test perplexity curves. Blue denotes training loss, whereas red denotes test loss. Black markers indicate the epoch at which the minimum test loss or minimum test perplexity was achieved. Both models were trained using an embedding dimension of 64, hidden dimension of 256, sequence length of 32, batch size of 256, learning rate of 1 × 10^−3^, seed 42, dropout of 0.1, and a 90:10 train-test split. Training employed early stopping with a patience of 5 epochs and a minimum improvement threshold of 0.001. The GPT achieved best test loss 2.496 (perplexity 12.13) at epoch 33; the LSTM achieved 2.444 (perplexity 11.52) at epoch 15. Despite similar headline values, the LSTM train–test gap (Δ = 0.75) was substantially larger than the GPT’s (Δ = 0.09), indicating continued overfitting in the recurrent model.

**FIG. S32.**
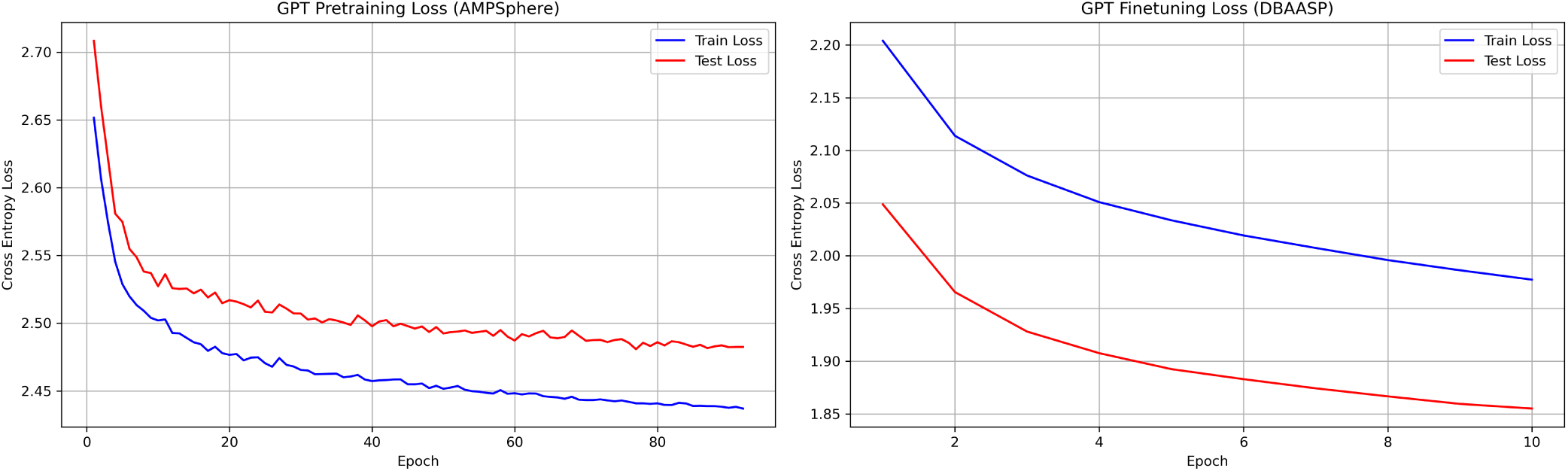
Training and test loss curves for best configuration GPT pre-training (AMPSphere, left) and fine-tuning (DBAASP, right). Fine-tuning achieved a best test loss of 1.79, a 28% reduction relative to the standalone GPT trained on DBAASP alone (2.49), confirming that AMPSphere pre-training provides an effective initialisation for activity-guided fine-tuning. The pre-trained model used embedding dimension 128, hidden dimension 512, 4 attention heads, query/key and value dimensions of 32, 4 layers, sequence length 128, batch size 256, dropout 0.1, and learning rate 1 × 10^−3^, with early stopping (patience = 15, minimum improvement = 0.001) and a 90:10 train-test split. The model was subsequently fine-tuned on DBAASP using the same architecture with a learning rate of 1 × 10^−4^, dropout 0.1 and batch size 32.

#### E. Transfer Learning

**FIG. S33.**
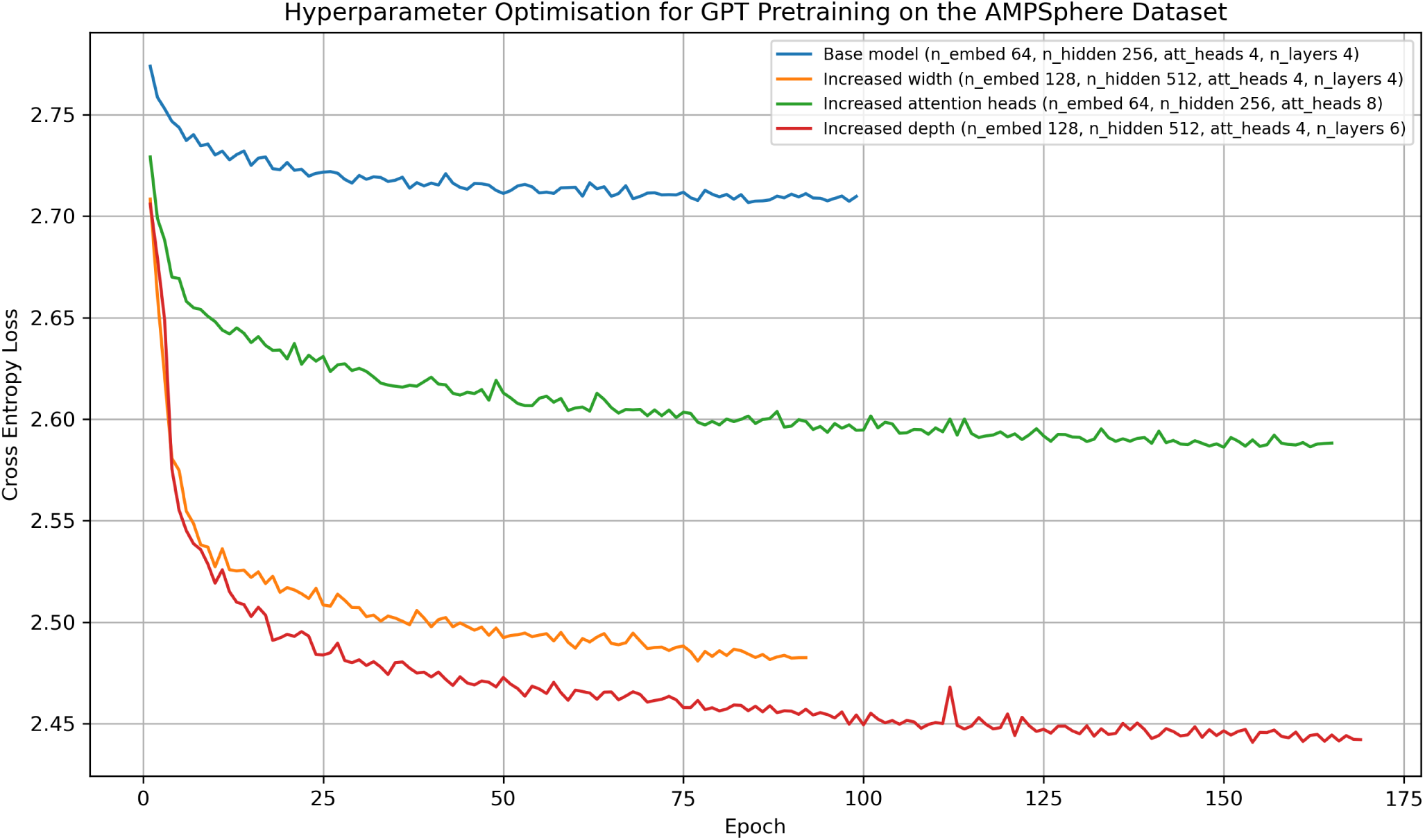
Hyperparameter optimisation for GPT pre-training on the AMPSphere dataset. Cross-entropy training loss is shown across epochs for the base model and selected architectural variants. The base model used embedding dimension = 64, hidden dimension 256, 4 attention heads, query/key and value dimensions of 16, 4 layers, sequence length = 128, batch size = 256, dropout = 0.1, learning rate 1 × 10^−3^, early stopping (patience = 15, minimum improvement = 0.001), and a 90:10 train-test split. Relative to the base model, increasing embedding dimension and model depth reduced the loss most substantially, with the deeper and wider configuration achieving the lowest final pre-training loss.

**FIG. S34.**
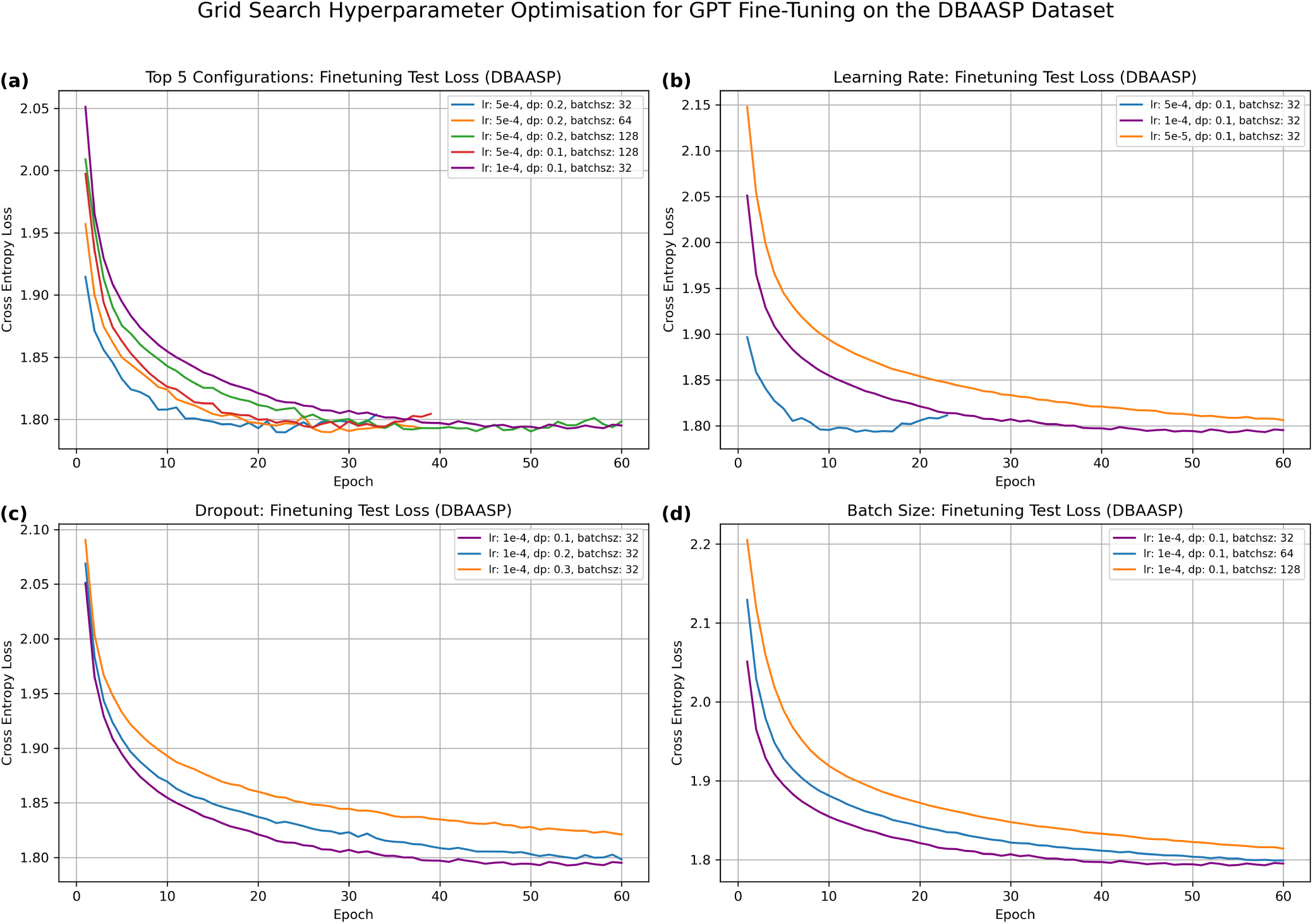
Hyperparameter grid search results for GPT fine-tuning on the DBAASP antimicrobial peptide dataset. Test cross-entropy loss is shown across training epochs for selected configurations. (a) The five best-performing configurations were identified during the grid search. (b–d) Sensitivity analyses showing the effect of learning rate, dropout, and batch size, respectively, while keeping the other parameters fixed. The grid search evaluated combinations of learning rate (5 × 10^−5^, 1 × 10^−4^, 5 × 10^−4^), batch size (32, 64, 128), and dropout (0.1, 0.2, 0.3) using the Adam optimiser. The pre-trained model used embedding dimension 128, hidden dimension 512, 4 attention heads, query/key and value dimensions of 32, 4 layers, sequence length 128, batch size 256, dropout 0.1, and learning rate 1 × 10^−3^, with early stopping (patience = 15, minimum improvement = 0.001) and a 90:10 train-test split.

**FIG. S35.**
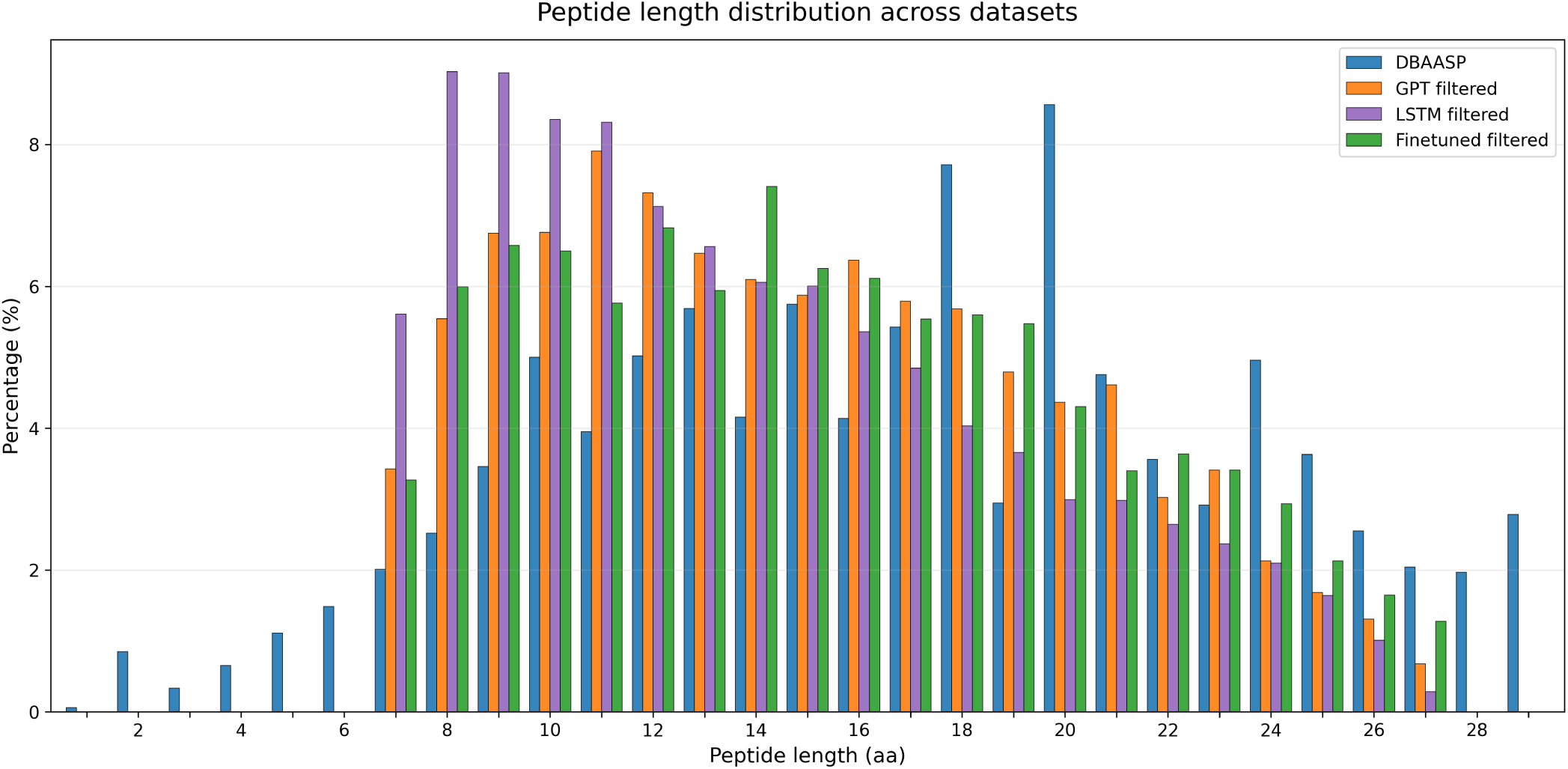
Peptide length distribution across the reference DBAASP dataset and generated peptide sets (GPT, fine-tuned GPT, LSTM) after physicochemical filtering. The DBAASP reference exhibits the highest central tendency (mean = 16.94, median = 17). The fine-tuned GPT most closely approximates this distribution (mean = 15.29, median = 15), followed by the standalone GPT (mean = 14.97, median = 14). The LSTM generates the shortest sequences (mean = 13.84, median = 13), with a distribution most shifted away from the reference.

### S5. SEQUENCE GENERATION AND PROPERTY FILTERING

**Table S22.** Retention percentage after physicochemical filtering for sequences generated by each model.

| Model | Training epoch | Retention percentage (%) |
| --- | --- | --- |
| LSTM | — | 28.8 |
| GPT | — | 38.0 |
| Fine-tuned GPT | 10 | 42.5 |
| Fine-tuned GPT | 7 | 42.6 |
| Fine-tuned GPT | 5 | 42.4 |
| Fine-tuned GPT | 3 | 43.7 |

**FIG. S36.**
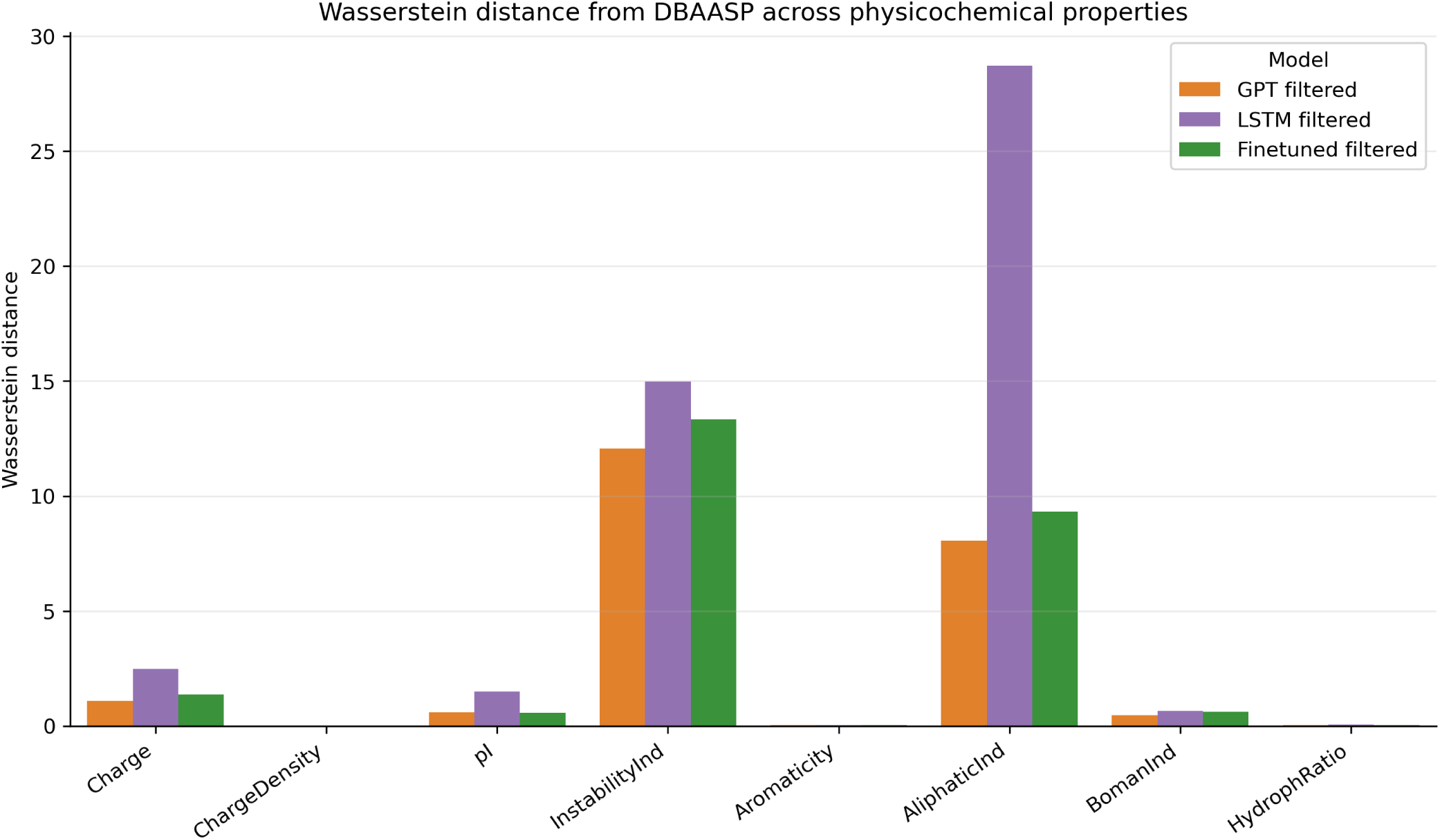
Wasserstein distance from DBAASP across physicochemical properties for generated peptide sets. The Wasserstein distance quantifies the minimum “cost” of transforming one probability distribution into another, where cost is defined as the amount of distribution “mass” moved multiplied by the distance it is transported^S39^. Lower values indicate closer agreement with the reference distribution, while higher values reflect greater divergence.

**FIG. S37.**
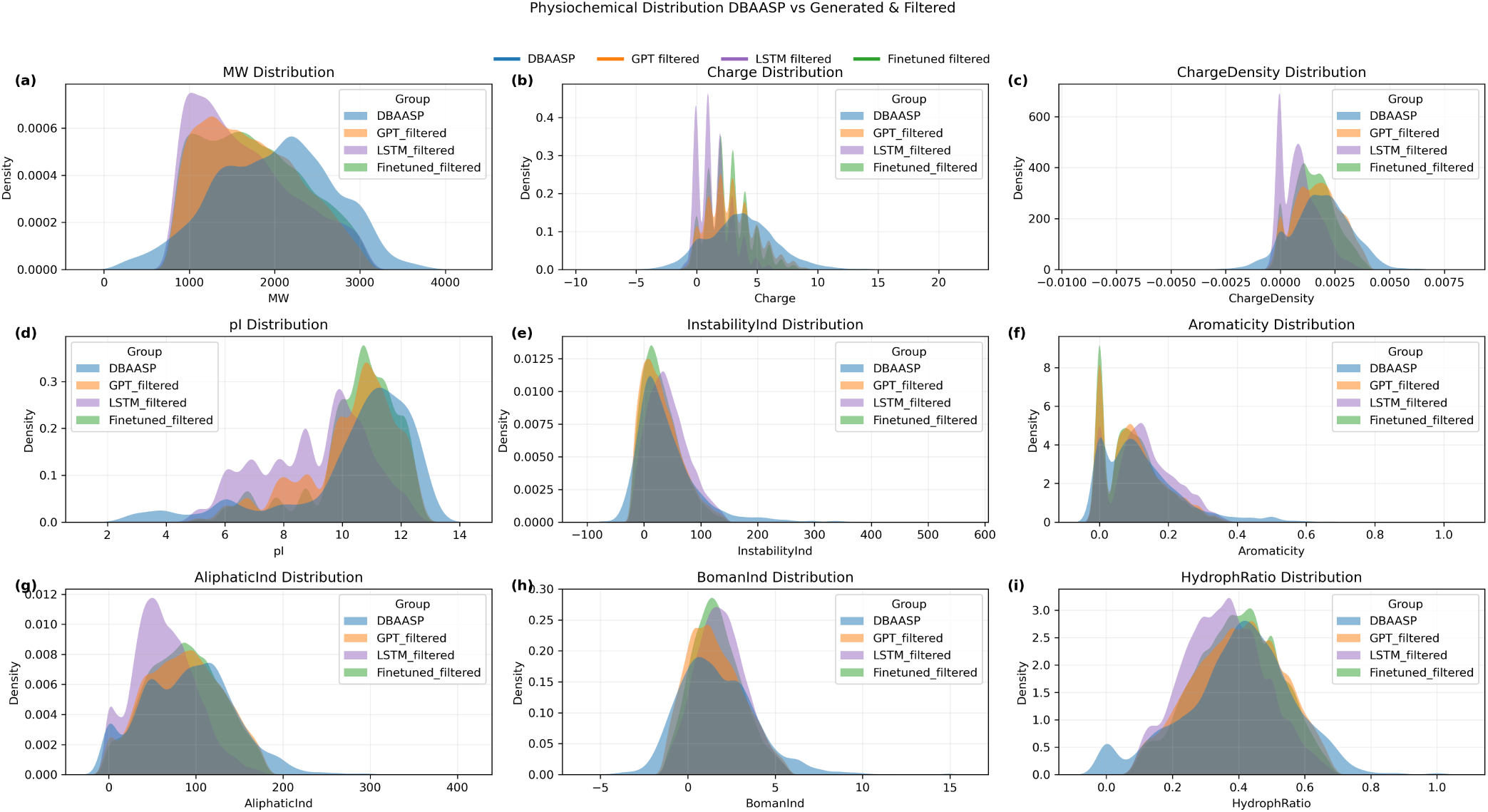
Distributions of key physicochemical properties for DBAASP peptides and generated peptides before and after filtering for the GPT, fine-tuned GPT, and LSTM models. Shown are molecular weight, charge, charge density, isoelectric point (pI), instability index, aromaticity, aliphatic index, Boman index, and hydrophobic ratio. Filtering narrows the generated distributions and improves their agreement with the DBAASP reference set, with the fine-tuned GPT generally showing the closest overall alignment.

**FIG. S38.**
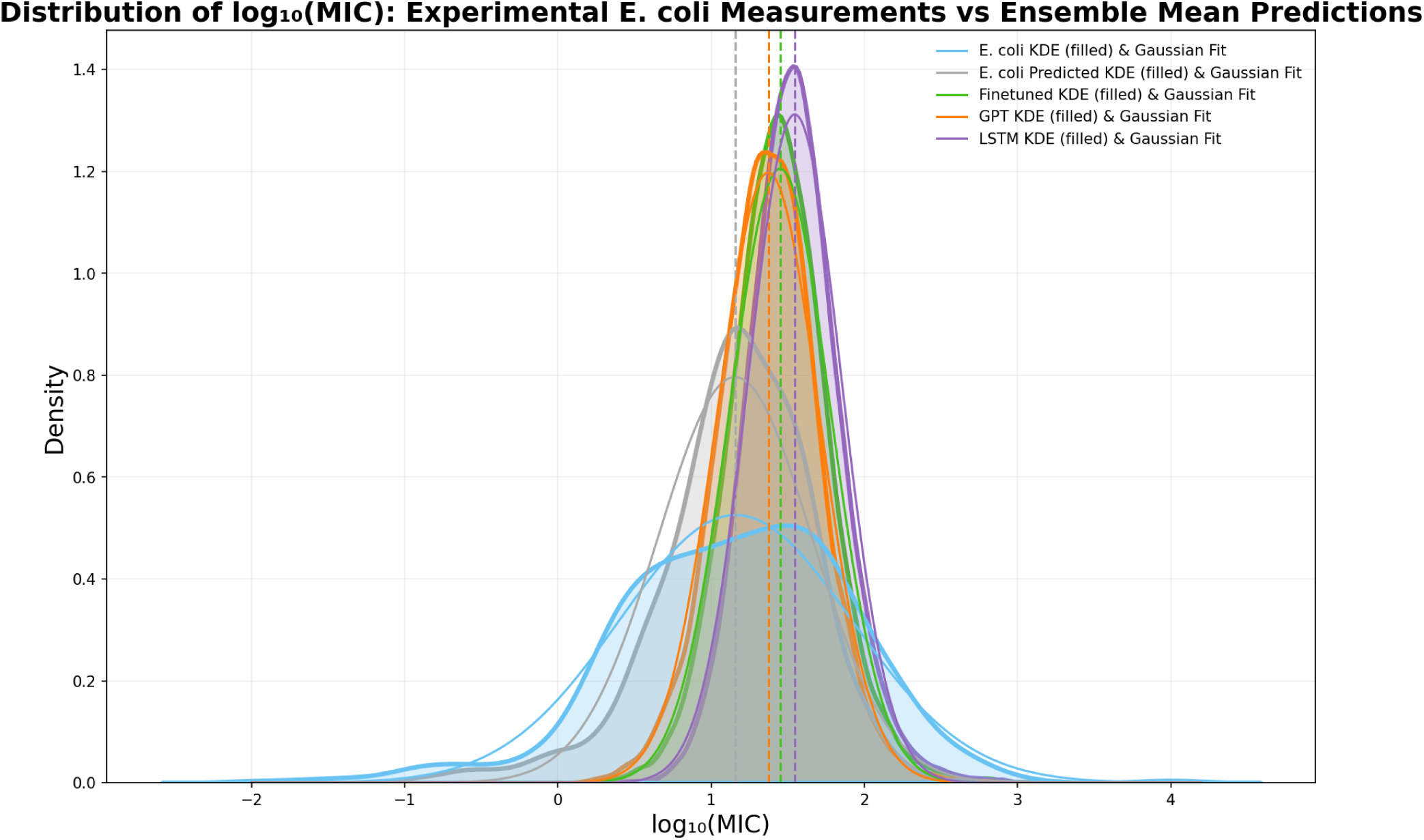
Distribution of log_10_(MIC) [µg mL^−1^] for experimentally validated *E. coli* peptides and ensemble-predicted activities for generated sequences. Predictions are obtained by averaging outputs from three models trained with different random initialisations (seeds 42, 77, 123). Summary statistics (mean *µ* and standard deviation *σ*) are reported for each group: *E. coli* (*µ* = 1.16, *σ* = 0.76), *E. coli* predicted (*µ* = 1.15, *σ* = 0.543), Fine-tuned (*µ* = 1.45, *σ* = 0.33), GPT (*µ* = 1.37, *σ* = 0.33), and LSTM (*µ* = 1.55, *σ* = 0.30). Lower log_10_(MIC) values indicate higher antimicrobial activity.

**Table S23.**
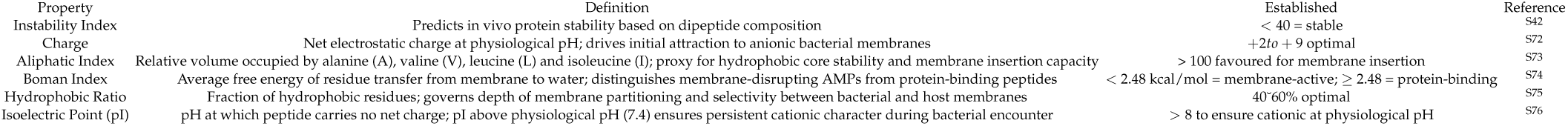
Physicochemical properties of AMPs and their literature-derived established thresholds.

**FIG. S39.**
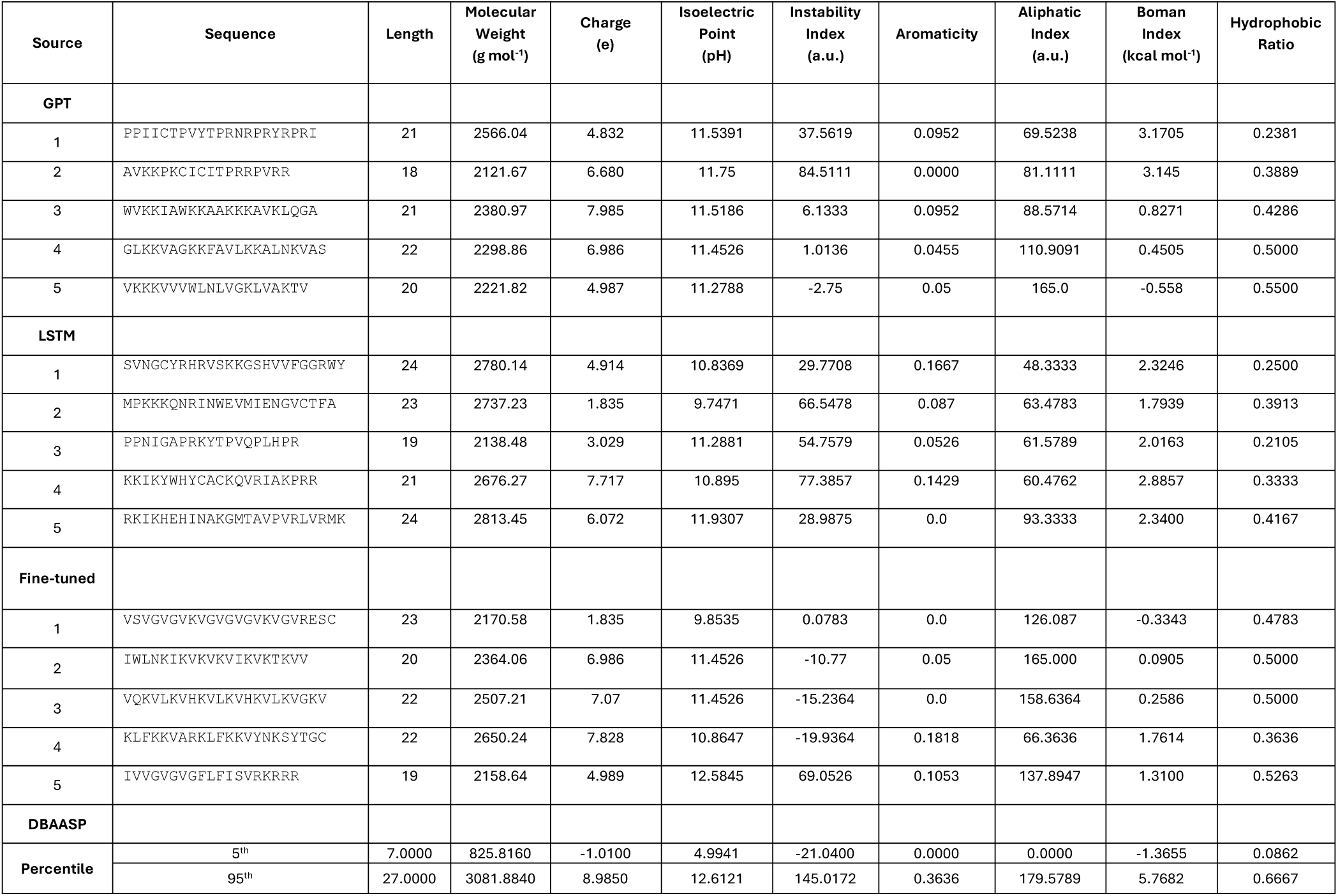
Physicochemical properties of the top five candidate peptides (P1–P5) generated by each model benchmarked against the DBAASP feasibility range (5th–95th percentile)

**Table S24.**
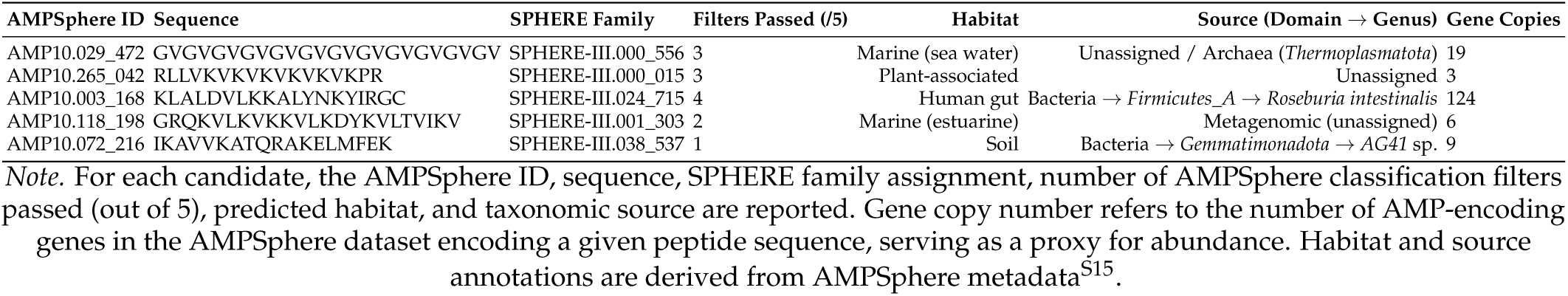
AMPSphere matches identified among the top 100 predicted candidates.

### S6. TOP CANDIDATES

**FIG. S40.**
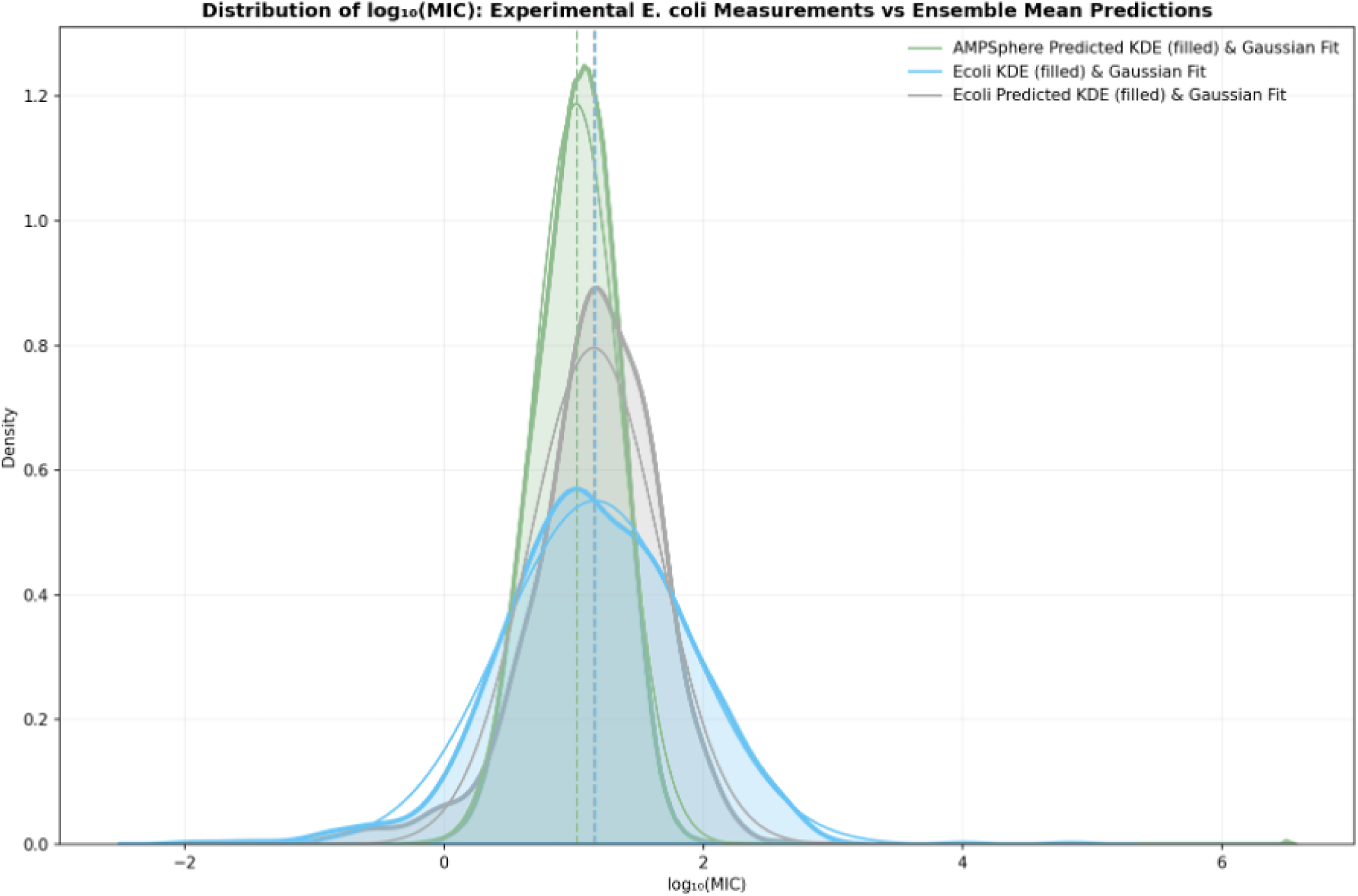
KDE and Gaussian fit of log_10_ MIC distributions for experimental *E. coli* measurements (DRAMP) and ensemble model predictions for *E. coli* and AMPSphere peptides. Filled areas show kernel density estimates; solid lines show fitted Gaussians. Vertical dashed lines indicate the fitted mean (*µ*): experimental *E. coli* (*µ* = 1.17, *σ* = 0.724), predicted *E. coli* (*µ* = 1.16, *σ* = 0.501), and predicted AMPSphere (*µ* = 1.02, *σ* = 0.336). AMPSphere predicted log_10_(MIC) values were clipped to [−3.5, 6.5] prior to fitting to exclude values outside the feasible experimental range.

**FIG. S41.**
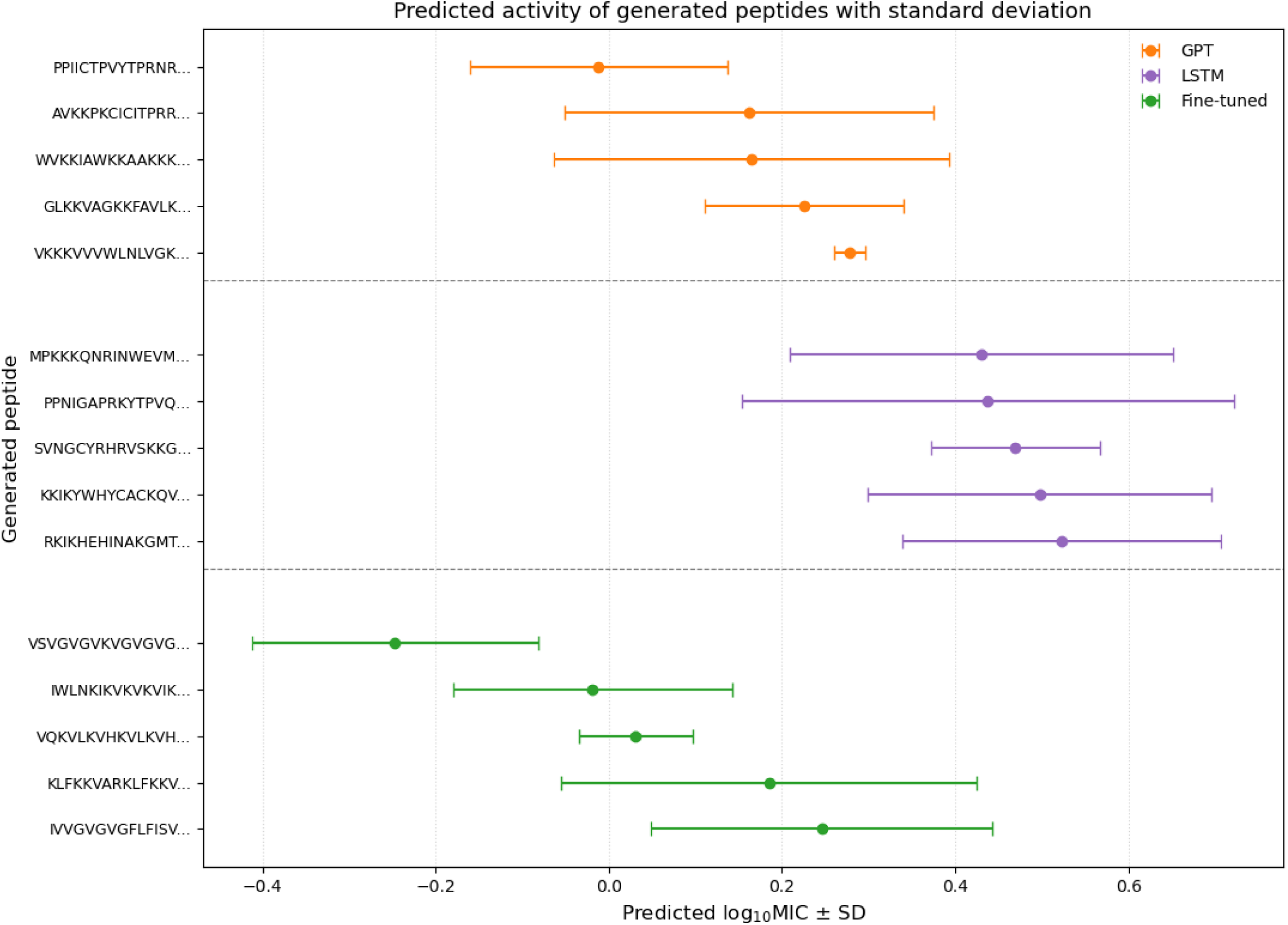
Predicted activity of the top 5 candidates generated by GPT, LSTM, and fine-tuned models. Full details in Figure S42

**FIG. S42.**
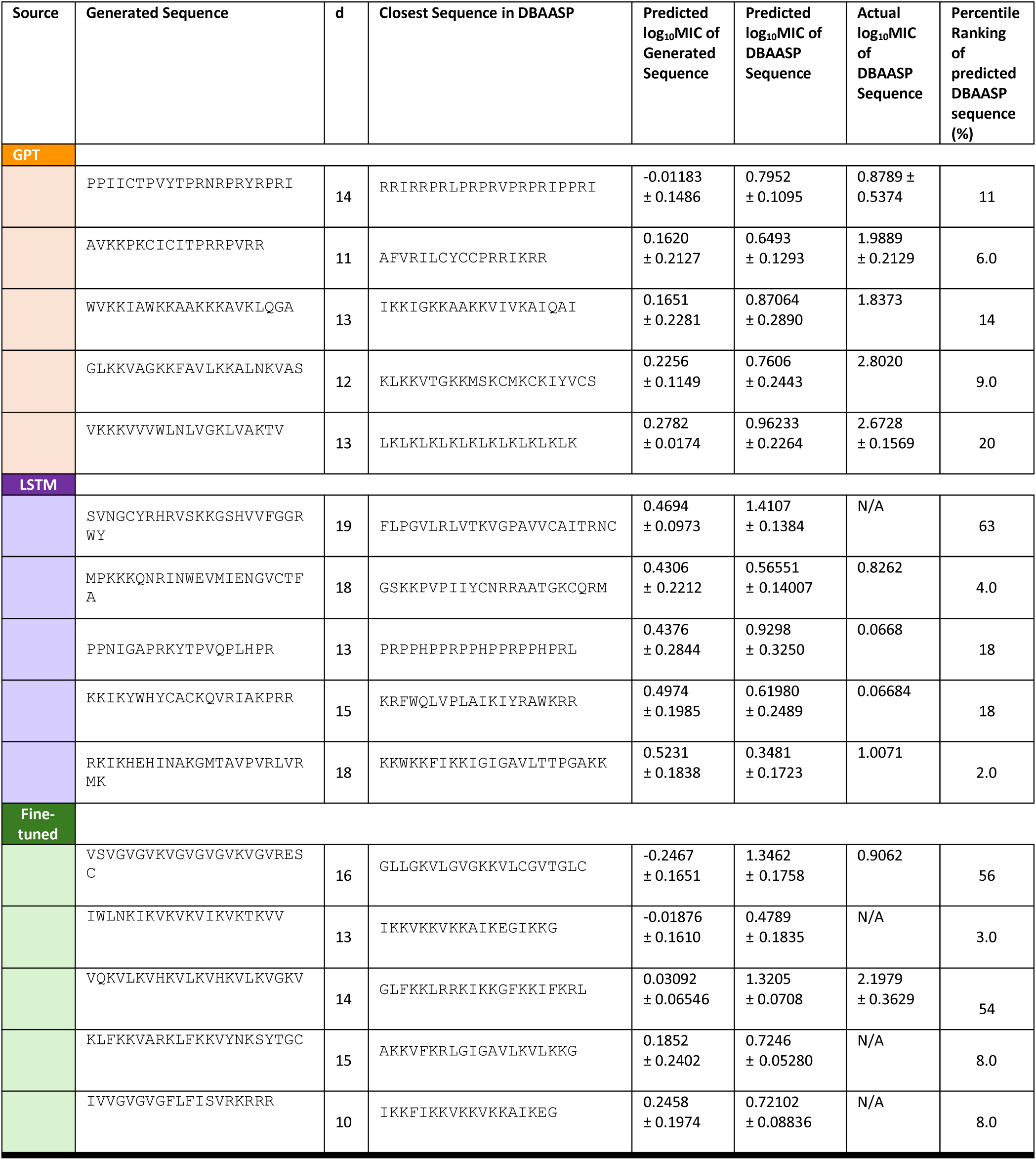
Predicted log_10_(MIC) [µg mL^−1^] against *E. coli* for the top 5 candidates generated by GPT, LSTM, and fine-tuned models, alongside their nearest DBAASP matches identified by weighted Levenshtein distance. *Note.* For each generated sequence, the nearest peptide in DBAASP was identified by weighted Levenshtein distance (*d*). Predicted log_10_(MIC) values (mean ± standard deviation across ensemble members) are reported for both the generated sequence and its closest DBAASP match. Experimental log_10_(MIC) values are included for DBAASP entries where available; N/A indicates no experimentally measured value is recorded in the database^S13^. The percentile rank of the predicted log_10_(MIC) for each DBAASP sequence is reported relative to all ensemble predictions. Across all three generative models, generated peptides tend to exhibit lower predicted log_10_(MIC) values than their closest database counterparts, suggesting improved predicted antimicrobial potency relative to known peptides of similar sequence. Experimental MIC values retrieved from DBAASP reflect assays conducted under varying conditions across different *E. coli* strains, ionic strengths, and growth media; direct quantitative comparison with model predictions — which were trained on aggregated *E. coli* MIC data — should therefore be interpreted with caution. Where multiple experimental measurements were available for a given peptide, the geometric mean log_10_(MIC) is reported.

**FIG. S43.**
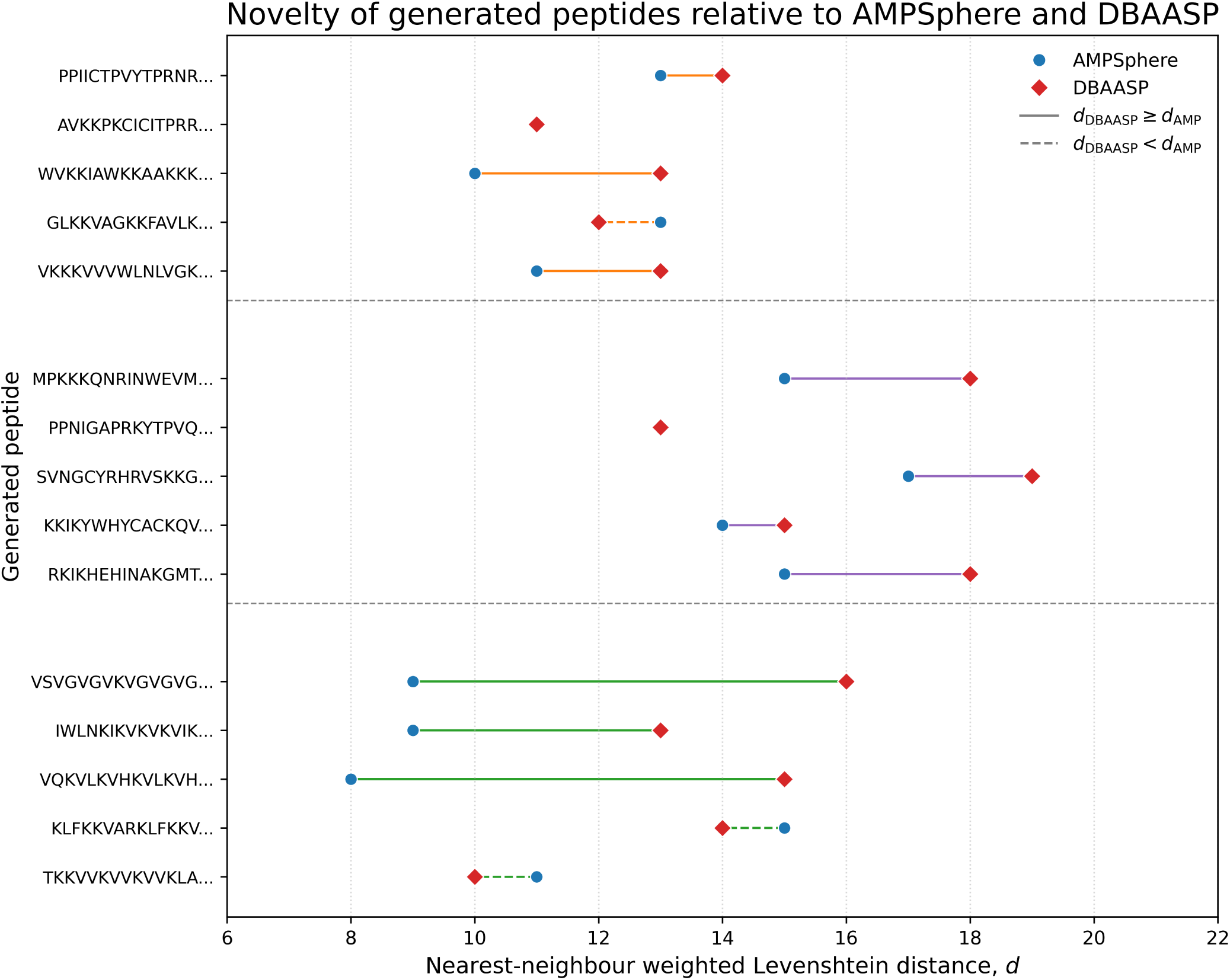
Weighted Levenshtein distances from AMPSphere and DBAASP of the top 5 candidates. Full details in S43.

**FIG. S45.**
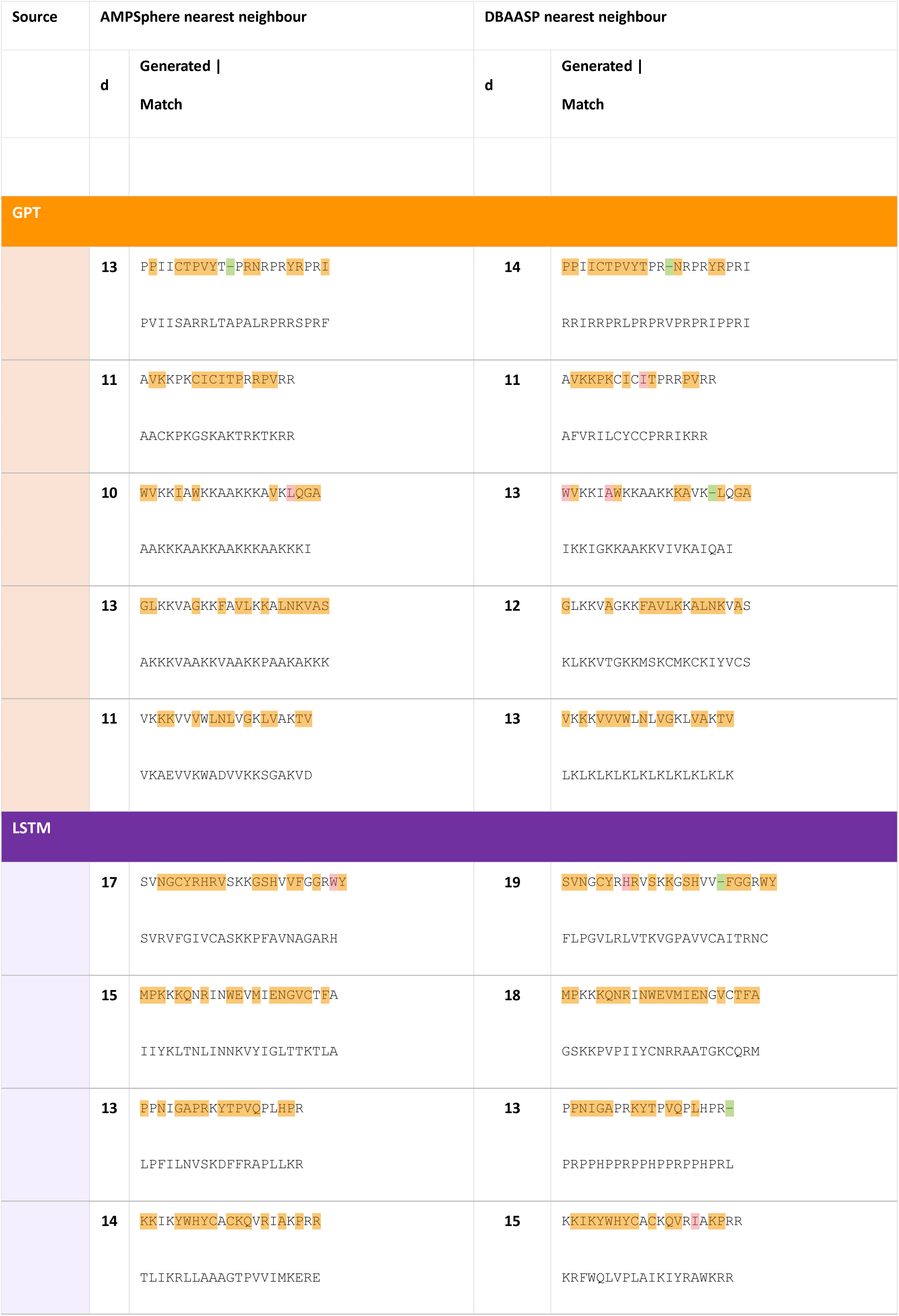
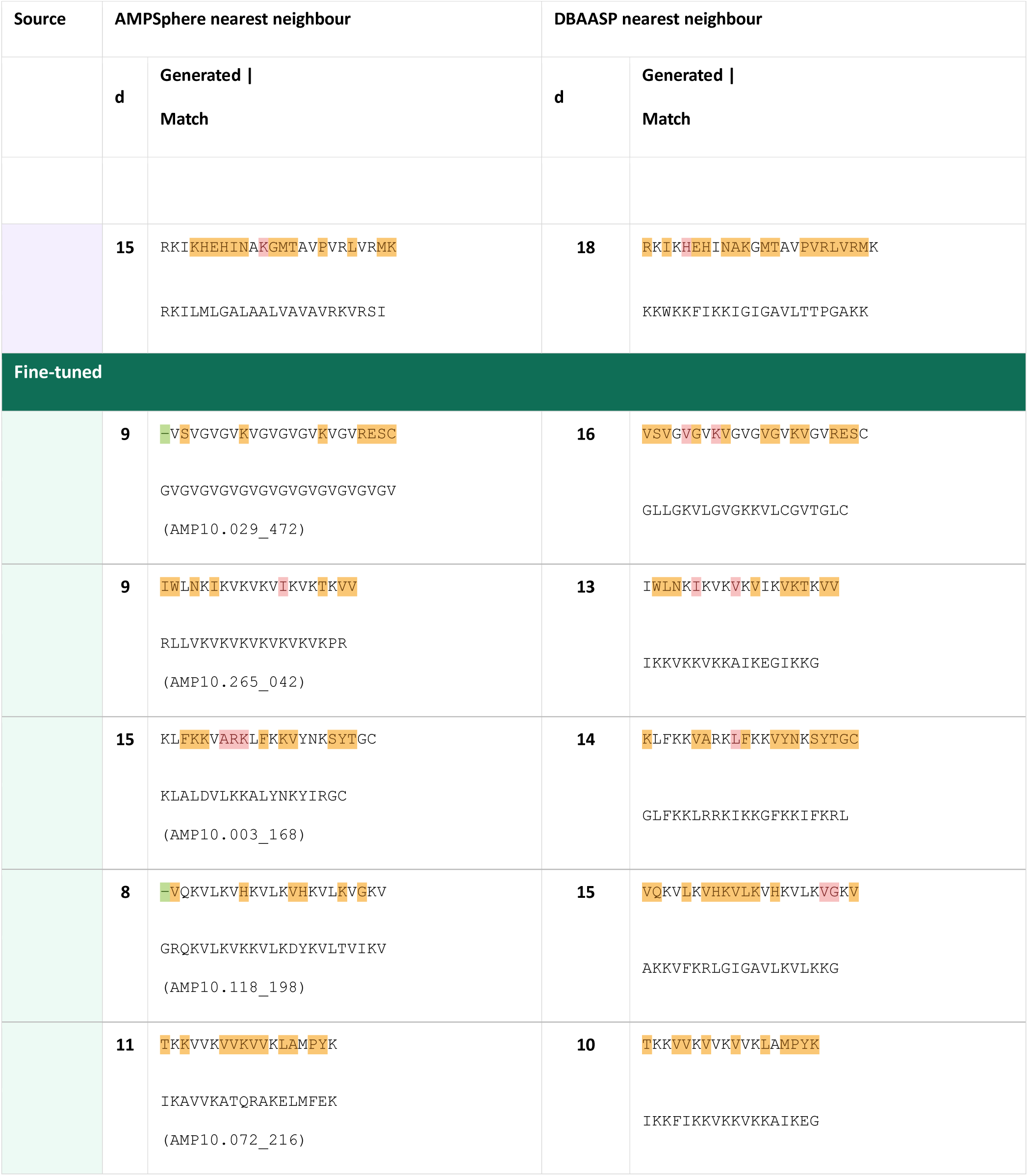
Weighted Levenshtein distances from AMPSphere and DBAASP of the top 5 candidates. Weighted Levenshtein (d): substitution (orange) = 1, insertion (green)/deletion(red) = 2. For each generated sequence the highlighted version (top) is aligned against its nearest AMPSphere match (left column) and nearest DBAASP match (right column); the plain match sequence appears directly below.

**Table S25.**
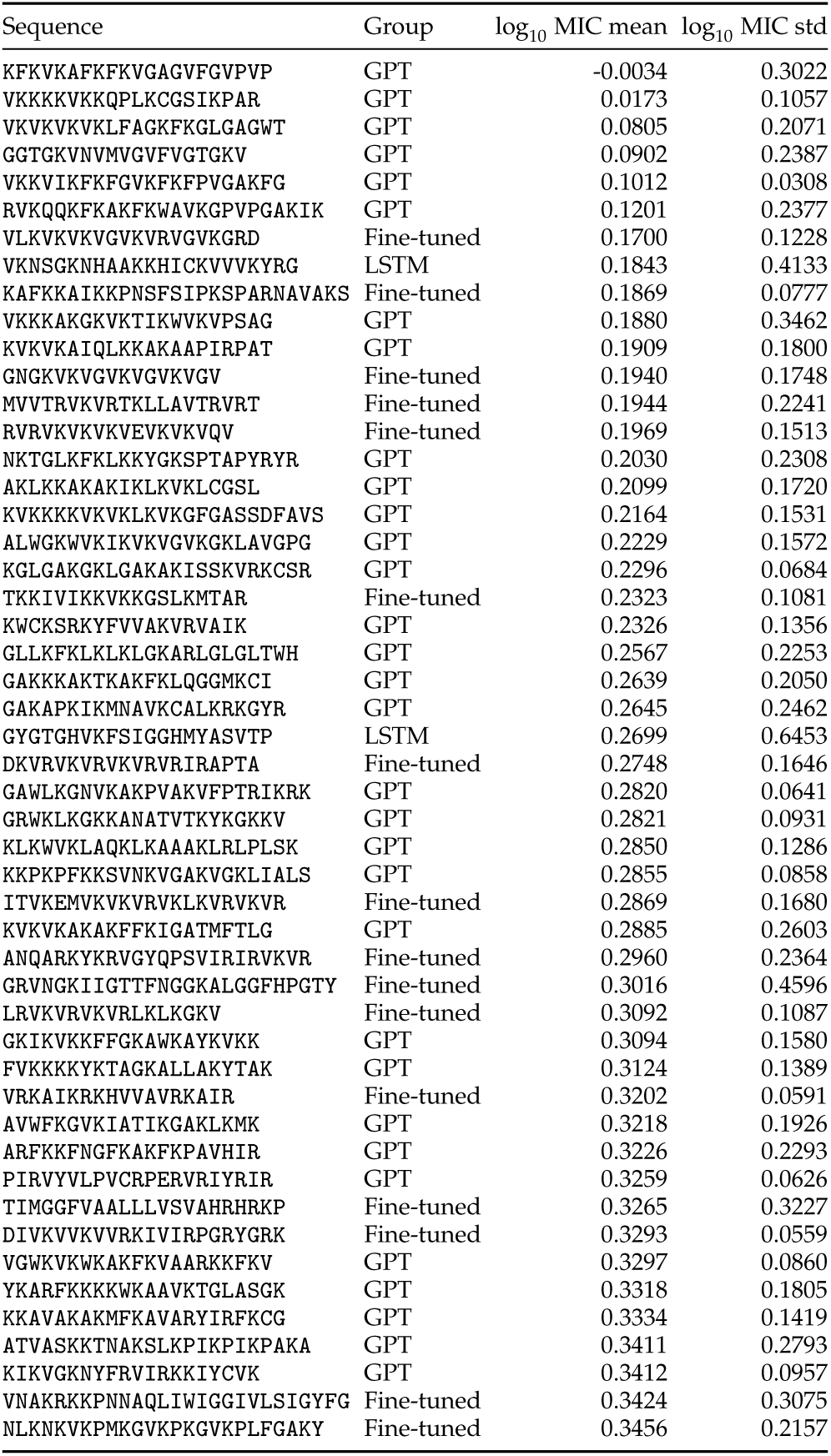
Top 100 Synthesisability-Filtered Peptides (1–50): Sequence, Generative Model, and Ensemble MIC Predictions.

**Table S26.**
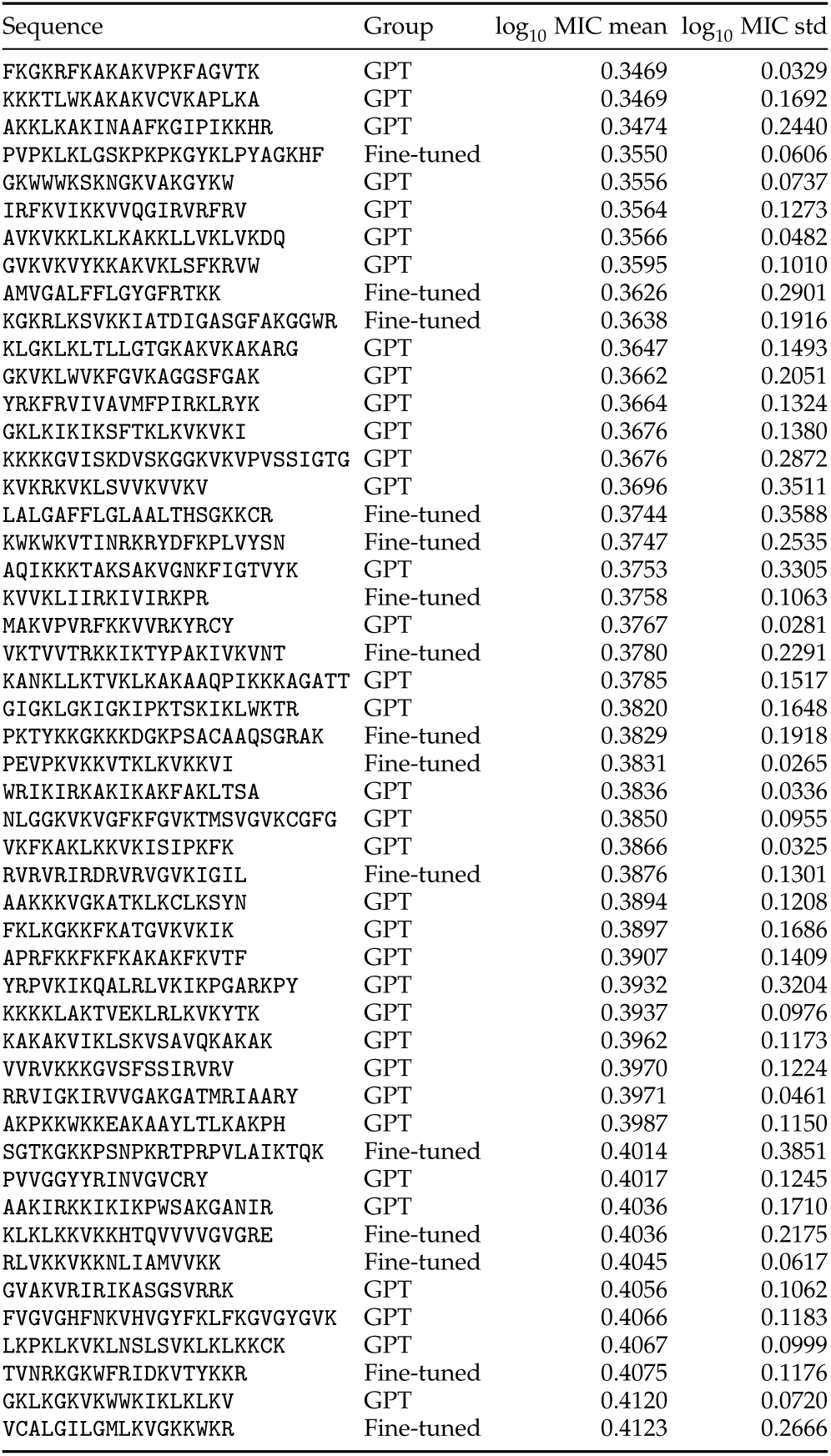
Top 100 Synthesisability-Filtered Peptides (51–100): Sequence, Generative Model, and Ensemble MIC Predictions (continued).

